# Comparison of whole-brain task-modulated functional connectivity methods for fMRI task connectomics

**DOI:** 10.1101/2024.01.22.576622

**Authors:** Ruslan Masharipov, Irina Knyazeva, Alexander Korotkov, Denis Cherednichenko, Maxim Kireev

**Affiliations:** N.P. Bechtereva Institute of the Human Brain, Russian Academy of Sciences, St., Petersburg, Russia

## Abstract

Higher brain functions require flexible integration of information across widely distributed brain regions depending on the task context. Resting-state functional magnetic resonance imaging (fMRI) has provided substantial insight into large-scale intrinsic brain network organisation, yet the principles of rapid context-dependent reconfiguration of that intrinsic network organisation are much less understood. A major challenge for task connectome mapping is the absence of a gold standard for deriving whole-brain task-modulated functional connectivity matrices. Here, we perform biophysically realistic simulations to control the ground-truth task-modulated functional connectivity over a wide range of experimental settings. We reveal the best-performing methods for different types of task designs and their fundamental limitations. Importantly, we demonstrate that rapid (100 ms) modulations of oscillatory neuronal synchronisation can be recovered from sluggish haemodynamic fluctuations even at typically low fMRI temporal resolution (2 s). Finally, we provide practical recommendations on task design and statistical analysis to foster task connectome mapping.

## Introduction

Building a comprehensive map of human brain connections, called the human connectome^1^, can be considered one of the largest and most challenging scientific projects in the field of human neuroscience over the past two decades. In the past decade, the trend in fMRI studies has shifted from functional segregation to functional integration, which is reflected in an increasing number of publications on functional connectivity (FC) compared to task activations (Fig. 1a). Despite the initial scepticism about resting-state FC (RSFC)^2,3^, it has become the most popular fMRI approach and has profoundly advanced our understanding of the intrinsic functional organisation of the brain in health^4,5^ and disease^6,7^. Furthermore, there is a growing awareness of the importance of whole-brain FC dynamics modulated extrinsically by various task demands called the *task connectome*^8^.

**Fig. 1.**
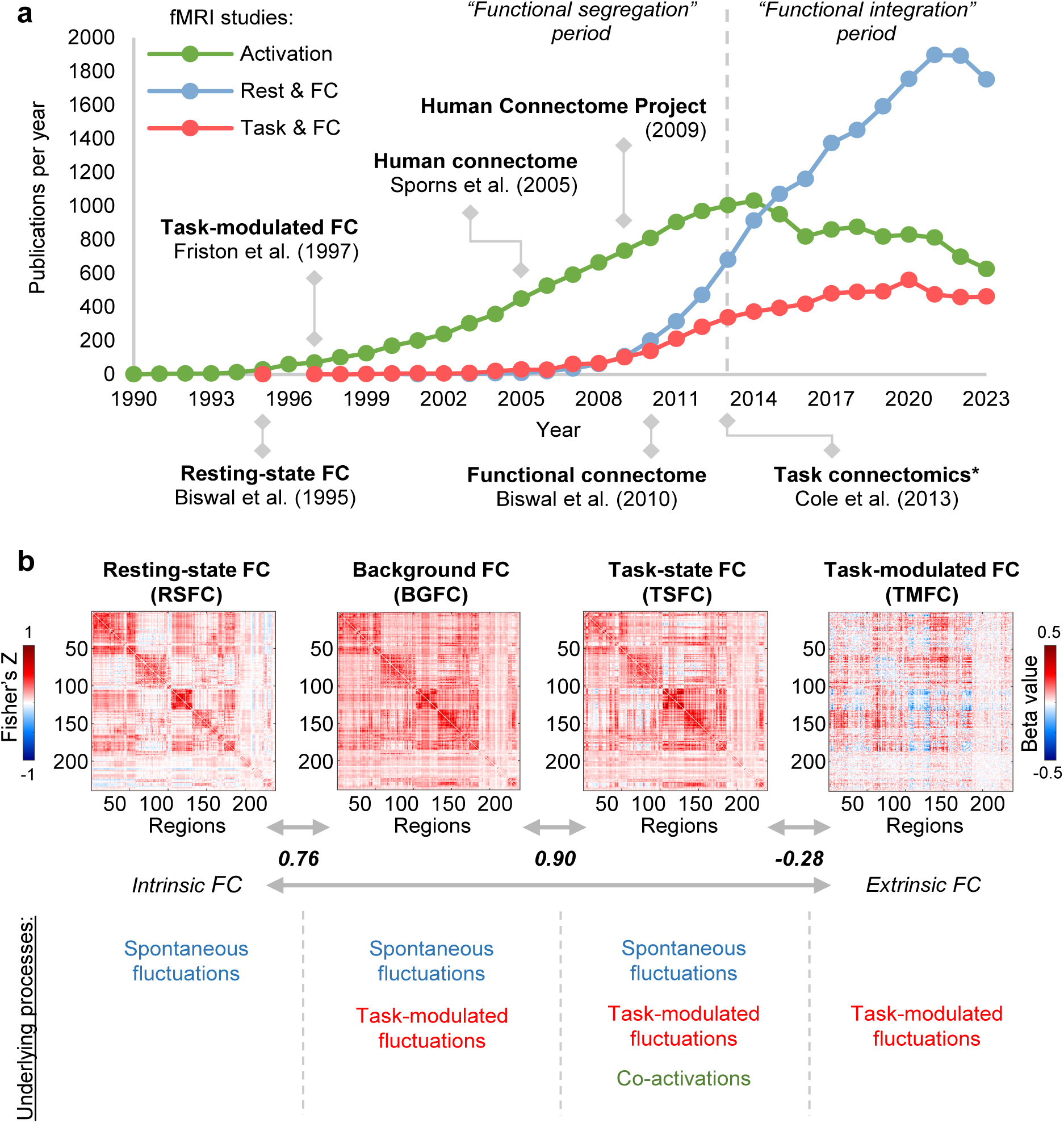
Overview of FC types. (**a**) fMRI publications per year mentioning “activation” (green), “functional connectivity” and “rest” (red), “functional connectivity” and “task” (blue). Data obtained via PubMed search from 1990 to 2023. Here, we note the trend shift in the fMRI field from studies of functional segregation (activation studies) to functional integration (connectivity studies) since 2013. We also observe the dominance of RSFC studies since 2015. (*) – The term “task connectomics” was introduced first introduced by Di et al. (2017)^20^, however, the first large-scale task connectomics study to analyze whole-brain TMFC across multiple tasks was conducted by Cole et al. (2013)^21^. (**b**) Illustration of various types of FC matrices along the intrinsic-extrinsic axis. To calculate FC matrices, we used resting-state and working memory task data from the Human Connectome Project. To estimate TMFC, we applied the gPPI method with the deconvolution procedure (“2-back > 0-back” contrast). To evaluate the similarity between matrices, we used Pearson’s r correlation. The color scales were adjusted for each matrix based on the maximum absolute value and were assured to be positive and negative symmetrical.

Different FC types can be assessed from the task-state blood oxygen-level-dependent (BOLD) signal, ranging from more intrinsic to more extrinsic FC (Fig. 1b). The simplest approach is to correlate the whole time series, similar to the RSFC calculation. We will refer to this type of FC as task-state FC (TSFC), although the terms “task-based FC” and “task FC” are also found in the literature (see Supplementary Table S1 for a review of terms used in previous studies to refer to different FC types). Three neuronal sources of variability underlie TSFC: spontaneous task-independent (intrinsic) fluctuations, task-modulated (extrinsic) fluctuations, and co-activations caused by simultaneous activations without communication between brain regions. As co-activations can spuriously increase FC estimates, it has been proposed to regress out task activations from the task-state BOLD signal and correlate the residuals, which is referred to as background FC (BGFC)^9^. TSFC and BGFC are typically very similar to RSFC since all are driven mainly by spontaneous fluctuations^10^. In contrast, task-modulated FC (TMFC) reflects dynamic changes in FC during one condition compared to another, eliminating the influence of spontaneous task-independent fluctuations and co-activations. Several TMFC methods have been proposed, including direct correlation difference (CorrDiff)^11^, standard^12^, generalised^13^ and correlational^14^ forms of psychophysiological interaction (sPPI, gPPI, cPPI) with and without deconvolution procedure^15^, and beta-series correlations (BSC) based on least-squares all (LSA)^16^, least-squares separate (LSS)^17^, and fractional ridge regression (FRR)^18,19^ procedures. Each TMFC approach has its own advantages and limitations. The lack of a gold standard for deriving whole-brain TMFC matrices and limited knowledge about the fundamental limitations of TMFC estimation related to sluggishness of the BOLD signal hinder the process of task connectome mapping.

Previous simulation studies evaluating TMFC methods have two main limitations. First, they often ignore the dynamic, oscillatory nature of neuronal population activity arising from interactions between excitatory and inhibitory neurons by using simple delta or boxcar functions to simulate neuronal activity in a pair of regions^13,15,22,23^. Second, studies with biophysically realistic simulations based on neural mass models consider only block designs and a limited range of TMFC methods^24,25^. Meanwhile, one of the key unresolved questions is the performance of different TMFC methods for fast event-related designs with slow data acquisition, which are typical for the majority of fMRI studies. Thus, to date, there are no biophysically realistic simulation studies systematically comparing existing TMFC methods in different experimental fMRI setups (see Supplementary Table S2 for an overview).

Here, using biophysically realistic simulations and empirical data, we determined the best-performing whole-brain TMFC methods for different fMRI task designs and identified their limitations. To simulate the neuronal dynamics of multiple interconnected brain regions, we applied a large-scale Wilson-Cowan neural mass model consisting of 100 excitatory-inhibitory units^26,27^. The BOLD signals were generated from simulated neuronal activity using the Balloon-Windkessel haemodynamic model^28^. To control the ground-truth TMFC, we manipulated synaptic weights between neural mass units depending on the task context, which corresponded to short-term plasticity^29,30,31,32^.

First, we demonstrate that one popular TMFC method, cPPI, is not capable of estimating TMFC. Second, we establish that TMFC methods are susceptible to spurious inflation of FC due to co-activations in event-related designs even more than in block designs^25^. Third, we show that the most sensitive methods for rapid event-related designs and block designs are sPPI and gPPI with a deconvolution procedure, while for all other designs, the best method is BSC-LSS. Despite the scepticism regarding the deconvolution procedure^33^, we demonstrate that deconvolution prominently increases the sensitivity of the PPI methods in both event-related and block designs. Fourth, haemodynamic response function (HRF) variability across brain regions and subjects markedly reduces the sensitivity of all TMFC methods. The BSC-LSS method is the most robust to HRF variability. Fifth, while some authors classify PPI as a FC method^33,34^ and others see PPI as a simple regression model of effective connectivity^35,36,37,38,39,40^, our results explicitly demonstrate that PPI with deconvolution can in principle provide information about the direction of causal influence at high signal-to-noise ratios (SNRs), long scan times, and canonical HRF shape. However, in most studies, the asymmetry of PPI matrices is likely to spuriously arise from a low SNR, short event duration, small number of events per condition, small sample size, and long repetition time (TR).

Finally, we demonstrate that rapid (100 ms) task-related modulation of gamma-band neuronal synchronisation can be uncovered from ultra-slow BOLD-signal fluctuations, even with typically slow data acquisition (TR = 2 s). Meanwhile, recently developed fast fMRI sequences (TR < 1 s) yield increased sensitivity of TMFC methods not only by increasing the amount of data but also due to more precise insights into fast neuronal dynamics hidden behind the sluggish haemodynamic processes.

## Results

To compare different TMFC methods, we first used empirical fMRI data from the Human Connectome Project (HCP)^41^ and the Consortium for Neuropsychiatric Phenomics (CNP)^42^. In particular, we considered two block design tasks (working memory and social cognition tasks, N = 100) from the HCP dataset^41,43^, two event-related tasks (stop-signal and task-switching tasks, N = 115) from the CNP dataset^42,44^ and resting-state data from both datasets. To construct empirical FC matrices, we used a set of functionally defined regions of interest (ROIs) covering the whole brain^45^. The ROIs were defined as spheres with a radius of 4 or 5 mm. We discarded ROIs for which data were incomplete for at least one subject. As a result, we utilised 239 ROIs for the HCP dataset and 246 ROIs for the CNP dataset.

Next, we performed a series of simulations to compare the sensitivity and specificity of TMFC methods in experiments with block and event-related designs, different SNRs, sample sizes, duration of events, mean interstimulus intervals (ISIs) and number of events. For block designs, we considered the CorrDiff approach and PPI methods, and for event-related designs, we considered the PPI methods and BSC methods. We also compared the effectiveness of fast data acquisition for a fixed number of scans and fixed total scan time. Then, we assessed the impact of HRF variability on the sensitivity of TMFC methods. Finally, we performed simulations with symmetric (undirected) and asymmetric (directed) ground-truth synaptic matrices to identify sources of PPI matrix asymmetry.

### Empirical comparison of TMFC methods

We found that all TMFC methods, except cPPI, produce similar unthresholded matrices for the block design (working memory task, Fig. 2a). The correlations between CorrDiff matrices and symmetrised sPPI and gPPI matrices ranged from 0.66 to 1. We also found that the correlations between these TMFC matrices and RSFC, BGFC, TSFC and cPPI matrices ranged from -0.13 to -0.28.

**Fig. 2.**
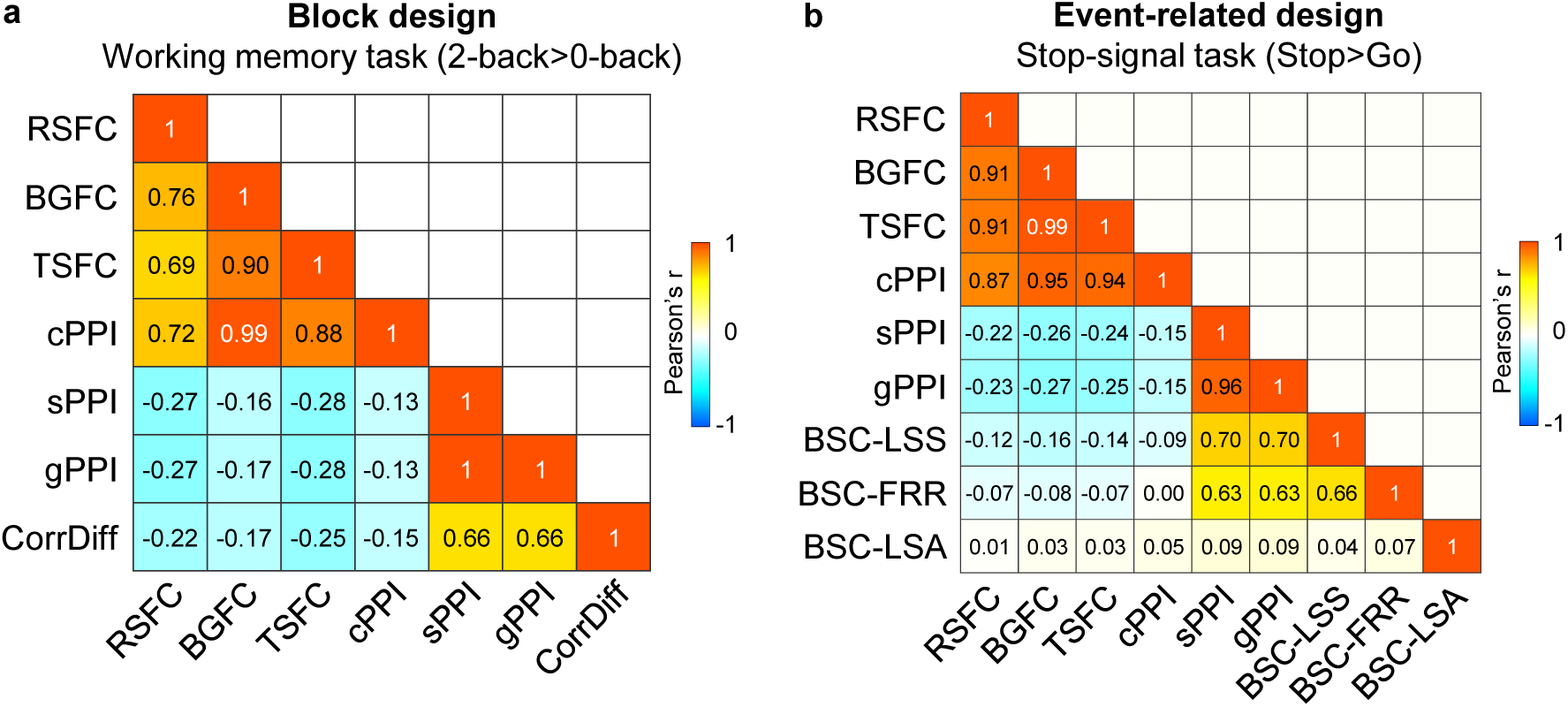
Correlations between unthresholded RSFC, BGFC and TSFC matrices and FC matrices obtained by different TMFC methods. To evaluate the similarity between the raw FC matrices, we calculated Pearson’s r correlations between lower diagonal elements. sPPI and gPPI matrices were symmetrised. All PPI terms were calculated with the deconvolution step. **(a)** Results for the block design: working memory task. **(b)** Results for the event-related design: stop-signal task.

For the event-related design (stop-signal task), similar TMFC matrices were produced by the sPPI, gPPI, BSC-LSS and BSC-FRR methods, with correlation coefficients ranging from 0.63 to 0.96 (Fig. 2b). At the same time, correlations between these TMFC matrices and the RSFC, BGFC, TSFC and cPPI matrices ranged from -0.09 to 0.004. Remarkably, the BSC-LSA approach produced a random-like matrix correlated with all other FC matrices with a correlation of 0.01–0.09. The similarities and differences between these FC methods are also confirmed by calculating the overlap between thresholded matrices (Supplementary Fig. S1 and S2). Analogous results were obtained for empirical data from other tasks with block and event-related designs (social cognition task and task-switching task), which demonstrates that the observed effects are specific to a design type rather than a particular task (Supplementary Fig. S3 and S4).

Notably, the matrices generated by the cPPI method are similar to matrices obtained using the RSFC, BGFC, and TSFC approaches, which mainly reflect task-independent spontaneous fluctuations^10^. Correlations between the cPPI, RSFC, BGFC and TSFC matrices ranged from 0.69 to 0.99 for block and event-related design (Fig. 2a, b). The partial correlation between two PPI terms controlling for physiological (BOLD signal in both regions) and psychological variables (task regressor), as implemented in cPPI, was as high as the simple correlation between these two PPI terms and the correlation between physiological regressors (Fig. 3). Therefore, we concluded that the cPPI approach is unable to separate task-modulated (extrinsic) from task-independent (intrinsic) sources of FC. However, comparison of FC matrices based solely on empirical data does not allow us to say which method better reflects the true TMFC. To answer this question, we applied a biophysically realistic simulation approach that enabled us to control the ground-truth TMFC.

**Fig. 3.**
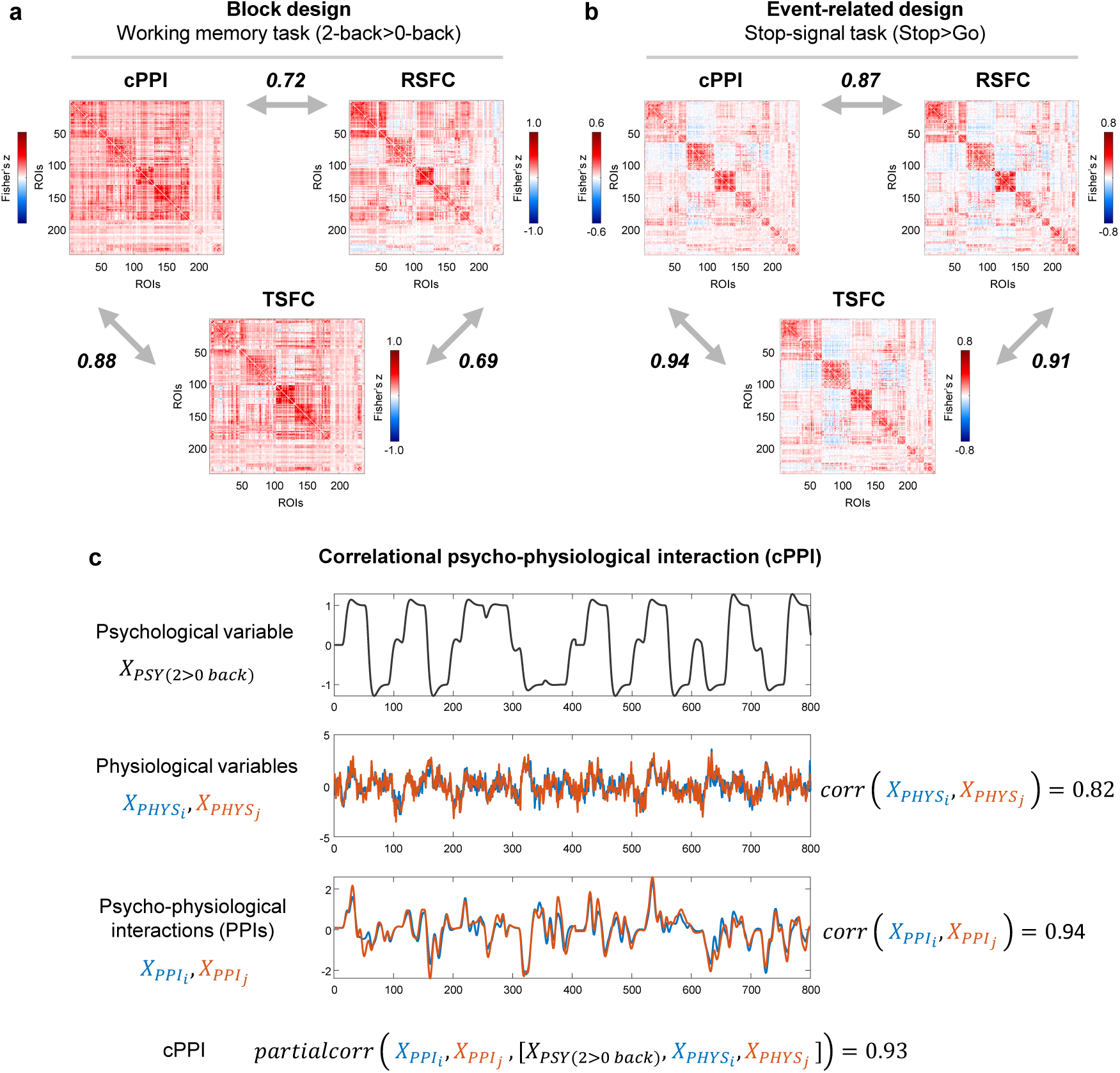
The cPPI approach produces FC matrices similar to the RSFC and TSFC matrices. (a) Correlations for the block design task (working memory task) and resting-state data from the HCP dataset. (b) Correlations for the event-related design task (stop-signal task) and resting-state data from the CNP dataset. (c) Illustration of the cPPI approach. The working memory task time series were taken from the two ROIs as an example. All PPI terms were calculated with the deconvolution step. To evaluate the similarity between matrices, we used Pearson’s r correlation. The color scales were adjusted for each matrix based on the maximum absolute value and were assured to be positive and negative symmetrical.

### Large-scale neural mass simulations

Our simulation approach was based on the coupled oscillator model for FC proposed by Mateo et. al (2017)^46^. Using optogenetic manipulations and concurrently measuring local field potential, arteriole diameter and blood oxygenation in wake mice, they showed that correlations between ultra-slow BOLD fluctuations (i.e., FC measured by fMRI) are caused by synchronised ultra-slow fluctuations in arteriole diameter, which in turn are caused by ultra-slow modulation of the envelopes of synchronised gamma-band oscillations (Fig. 4a). Accordingly, we modelled gamma-band activity of 100 interconnected regions using the large-scale Wilson-Cowan model, and also modelled arteriole diameter dilations and blood oxygenation change using the Balloon-Windkessel model (Fig. 4b). For details about the simulation procedures, see Methods. The matrices of gamma-band neuronal synchronisation measured by the phase-locking value were closely matched to synaptic weight matrices (Fig. 4c, d). In accordance with previous empirical observations by Mateo et. al (2017)^46^, we observed strong correlations between simulated ultra-slow fluctuations of the gamma-band envelope and time-shifted BOLD signal. For the event-related design, the simulated BOLD signal was correlated with the gamma-band envelope at r = 0.79 with a 3.5 s time lag (Fig. 4e-j). Similar results were obtained for the block design, where the correlation was r = 0.81 with a 3.5 s time lag (Supplementary Fig. S5).

**Fig. 4.**
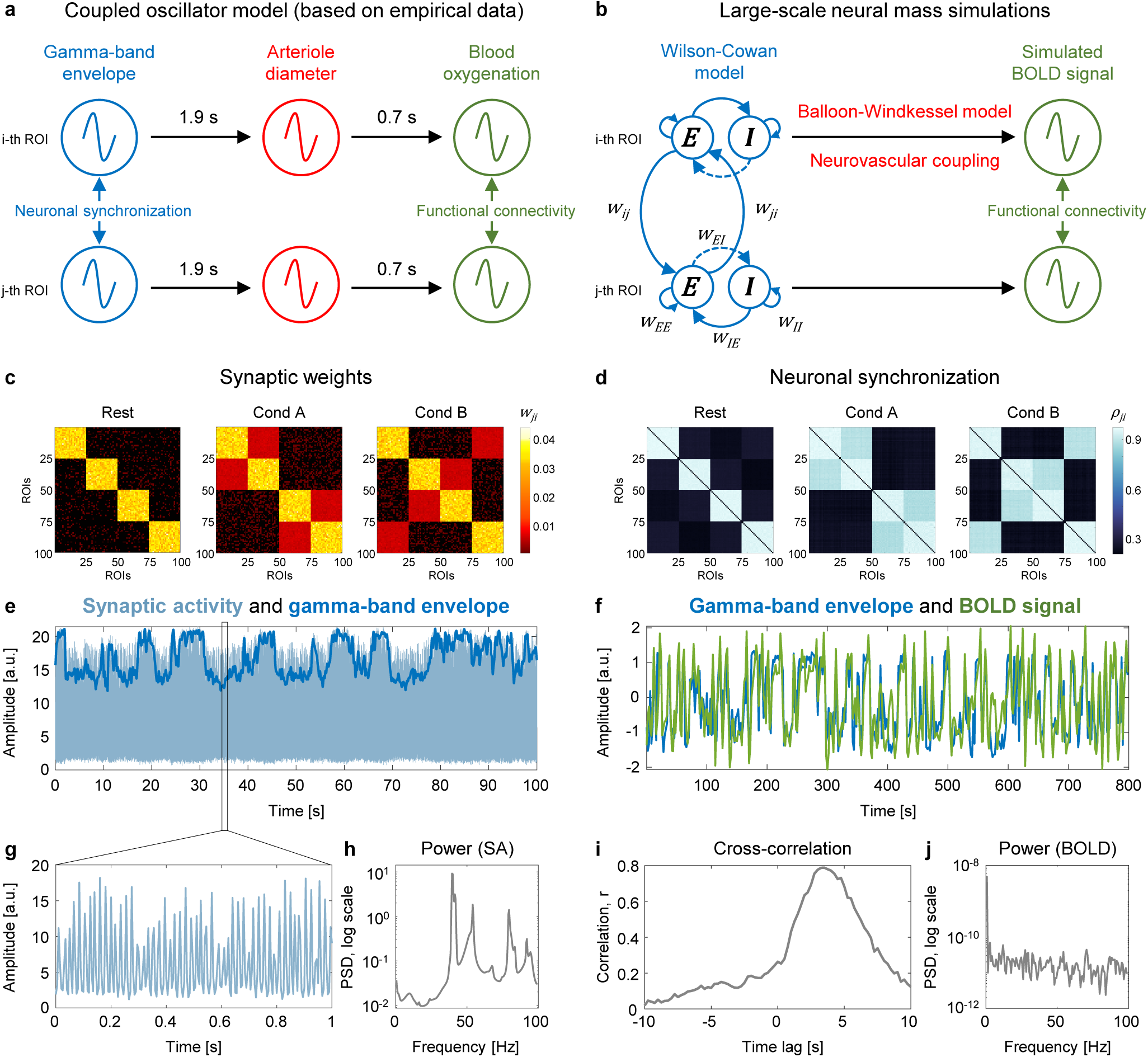
The emergence of ultra-slow BOLD signal fluctuations from ultra-slow modulations of fast oscillatory activity. **(a)** Coupled oscillator model for functional connectivity proposed by Mateo et al.^46^ Our simulation approach was based on the empirical evidence^46^ that correlations between ultra-slow BOLD fluctuations are caused by ultra-slow modulation of the envelopes of synchronised gamma-band oscillations. **(b)** Scheme for generating the BOLD signal from gamma-band oscillations simulated by the Wilson-Cowan model. **(c)** Ground-truth symmetric synaptic weight (*w_ji_*) matrices for a single subject. For the long-range connections between Wilson-Cowan units (*w_ji_*), we used three synaptic weight matrices corresponding to two task conditions (“Cond A” and “Cond B”) and interim “Rest” periods. Each synaptic weight matrix consisted of 100 brain regions, which corresponded to the minimum number of areas in brain atlases^47^, and four functional modules, given the presence of such modules in the human brain^5^. The highest synaptic weights were within each module during the “Rest” periods^5^. Under the task conditions, we increased synaptic weights between functional modules relative to the “Rest” periods^8^. **(d)** Gamma-band neuronal synchronisation estimated by the phase-locking value (*ρ_ji_*) for a single subject. The simulation was performed for the event-related design with one hundred 1 s events per condition and mean ISI = 6 s. **(e)** Example of simulated synaptic activity (SA) and gamma-band envelope for one of 100 connected brain regions (ROIs). **(f)** The time series of the gamma-band envelope and BOLD signal generated by the Balloon-Windkessel model based on simulated SA. We used the standard parameters of the haemodynamic model^28^, which have previously been used in whole-brain RSFC simulation studies^27,48^. **(g)** One second of simulated SA. **(h)** Power spectral density (PSD) of simulated SA. The main peak at 40 Hz (gamma-band oscillations). **(i)** Cross-correlation between the gamma-band envelope and BOLD signal. The maximum correlation *r* = 0.79 corresponds to a time lag of 3.5 seconds. **(j)** Power spectral density of the BOLD signal.

### The cPPI method fails to estimate TMFC

We first considered simulations without co-activations to investigate whether different TMFC methods produce FC matrices similar to ground-truth synaptic weight matrices for a sample size N=100, SNR = 0.4, and TR = 2 s. For differences between simulated conditions (“Cond A-B”), almost all tested methods produced TMFC matrices similar to the ground truth (Fig. 5a-g). In contrast, the cPPI method produced matrices similar to BGFC and TSFC (Fig. 5h), which replicated the above-mentioned findings from the empirical data analysis (Fig. 2). Knowing the ground-truth FC patterns, we revealed that the cPPI method, unlike other TMFC methods, does not show the modulation of FC between compared task conditions (the “Cond A-B” effect), but rather shows the sum of all FC during both conditions, that is, the “Cond A+B” effect (Supplementary Fig. S6). Since task-unrelated FC is present in both conditions, we see both task-unrelated and task-related effects in the cPPI matrix. Other TMFC methods effectively remove task-unrelated effect by subtracting FC in one condition from another. Thus, the cPPI method does not contrast FC in one condition relative to another and does not remove task-unrelated effects (functional connections that is high at rest). Therefore, we excluded the cPPI method from further analysis, as it is unable to assess TMFC.

**Fig. 5.**
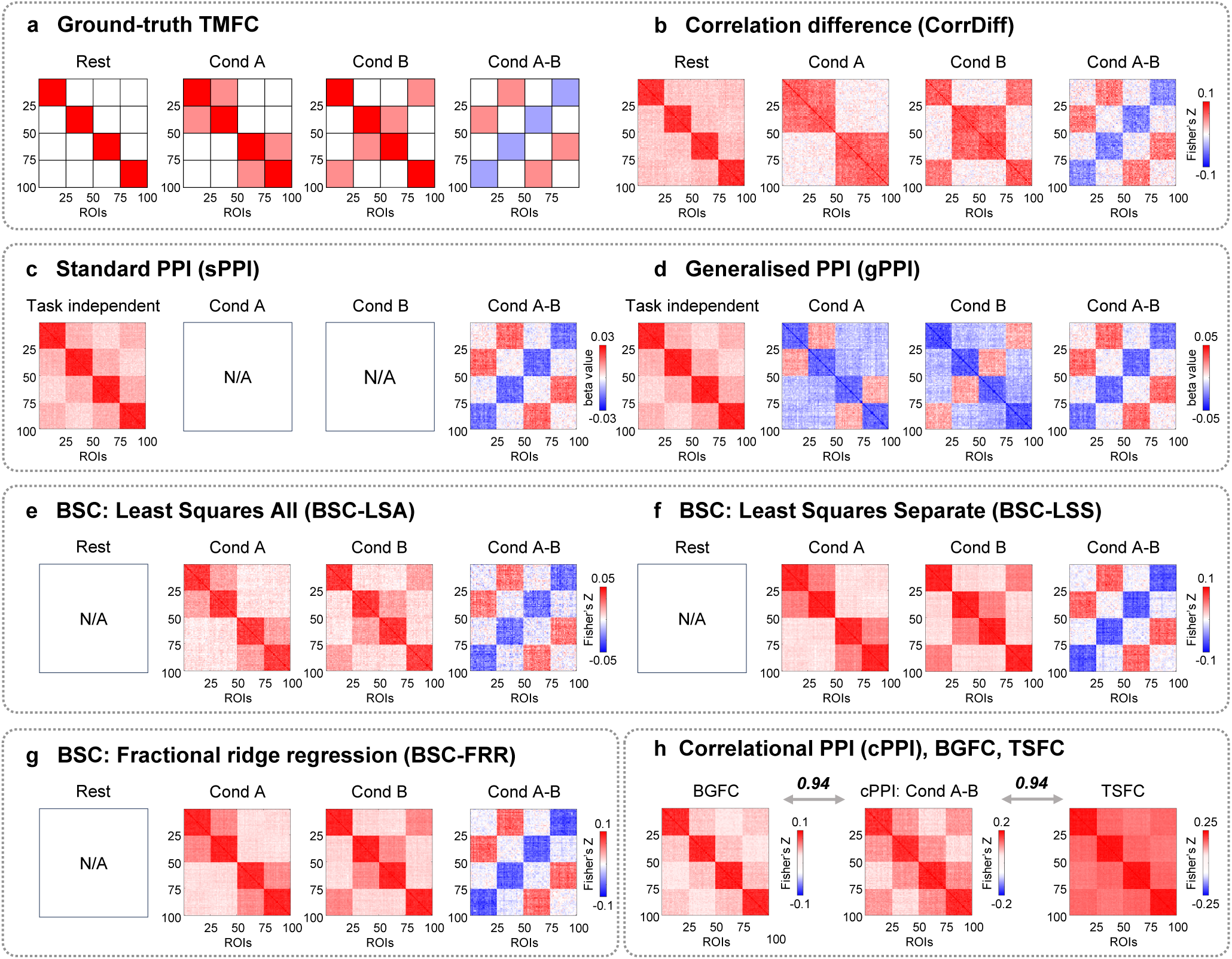
Illustration of FC matrices produced by different TMFC methods based on simulated data. An example of simulation results for sample size N = 100, SNR = 0.4 and TR = 2 s. **(a)** Expected FC matrices based on ground-truth synaptic weight matrices. **(b-d)** For the CorrDiff, sPPI and gPPI methods, we considered the block design with ten 20 s blocks per condition. **(e-g)** For the BSC-LSA, BSC-LSS and BSC-FRR methods, we considered the event-related design with one hundred 1 s events per condition and mean ISI = 6 s. **(h)** The cPPI, BGFC and TSFC matrices were calculated based on the block design simulation. Analogous results were obtained for the event-related design simulation. To evaluate the similarity between matrices, we calculated Pearson’s r correlations between lower diagonal elements. All PPI terms were calculated using the deconvolution step. sPPI and gPPI matrices were symmetrised. The color scales were adjusted for each matrix based on the maximum absolute value and were assured to be positive and negative symmetrical.

Despite its simplicity, the CorrDiff approach allows the computation of easily interpretable FC matrices separately for task and rest blocks; these matrices adequately reflect the underlying ground-truth synaptic weight matrices (Fig. 5a, b). The sPPI and gPPI approaches enable estimation of both TMFC and task-independent FC (Fig. 5c, d). Task-independent FC is based on beta coefficients for physiological regressors delivered from selected ROIs and is similar to FC during rest periods. In addition, the gPPI approach can be used to calculate FC separately for each of the task conditions (i.e., Condition > Baseline). However, the gPPI matrices for each task condition should be interpreted with caution. The sign of the PPI estimates between nodes that exhibit high connectivity during rest periods depends on the deconvolution procedure and mean centering of the psychological regressor prior to PPI term calculation. With deconvolution and mean centering, the PPI estimates between these nodes become negative, deviating from the ground truth, see Fig. 5d (for more details, see Methods and Supplementary Information 6). The BSC approaches enable the calculation of FC matrices for each of the task conditions that are consistent with the ground truth, but do not allow to calculate FC for rest periods (task-independent FC) since rest periods are usually not modelled explicitly (Fig. 5e-g).

### Co-activations spuriously inflate TMFC estimates

Next, we considered simulations with co-activations to investigate how different TMFC methods address artificial inflation of TMFC estimates due to simultaneous activation of brain regions without task-related modulation of synaptic weights between them (Fig. 6a-c). Sensitivity (true positive rate, TPR) and specificity (true negative rate, TNR) were calculated based on TMFC matrices thresholded at α = 0.001 with false discovery rate (FDR) correction (Supplementary Information 1).

**Fig. 6.**
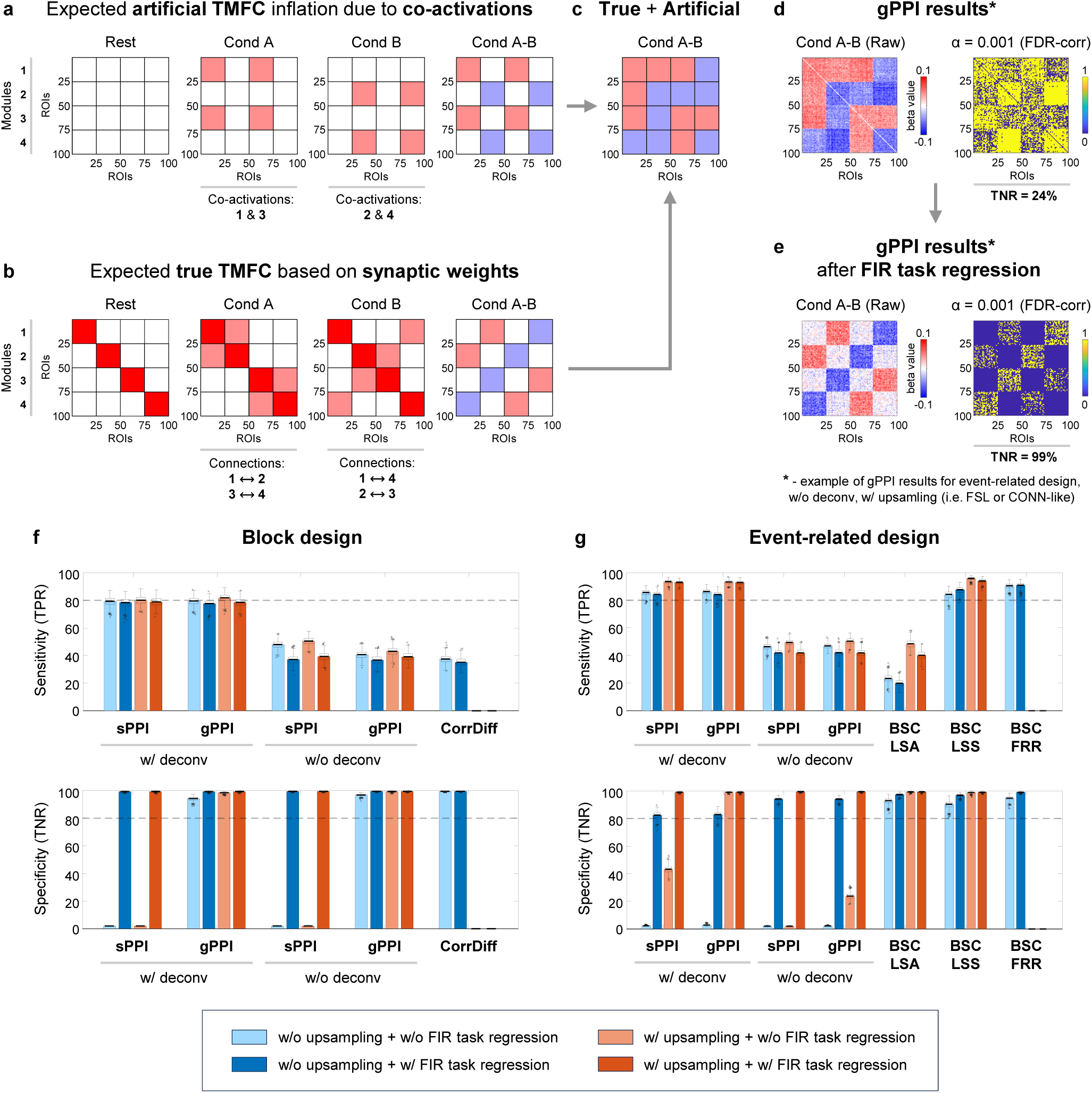
Inflation of TMFC estimates due to co-activations. Simulation results for sample size N = 100, SNR = 0.4 and TR = 2 s. **(a)** Expected influence of co-activations on FC estimates (artificial TMFC). To model co-activations, we added simultaneous haemodynamic responses for different functional modules to the simulated BOLD signal without changing the synaptic weights between them. Functional modules №1 and №3 are co-activated in “Cond A”, while modules №2 and №4 are co-activated in “Cond B”. Artificial FC inflation was expected within and between these modules. **(b)** Expected FC matrices based on ground-truth synaptic weight matrices (true TMFC). In “Cond A”, synaptic weights were increased between modules №1 and №2 and modules №3 and №4. In “Cond B”, synaptic weights were increased between modules №1 and №4 and modules №2 and №3. **(c)** If the TMFC method fails to eliminate co-activations, we will observe FC changes between all ROIs (artificial + true TMFC). **(d-e)** Raw and thresholded TMFC matrices obtained using the gPPI method without (w/o) the deconvolution step and with (w/) design matrix upsampling, similar to the gPPI implementation in the FSL or CONN toolbox. TMFC matrices were thresholded at α = 0.001 (two-sided one-sample t test, false discovery rate (FDR) correction). **(d)** Without task activation regression, we observed FC changes between almost all ROIs (low specificity). **(e)** After finite impulse response (FIR) task regression, artificial TMFC was removed, leaving mostly true TMFC (high specificity). **(f)** Sensitivity (true positive rate, TPR) and specificity (true negative rate, TNR) of different TMFC methods for the block design with ten 20 s blocks per condition. **(g)** TPR and TNR for the event-related design with one hundred 1 s events per condition and mean ISI = 6 s. All TMFC matrices were calculated with and without FIR task regression, as well as with and without upsampling of the design matrix before convolution, except for CorrDiff and BSC-FRR. We did not perform upsampling for the CorrDiff and BSC-FRR methods. All PPI terms were calculated with and without the deconvolution step. sPPI and gPPI matrices were symmetrised. Boxplots whiskers are drawn within the 1.5 interquartile range (IQR), computed from 1000 random resamplings with replacement. The color scales were adjusted for each matrix based on the maximum absolute value and were assured to be positive and negative symmetrical.

As a result, we found that if co-activations are not removed from the fMRI time series before TMFC analysis, they spuriously inflate TMFC estimates in three cases. First, the sPPI method practically does not eliminate co-activations and therefore has near-zero specificity in both block and event-related designs (Fig. 6f-g). Second, the gPPI method demonstrates low specificity when applied to event-related designs without deconvolution (Fig. 6d, g). Third, the specificity of all TMFC methods decreases if the design matrix is not upsampled before convolution with the haemodynamic response function (Fig. 6f,g).

Upsampling of the design matrix is used to improve the convolution procedure and is implemented in many popular neuroimaging software packages (SPM, FSL, AFNI, CONN toolbox). However, it may be absent in some in-house TMFC analysis scripts. The most prominent effect of upsampling on specificity can be seen for the gPPI method with deconvolution in event-related designs (Fig. 6g).

To better isolate TMFC from co-activation effects, it has been proposed to regress out task activations using finite impulse response (FIR) functions prior to TMFC analysis^21^. FIR task regression substantially improved the specificity of all TMFC methods in both block and event-related designs (Fig. 6f, g). For instance, the gPPI method without deconvolution had a specificity of 24% before FIR task regression and 99% after (Fig. 6e). The downside of FIR task regression is that it slightly reduces sensitivity, most notably for the sPPI and gPPI methods without deconvolution and the BSC-LSA method.

In all subsequent sections, we consider simulations with co-activations and perform TMFC analyses with FIR task regression and design matrix upsampling, unless otherwise stated. We did not perform upsampling for CorrDiff and BSC-FRR, since the CorrDiff method does not rely on general linear models, and FRR implementation in the GLMsingle toolbox requires the task design matrix to have the same temporal resolution as the data to be convolved^19^. We will not report specificity further since in no case did it fall below 95%.

### Influence of noise and sample size on the sensitivity of TMFC methods

In this section, we describe the robustness of TMFC methods to high noise levels and low sample sizes. For the block design, the PPI methods with deconvolution were the most sensitive and robust to noise (Fig. 7a, Supplementary Table S3). The least sensitive method was CorrDiff since it needed a sample size > 100 to achieve > 80% sensitivity at high SNR = 0.5, while the PPI methods with deconvolution needed a sample size > 50 (Fig. 7c).

**Fig. 7.**
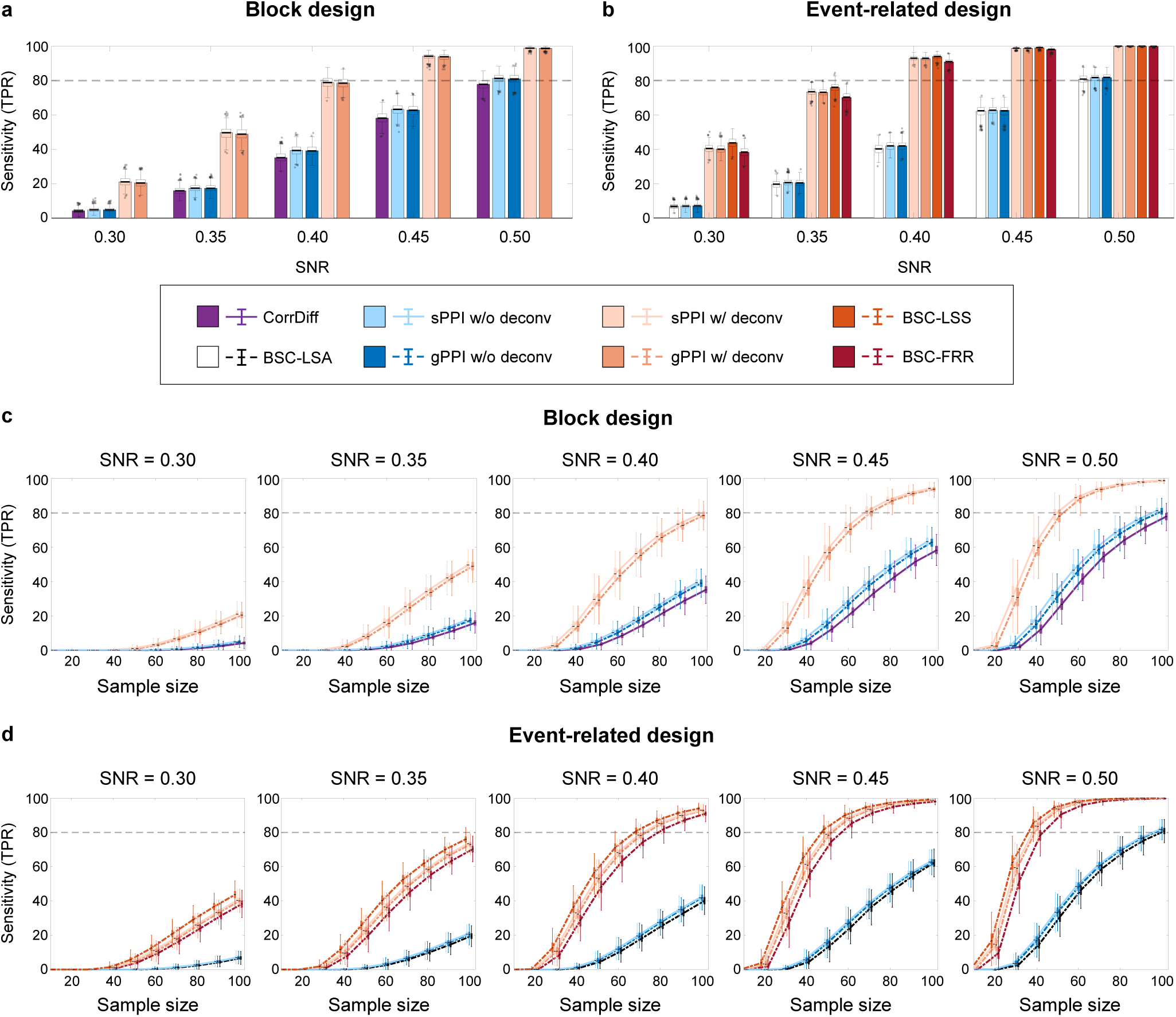
Sensitivity of all TMFC methods depending on the signal-to-noise ratio (SNR) and sample size. Simulation results for TR = 2 s. **(a-b)** Sensitivity of different TMFC methods at sample size N = 100 for **(a)** the block design with ten 20 s blocks per condition and **(b)** event-related design with one hundred 1 s events per condition and mean ISI = 6 s. **(c-d)** Sensitivity of different TMFC methods depending on the sample size for **(c)** the block design and **(d)** event-related design. PPI terms were calculated with (w/) and without (w/o) the deconvolution step. sPPI and gPPI matrices were symmetrised. Boxplots whiskers are drawn within the 1.5 interquartile range (IQR), computed from 1000 random resamplings with replacement.

For the event-related design, the BSC-LSS method was the most sensitive (Fig. 7b, Supplementary Table S4). The difference between the BSC-LSS method, the BSC-FRR method and PPI methods with deconvolution was relatively small and more pronounced at high noise levels (SNR < 0.40). At the same time, the BSC-LSA method had the lowest sensitivity due to the multicollinearity problem^17^. The BSC-LSS and BSC-FRR methods, as well as the PPI methods with deconvolution, needed a sample size N > 50 to achieve > 80% sensitivity at SNR = 0.5 (for the event-related design), while the BSC-LSA method needed a sample size N > 100 (Fig. 7d).

In contrast to previous simulation studies^13,22^, we did not detect a noticeable increase in the sensitivity of the gPPI method compared to the sPPI method. Additional Bayesian analysis provided evidence for the absence of differences between these methods for the block and event-related designs (Supplementary Tables S3 and S4).

It has also been previously suggested that deconvolution may benefit only event-related designs and can be omitted for block designs^8^. Indeed, some popular neuroimaging packages implement the PPI method without the deconvolution step (e.g., FSL and CONN toolbox). Here, we show that the deconvolution procedure substantially increases the sensitivity of the PPI methods in both block and event-related designs (Fig. 7a, b). Without deconvolution, these methods failed to achieve > 80% sensitivity for sample sizes N < 100 and SNR = 0.5 (Fig. 7c, d). Therefore, in the remaining sections, we will consider the sPPI and gPPI methods only with deconvolution, unless otherwise stated.

### Sensitivity of the TMFC methods for event-related designs with different timing parameters

Next, we independently varied the temporal parameters of the event-related design, including event duration, mean ISI, number of events and data acquisition (TR), to assess their impact on the sensitivity of TMFC methods with a sample size N =100 and medium SNR = 0.40. Shortening the event duration substantially decreased the sensitivity of all TMFC methods (Fig. 8a, Supplementary Table S5). The BSC-LSS method was slightly more sensitive than the BSC-FRR method and the PPI methods, which was more noticeable for short event durations < 1 s; in contrast, the BSC-LSA method achieved > 80% sensitivity only for long event durations > 2 s.

**Fig. 8.**
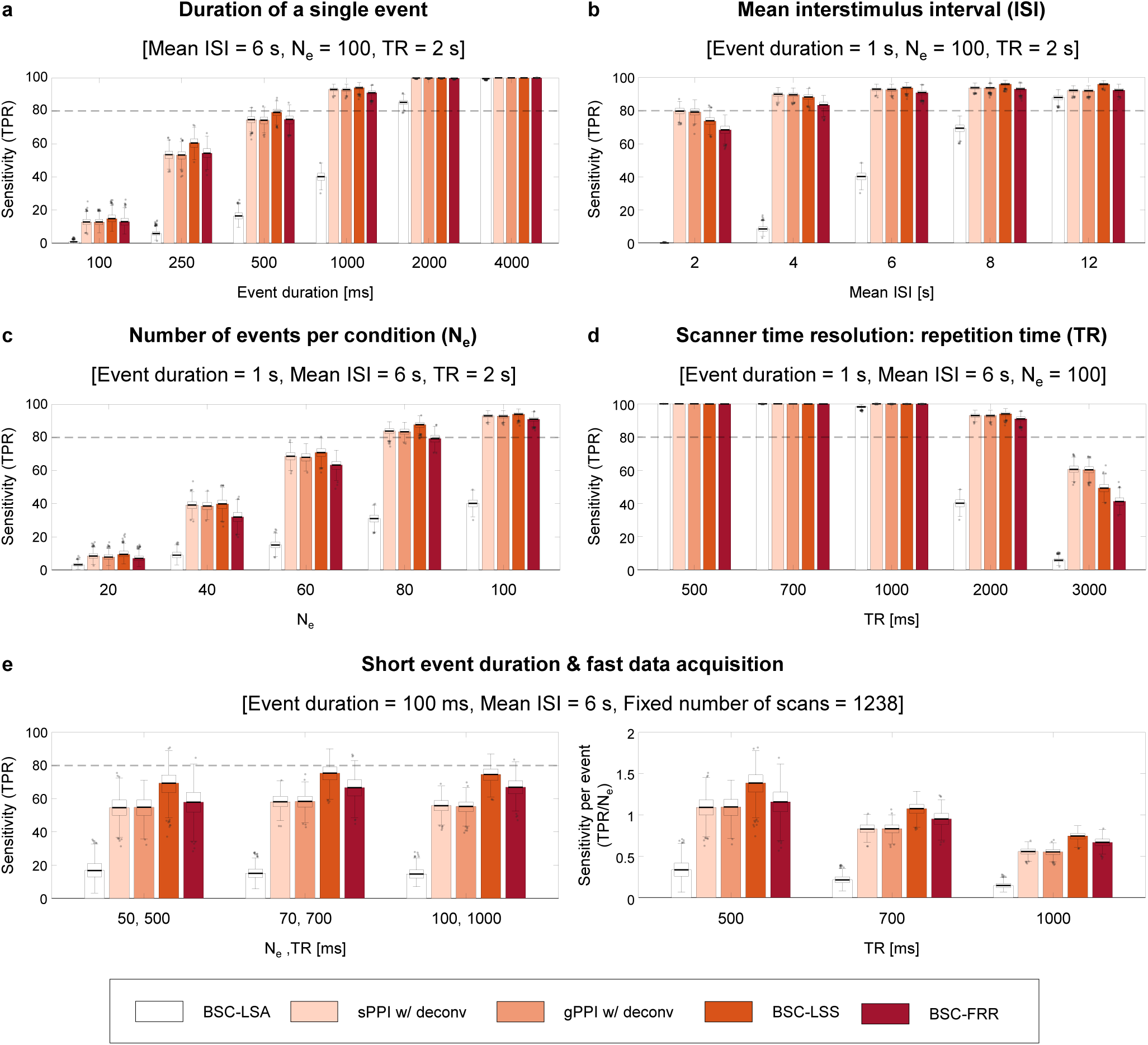
Sensitivity of TMFC methods for event-related designs with different temporal parameters. Simulation results for sample size N = 100 and SNR = 0.4. The default event-related design consisted of one hundred 1 s events per condition and mean ISI = 6 s. **(a)** Sensitivity depending on the duration of a single event. **(b)** Sensitivity depending on the mean ISI. **(c)** Sensitivity depending on the number of events per condition (N_e_). **(d)** Sensitivity depending on the repetition time (TR) for a fixed scan time (23.6 minutes). **(e)** Sensitivity for the event-related design with short event duration (100 ms), fast data acquisition (≤ 1000 ms) and fixed number of scans (1238). Fixing the number of scans resulted in variable scan times and numbers of events. The panel on the left side represents raw sensitivity (true positive rate, TPR). The panel on the right side represents normalised sensitivity per single event (TPR/N_e_). sPPI and gPPI matrices were symmetrised. Boxplots whiskers are drawn within the 1.5 interquartile range (IQR), computed from 1000 random resamplings with replacement.

Shortening the mean ISI slightly decreased the sensitivity of the BSC-LSS, BSC-FRR and PPI methods and substantially reduced the sensitivity of the BSC-LSA method (Fig. 8b, Supplementary Table S6). The BSC-LSA method had > 80% sensitivity only for slow event-related designs with mean ISI = 12 s. For the rapid event-related designs (ISI ≤ 4 s), the PPI methods were slightly more sensitive than the BSC-LSS and BSC-FRR approaches. At longer ISI (≥ 6 s), the BSC-LSS approach was more sensitive than the BSC-FRR and PPI methods.

Reducing the number of events also substantially decreased the sensitivity of all TMFC methods (Fig. 8c, Supplementary Table S7). The BSC-LSS approach was slightly more robust to shortening scan time than other methods. The BSC-LSS, BSC-FRR and PPI methods needed at least 80 events per condition to achieve > 80% sensitivity.

Finally, fast data acquisition (TR < 1 s) yielded the maximum sensitivity for all methods (Fig. 8d, Supplementary Table S8), while reducing the fMRI temporal resolution from the typical TR = 2 s to the frequently used TR = 3 s reduced sensitivity below 80%. Notably, the PPI methods were more sensitive than the BSC-LSS and BSC-FRR methods for TR = 3 s.

We also considered TMFC simulations with task designs parameters derived from empirical HCP and CNP tasks. For the working memory task with 8 blocks per condition, block duration = 27.5 s, interleaved by 15 s rest blocks, TR = 0.72 s, total scan time ≈ 10 min (810 dynamics), SNR = 0.4, and sample size N = 100, the sensitivity of the CorrDiff and PPI methods with deconvolution was 61% and 99%, respectively. For the social cognition task with 5 blocks per condition, block duration = 23 s, interleaved by 15 s rest blocks, TR = 0.72 s, total scan time ≈ 7 min (548 dynamics), SNR = 0.4, and sample size N = 100, the sensitivity of the CorrDiff and PPI methods with deconvolution was 23% and 97%, respectively. For the stop-signal task with 96 “Go” and 32 “Stop” events, mean ISI = 1 s (ranged from 0.5 to 4 s), event duration = 1.5 s, TR = 2 s, total scan time ≈ 7 min (184 dynamics), SNR = 0.4, and sample size N = 115, the sensitivity of the BSC-LSS and PPI methods with deconvolution was 3% and 15%, respectively. The sensitivity of BSC-LSS was lower than that of PPI methods due to the very short ISI. When we halved the number of “Stop” events (i.e., only considered “Correct Stop” events), the sensitivity dropped to 0% and 8%, respectively. For the task-switching task with 24 “Switch” and 72 “No Switch” events, mean ISI = 3 s, event duration = 1 s, TR = 2 s, total scan time ≈ 6 min (208 dynamics), SNR = 0.4, and sample size N = 115, the sensitivity of the BSC-LSS and PPI methods with deconvolution was 13% and 15%, respectively.

Therefore, block design tasks from the HCP dataset are well suited for TMFC estimation. Although the total duration of these tasks is relatively short, it is compensated by short TR. At the same time, event-related tasks from the CNP dataset may not have sufficient sensitivity to capture whole-brain TMFC. In particular, these tasks are unbalanced and have very few events of interest (32 “Stop” and 24 “Switch” events).

### Rapid synchronisation can be revealed even with typically slow fMRI data acquisition

One intriguing question is the principal ability of BOLD fMRI to assess rapid modulations of gamma-band neuronal synchronisation evoked by short events given typically slow data acquisition (TR = 2 s). In the previous section, we showed that the most sensitive TMFC method (BSC-LSS) revealed a fairly small number of true positives (TPR = 15%) for the event-related design with one hundred 100 ms events per condition (Fig. 8a). When we doubled the number of events, the sensitivity was increased to TPR = 57%. Therefore, typically slow fMRI sequences can in principle detect task-related neuronal synchronisation on the order of 100 ms. However, the sensitivity to such rapid synchronisation with slow data acquisition is relatively weak, and many events are required to detect them.

### Importance of fast fMRI data acquisition for TMFC analysis

Another possible way to increase the sensitivity of TMFC methods to rapid neuronal synchronisation is to employ fast fMRI data acquisition techniques. Increasing the fMRI temporal resolution from TR = 2 s to TR = 500 ms resulted in 99% sensitivity for the event-related design with one hundred 100 ms events per condition when the BSC-LSS method was applied. At a fixed task duration (scan time = 23.6 minutes), decreasing TR from 2 s to 500 ms increased the number of fMRI data points fourfold (from 616 to 2464 scans). Thus, with the same scanning time, fast data sampling enables one to increase the temporal degree of freedom and thereby significantly increase the statistical power of TMFC methods.

Furthermore, fast fMRI data sampling may improve sensitivity *per se*, that is, by more precise insights into neuronal temporal dynamics rather than simply by providing more data points for a fixed scan time^49^. To test this assumption, in contrast to our previous simulations with fixed scan time and number of events, we considered a fixed number of data points (scans). As a result, the sensitivity of TMFC methods remained at the same level at different temporal resolutions, TR = 500/700/1000 ms, despite a reduction in total scan time and number of events per condition, N_e_ = 50/70/100 (Fig. 8e, left panel). This meant an increase in normalised sensitivity per single event (TPR/N_e_) at shorter TRs (Fig. 8e, right panel). The BSC-LSS method had the highest sensitivity per event for short event duration and fast fMRI data acquisition. Therefore, despite the sluggishness of haemodynamic processes, a more accurate characterisation of ultra-slow BOLD-signal fluctuations allows for more precise insights into rapid task-related modulation of gamma-band synchronisation.

### Haemodynamic variability markedly decrease sensitivity of all TMFC methods

In the previous sections, we considered simulations with fixed parameters of the Balloon-Windkessel haemodynamic model. However, it is well known that the HRF shape varies across brain regions and subjects^50^. To study this issue, we performed simulations with variable haemodynamic parameters providing a time-to-peak range of 3 to 7 s, consistent with empirical studies^51^. Here, we compared the best-performing methods (gPPI, BSC-LSS, BSC-FRR), which assume canonical HRF shape in general linear models (GLMs), and the CorrDiff approach, which does not use GLMs. The gPPI and BSC-LSS methods were compared with and without upsampling of the task design matrix prior to convolution with canonical HRF. The upsampling procedure can improve sensitivity for an fMRI signal with a fixed canonical HRF, however its interaction with variable HRF is unknown. Additionally, we compared the gPPI method with and without deconvolution, since this procedure assumes a canonical HRF shape.

As a result, we revealed a marked decrease in sensitivity of all TMFC methods (Fig. 9). For the block design with SNR = 0.4, the sensitivity of the gPPI method with task design upsampling and deconvolution dropped from 78% to 5% (Fig. 9a). Disabling upsampling and deconvolution further decreased sensitivity of the gPPI method. As the SNR increases to 0.7, the sensitivity of the gPPI method increased to 71%. Therefore, the gPPI method requires a relatively high SNR to estimate TMFC under conditions of variability in the HRF shape.

**Fig. 9.**
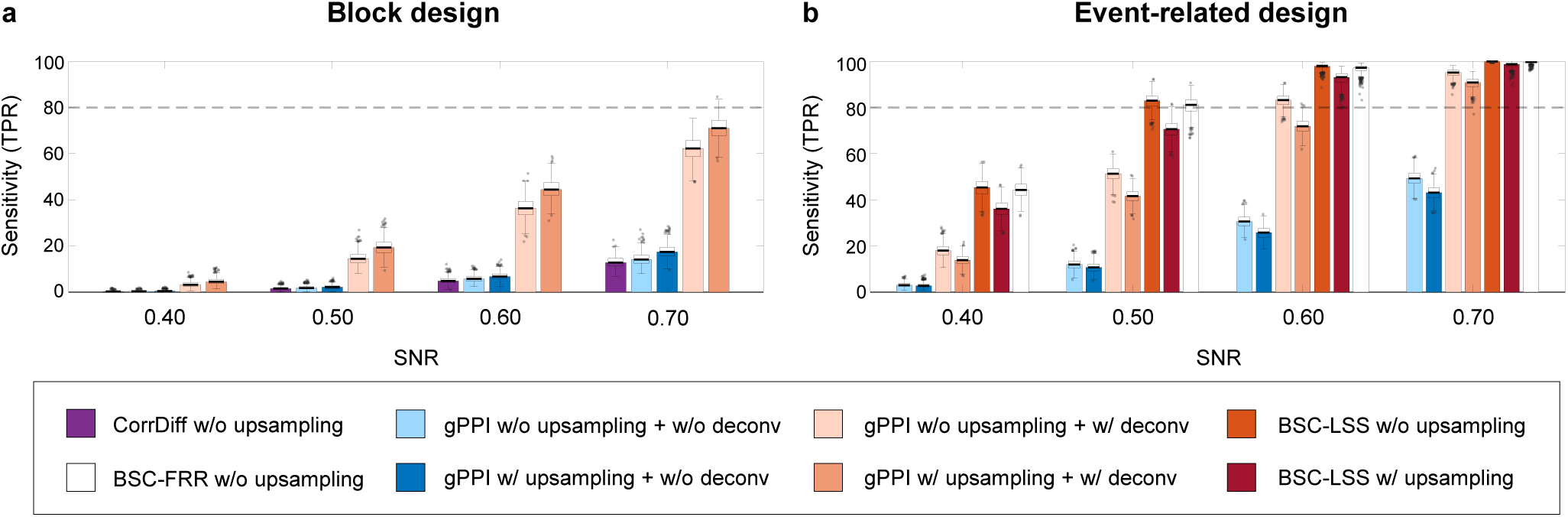
Influence of haemodynamic variability on the sensitivity of TMFC methods. Simulation results for sample size N = 100, TR = 2 s and variable parameters of the Balloon-Windkessel haemodynamic model. Sensitivity of different TMFC methods **(a)** for the block design with ten 20 s blocks per condition and **(b)** event-related design with one hundred 1 s events per condition and mean ISI = 6 s. The BSC-LSS and gPPI methods were implemented with (w/) and (w/o) upsampling of the design matrix. PPI terms were calculated with (w/) and without (w/o) the deconvolution step. gPPI matrices were symmetrised. Boxplots whiskers are drawn within the 1.5 interquartile range (IQR), computed from 1000 random resamplings with replacement.

For the default event-related design with SNR = 0.4, the sensitivity of the BSC-LSS and gPPI methods with upsampling dropped from about 93% to 36% and 14%, respectively (Fig. 9b). At the same time, the BSC-FRR sensitivity decreased from 91% to 44%, making it more robust to HRF variability than the BSC-LSS method. This robustness could potentially be due to fractional ridge regression or the lack of design matrix upsampling in the BSC-FFR method. It turned out that this was due to the second option. In the case of variable HRF simulation, the absence of upsampling of the task design matrix prior to convolution with canonical HRF increased the sensitivity of the BSC-LSS method from 36% to 45% (Fig. 9b). This may be due to the fact that convolution of an upsampled design matrix with a canonical HRF (convolution with high temporal resolution) implies greater model dependence on the canonical HRF shape than convolving the design matrix without upsampling (convolution with low temporal resolution).

Thus, the BSC-LSS method without upsampling was the most robust to haemodynamic variability. The BSC-FRR method without upsampling was slightly less sensitive than the BSC-LSS method, but still much robust than the gPPI method. Without upsampling, gPPI sensitivity was increased from 14% to only 18% for SNR = 0.4 (Fig. 9b). The gPPI method was half as sensitive than the BSC-LSS and BSC-FRR methods. Disabling deconvolution, reduced gPPI sensitivity to 3%, even though deconvolution assumes the canonical HRF shape. At higher SNRs, the difference in sensitivity between the gPPI and BSC methods became less noticeable.

### Genuine and spurious asymmetry of the PPI matrices

Previously, we considered only symmetrised PPI matrices because averaging the upper and lower diagonal elements has become a standard procedure in TMFC analysis since they are considered quite similar^8,11^. Here, we use empirical and simulated data to determine when PPI matrices become asymmetric, making the averaging procedure problematic. Below, we report results only for the gPPI method since the sPPI and gPPI methods produce nearly identical results.

For the block design (working memory task), the correlations between the upper and lower diagonal elements of the group-mean gPPI matrices without and with deconvolution were 0.85 and 0.77, respectively (Supplementary Fig. S7a). Without deconvolution, the correlation coefficients for individual subjects ranged from 0.77 to 0.89 with a mean of 0.83 (Supplementary Fig. S7c). With deconvolution, individual correlation coefficients ranged from 0.56 to 0.75 (mean 0.68). For the event-related design (stop-signal task), the correlations between the upper and lower diagonal elements of the group-mean gPPI matrices without and with deconvolution were 0.90 and 0.40, respectively (Supplementary Fig. S7b). The correlation coefficients for individual subjects ranged from 0.78 to 0.93 with a mean of 0.90 without deconvolution (Supplementary Fig. S7d). In contrast, when deconvolution was applied, individual correlation coefficients ranged from 0.24 to 0.41 (mean 0.34). Similar results were obtained for other tasks with block and event-related designs taken from the HCP and CNP datasets (Supplementary Fig. S8). Therefore, empirical data showed that deconvolution slightly increases the asymmetry of gPPI matrices for block designs and substantially increases the asymmetry for event-related designs.

Next, we used simulations with symmetric ground-truth matrices to determine which parameters of the fMRI experiment can artificially increase the asymmetry of the PPI matrices. We found that the main factors for artificial matrix asymmetry are a low SNR (Supplementary Fig. S9a), small sample size (Fig. S9b), short event duration (Fig. S9c) and small number of events per condition (Fig. S9e). In addition, the asymmetry of gPPI matrices increases slightly with larger TRs (Fig. S9f) and is practically independent of the mean ISI duration (Fig. S9d). Therefore, the large asymmetry of the gPPI matrices for the event-related designs from the CNP dataset is most likely related to the short scan time and low SNR.

Finally, we considered simulations with *asymmetric* ground-truth matrices (Supplementary Fig. S10), fixed HRF, and without adding co-activations to test whether the gPPI method could in principle provide information about the true causal directionality. The causal influence that one neuronal system exerts over another at a synaptic or neuronal population level is referred to as effective connectivity (EC). Here, we calculated correlation between the asymmetric ground-truth matrix and the asymmetric gPPI matrix, as well as the ratio between correctly identified sign of connections to the total number of non-zero ground-truth connections (correct sign rate, CSR, see Supplementary Information 9, Eq. S33). CSR of 50% means the sign was determined by chance. CSR of 100% means that the signs of all connections presented in the ground truth were correctly identified. For the block designs, we also calculated task-modulated EC (TMEC) matrices using the regression dynamic causal modelling (rDCM) method^52,53,54^ (see Methods). The rDCM method, which is a conventional EC method, was used as a reference. As rDCM requires a relatively high SNR^52^, we used SNR = 5 and twice the total scan duration. If the gPPI method fails to correctly estimate the direction of information flow at a high SNR, then it will also fail at lower SNRs. A systematic comparison of TMEC methods such as Granger causality or structural equation modeling is beyond the scope of the current study.

As a result, the gPPI method with deconvolution was able to reflect the actual direction of the information flow for the block design with twenty blocks per condition (Fig. 10). The correlation between the group-mean asymmetric gPPI matrix and ground-truth matrix for the “Cond A – Cond B” difference was 0.93 (CSR = 100%), and correlation between the group-mean rDCM matrix and ground-truth matrix was 0.88 (CSR = 99%). We also ensured that the asymmetry of gPPI regression coefficients was not due to amplitude differences between ROIs during task conditions (see Supplementary Information 9, Fig. S11, S12).

**Fig. 10.**
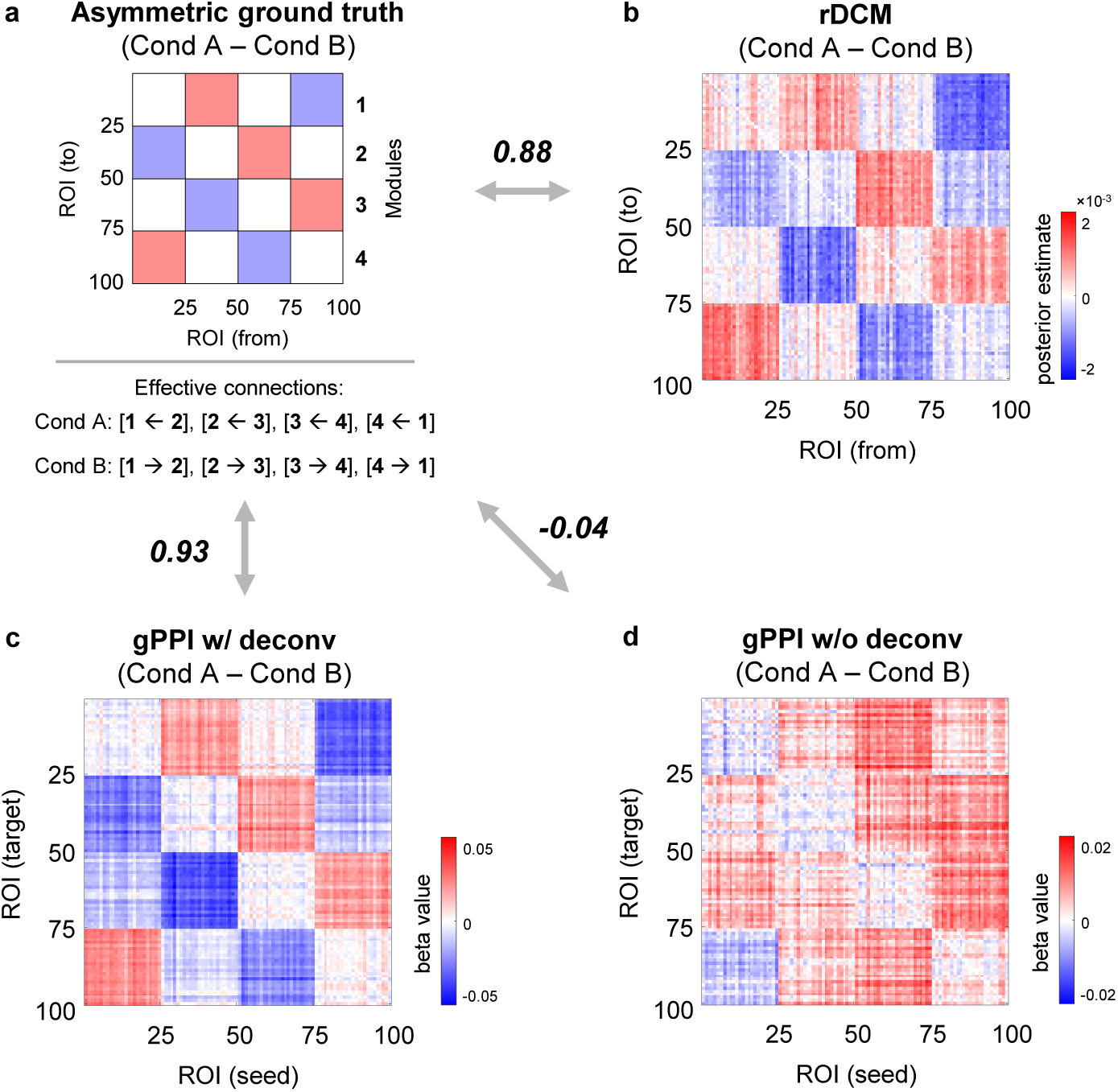
gPPI with deconvolution can reveal the direction of information flow under fixed HRF, high SNR and long scan time. Simulation results for the block design with twenty 20 s blocks per condition, sample size N = 100, TR = 2 s and very high SNR = 5. **(a)** Asymmetric ground-truth matrix of task-modulated effective connections. In “Cond A”, synaptic weights were increased from module №1 to №4, from №4 to №3, from №3 to №2, and from №2 to №1. In “Cond B”, synaptic weights were increased in the opposite direction. **(b)** Group-mean difference between rDCM matrices calculated for the “Cond A” and “Cond B” blocks. **(c)** Group-mean asymmetric matrix generated by gPPI with (w/) deconvolution. Deconvolution allows modelling of psychophysiological interactions at the neuronal level. **(d)** Group-mean asymmetric matrix generated by gPPI without (w/o) deconvolution. Without deconvolution psychophysiological interactions, are modelled at the haemodynamic level. To evaluate the similarity between matrices, we used Pearson’s r correlations. The color scales were adjusted for each matrix based on the maximum absolute value and were assured to be positive and negative symmetrical.

When the scan duration was halved, the correlation between the gPPI and ground-truth matrices decreased to 0.76 (CSR = 88%), and dropped to 0.12 (CSR = 56%) when SNR was reduced to 0.5. Similar results were obtained for the event-related designs: the correlation between the gPPI and ground-truth matrices was 0.85 (CSR = 99%) for two hundred events per condition, decreased to 0.57 (CSR = 75%) when the scan duration was halved, and dropped to 0.03 (CSR = 51%) when SNR was reduced to 0.5. Without deconvolution, the correlation between the ground-truth and gPPI matrices was always close to zero (|r| < 0.05, CSR ≈ 50%).

Therefore, gPPI without deconvolution failed to estimate the actual direction of the information flow, determined by asymmetric ground-truth synaptic weights, even in the best-case scenario (fixed HRF, high SNR, long scan duration). At the same time, gPPI method with deconvolution was able to reveal the true causal directionality in the best case. However, none of the connections estimated by gPPI survived the FDR-corrected threshold of 0.001 (even though the connection signs were correctly identified). Moreover, when we shortened the scan duration, reduced the SNR, and, most importantly, introduced HRF variability, the ability of the gPPI method to correctly identify the direction of information flow was reduced to almost zero.

## Discussion

This is the first evaluation of existing whole-brain task-modulated functional connectivity (TMFC) techniques using biophysically realistic large-scale neural mass simulations for a wide range of fMRI experimental settings. We identified the most effective TMFC methods for block and event-related designs, determined which data analysis procedures and parameters of the fMRI experiment increase their sensitivity and specificity, and demonstrated the principal capability of fMRI to detect rapid task-related neuronal synchronisation from sluggish BOLD signals at various temporal resolutions.

The simplest and most intuitive approach for constructing a whole-brain TMFC matrix is to directly calculate the correlation difference (CorrDiff) between task conditions, accounting for transient haemodynamic effects^11^. However, this approach is suitable only for task designs with long blocks since there must be enough time points after the removal of the transition periods (first six seconds of each block). We determined that the CorrDiff method has substantially lower sensitivity than other methods since cutting out the transition periods significantly reduces the temporal degrees of freedom.

A more sophisticated TMFC method available for both block and event-related designs is the psychophysiological interaction (PPI) approach^12^. The psychophysiological interaction can be modelled at the haemodynamic level or at the neuronal level using the deconvolution procedure^15^. Standard PPI (sPPI) approach was originally proposed for seed-to-voxel analysis and optimised for designs with two task conditions^12^. Later, it was extended to a generalised form (gPPI) that enables the assessment of more than two task conditions per statistical model^13^ and adapted for whole-brain ROI-to-ROI analysis^36,37,38^. In this study, we established that the most sensitive TMFC techniques for block designs and rapid event-related designs (mean ISI ≤ 4 s) are the sPPI and gPPI methods incorporating the deconvolution procedure. Although previous simulation studies suggested that the gPPI method is more sensitive than the sPPI method^13,22^, we showed that the sensitivity of both methods is practically equivalent for block and event-related designs. This discrepancy is most likely due to the difference between previously used biologically simplified simulations based on box-car or delta functions^13,22^ and biologically realistic simulations of TMFC based on the large-scale neural mass model used in the present work. Nevertheless, the gPPI method is preferable to the sPPI method since it provides more flexibility regarding task design and higher specificity if co-activations have not been removed before TMFC analysis using finite impulse response (FIR) task regression.

Previous empirical^55^ and simulation studies^15,24^ showed that the PPI approach with and without deconvolution produces similar results for block designs. However, disparities between PPI terms calculated at the haemodynamic and neuronal levels were more prominent for event-related designs since there are more high-frequency components^15^. Some authors suggest that the deconvolution step is mandatory for event-related designs^8^, while others warn against its use, arguing that there is no deterministic way to deconvolve the haemodynamic response if its shapes are not known exactly^33^. While the deconvolution step is a default setting in SPM, it is not implemented in other popular neuroimaging software packages, such as FSL and the CONN toolbox. Based on our results, we argue that deconvolution can increase sensitivity by up to a factor of two for both event-related and block designs. Moreover, deconvolution can increase specificity several fold, effectively eliminating co-activations even without FIR regression. Without deconvolution and FIR task regression, the gPPI method is susceptible to spurious inflation of TMFC estimates due to simple co-activations.

Since PPI is a regression method, it produces asymmetric (directed) TMFC matrices. However, to date there is no consensus whether PPI can be used to reveal the direction of information flow. Some authors used PPI without deconvolution as an effective connectivity method^40^. Others have used PPI with deconvolution to determine the direction of connectivity without making strong claims about the underlying causal structure^38^. A popular view is that by using PPI we are making arbitrary directional assumptions based on a priori assignment of ROIs as sources or targets^14,34^. Here, we showed that PPI can in principle reveal the direction of information flow, but only in the best-case scenario (fixed HRF, high SNR, long scan time) and only if the psychophysiological interaction is modelled at the neuronal level by using the deconvolution procedure. Without deconvolution, the PPI method provide no information about the underlying causal structure even in the best-case scenario. In addition, we found that PPI matrices asymmetry can spuriously arise due to low SNR, small sample size, short scan time (small number of events) and short event duration. To make correct directional inferences, the PPI method, as well as the more sophisticated effective connectivity method, regression dynamic causal modelling (rDCM), requires low haemodynamic variability across brain regions, high SNR and long scan times.

To create symmetrical PPI matrices, it was proposed to average upper and lower diagonal elements based on the empirical evidence that these elements are strongly correlated in block design tasks^11^. However, for task designs with low SNR and short scan time, the PPI matrices can become largely asymmetric, making the averaging procedure problematic. To completely avoid the asymmetry problem, it has been proposed to calculate symmetric TMFC matrices using correlational form of PPI (cPPI) based on partial correlations^14^. The cPPI method has not been previously validated on empirical or simulated data or compared with other TMFC methods. Here, we demonstrated that the cPPI method cannot estimate TMFC: it is unable to eliminate spontaneous task-independent activity and produces matrices similar to the task-state functional connectivity (TSFC) and background functional connectivity (BGFC) matrices that reflect the sum of FC across all task conditions and intervening rest periods.

An alternative method for obtaining symmetrical whole-brain TMFC matrices for event-related designs is the beta-series correlation (BSC) approach^16,17^, which is based on correlations between beta estimates of the BOLD signal change for each individual trial. A simulation study by Cisler et al.^22^ suggested that the BSC approach is more powerful than gPPI for event-related designs with many trial repetitions, while an empirical study by Di & Biswal^8^ with a large sample size did not support this notion. The inconsistency between the results of these studies may be due to the discrepancy between real neurophysiological processes that cause TMFC and the biophysically unrealistic simulations used to mimic TMFC^22^. We determined BSC based on the least-squares separate (LSS) method is superior for most types of event-related designs (mean ISI > 4 s). The increase in sensitivity compared to that of the gPPI method is especially noticeable for short event durations (< 1 s). At the same time, BSC based on the least-squares all (LSA) method had the lowest sensitivity due to the multicollinearity problem (resulting in noisy individual trial estimates) and achieved reasonable sensitivity only for long ISI (≥ 12 s) and/or long event duration (≥ 2 s). Note that one of the popular BSC software packages (BASCO toolbox) utilises the LSA method.

We also tested for the first time the application of fractional ridge regression (FRR) to BSC analysis^18,19^. The LSS can be thought as an extreme regularisation approach, as it shrinks noisy parameter estimates toward more stable values uniformly across all brain regions^18^. At the same time, the FRR approach uses the same general linear model (GLM), as in the LSA approach, but applies ridge regression and cross-validation to determine the optimal shrinkage fraction of individual trial estimates for each voxel or brain region. Therefore, the FRR approach can be thought as a tunable regularisation where regularisation is applied only if the data need it. As a result, the BSC-FRR approach turned out to be slightly less sensitive than the BSC-LSS approach, but at the same time noticeably more computationally efficient (faster than one and a half times compared to LSS).

One of the objectives of this study was to evaluate the influence of task design parameters on the sensitivity of TMFC methods. The two most important factors influencing sensitivity are the duration of a single trial and the number of trial repetitions. A noticeable decrease in sensitivity can be observed for trial durations shorter than 500 ms. The BSC-LSS method is the most robust to shortening trial duration. In addition, TMFC analysis requires a relatively large number of trial repetitions. Preferably, at least 80-100 trial repetitions per task condition. The block designs can be thought as designs with few trial repetitions and long trial durations. For them, the gPPI method with deconvolution is most sensitive. The sensitivity of BSC-LSS for block designs is very small, since it is based on correlations between individual trial or block estimates, the number of which is small in block designs. It is recommended to use Pearson correlation only if there are more than 25 data points^56^. However, block design with a large number of blocks becomes unreasonably long.

Another factor that may influence the sensitivity of TMFC methods is the mean interstimulus interval (ISI). As ISI decreases (mean ISI ≤ 4 s), the sensitivity drops relatively little, being more noticeable for BSC methods. The most robust to short ISI was the gPPI method with deconvolution. Increasing the mean ISI beyond 6 s did not affect sensitivity. Ideally, to improve the power of the fMRI design for assessing TMFC, trial repetitions, trial duration, and mean ISI should be increased. However, in real-world cases, there are practical limitations of the total duration of the fMRI study (e.g., fatigue of subjects, habituation to the task, limited access time to the MRI scanner, etc.). Moreover, in many cases, increasing the trial duration is impossible, since it depends on the nature of the psychophysiological processes under investigation (e.g., it is impossible to increase the duration of subliminal stimuli perception). Based on our simulations, we can infer what trade-offs can be made when designing fMRI experiments. Given that shortening the ISI reduces the sensitivity remarkably less than reducing number of trial repetitions, a rapid event-related design with a short ISI and a large number of events is more preferable than a slow event-related design with a small number of events. If possible, it is advisable to keep the event duration to about a second. For instance, an event-related design with 200 trial repetitions for each of the two task conditions, a trial duration of 1 s, and a mean ISI of 4 s would take 33.3 min. This task can be spitted into three 10-minute sessions, which is reasonable for an fMRI study. Alternatively, one can reduce the number of trial repetition (e.g., to 100 trials per condition) and the total task duration, if fast fMRI data acquisition is applied (e.g., TR = 0.72 s, as in the HCP dataset).

Importantly, we showed that short-term (100 ms) modulations of gamma-band neuronal synchronisation can in principle be recovered from ultra-slow BOLD signal fluctuations, even at typically low temporal resolution (TR = 2 s). However, this requires many trials (> 200 per condition). Furthermore, fast fMRI data acquisition increases the sensitivity for short-term modulations not only by increasing the number of data points for a fixed scan time but also by providing more insights into neuronal temporal dynamics. Our results show the particular importance of fast fMRI sequences for TMFC analysis. Fast data acquisition (TR < 1 s) prominently increased the sensitivity of all TMFC methods, even when the number of data points (fMRI scans) was fixed. In the case of slow data acquisition (TR = 3 s), the gPPI method with deconvolution was the most sensitive method.

Finally, we examined how heamodynamic variability across brain regions and subjects influences TMFC assessment. The introduction of HRF variability into simulations markedly decreased the sensitivity of all TMFC methods. It also abolished the ability of gPPI with deconvolution to determine the direction of information flow. The BSC-LSS and BSC-FRR methods were found to be much more robust to heamodynamic variability than the gPPI method. One possible explanation is that the latter makes more assumptions about the canonical shape of HRF. gPPI uses this assumption to create a convolved task design matrix for the GLM and to perform deconvolution and re-convolution of the PPI term, whereas BSC methods assume a canonical HRF only in GLMs. However, we argue for a different explanation.

First, disabling deconvolution did not increase, but rather decreased gPPI sensitivity. Second, we can see a dramatic reduction in the sensitivity of the CorrDiff approach, which does not use GLMs and does not directly assume canonical HRF shape. It simply calculates correlation difference between time points related to different task conditions, cutting out transitory periods. We removed the first 6 s in each block. Changing the length of these periods from 3 to 7 s did not significantly change CorrDiff sensitivity. This means that haemodynamic variability across brain regions decreases correlations between time series from different brain regions *per se*, without any model assumptions about the canonical HRF shape. The gPPI method can be thought of as the difference in weighted correlations between time series relating to different task conditions^57^. Meanwhile, BSC methods are not based on correlations between time series, but on correlations between trial-by-trial amplitude fluctuations. Although estimating response amplitudes of individual trials requires an assumption the HRF shape, it turned out, that correlations between trial-by-trial amplitude fluctuations are more robust to HRF variability than correlations between time series. The block design is generally considered more statistically efficient in fMRI studies (especially for activation studies). However, our results suggest that, counterintuitively, event-related designs with a large number of trial repetitions may be more powerful for TMFC assessment in the case of high haemodynamic variability, as they enable analysis of trial-by-trial amplitude variations. To demonstrate this, we considered block and event-related simulations with comparable total scan duration, variable HRF and SNR = 0.5. For a block design with 20 blocks per condition and a total duration of 26.6 minutes, the gPPI sensitivity was 56%, whereas the BSC-LSS sensitivity was 6%. At the same time, for an event-related design with 100 events per condition and a total duration of 23.6 minutes, the sensitivity of the gPPI and BSC-LSS methods was 51% and 83%, respectively.

In conclusion, we provide practical recommendations for TMFC analysis:

1. Remove co-activations from time series using FIR regression, especially if you use gPPI without deconvolution or sPPI with/without deconvolution.
2. For primary TMFC analysis, use:

a. gPPI for block designs and rapid event-related designs (mean ISI ≤ 4 s), if low HRF variability is expected;
c. gPPI for fMRI studies with long TR (≥ 3 s), if low HRF variability is expected;
b. BSC-LSS in all other cases (especially if high HRF variability is expected.
3. Perform cross-method validations (secondary TMFC analysis):

a. High correlation between raw TMFC matrices calculated by gPPI and BSC methods suggests that the results are reliable;
b. Low correlation indicates insufficient or low-quality data for TMFC analysis.
4. Use the deconvolution procedure to increase sensitivity and specificity of PPI methods.
5. Report whether mean centering was applied to the task regressor prior to PPI term calculation. Interpret condition-specific gPPI matrices (“Condition > Baseline”) with caution regarding connections that exhibit high connectivity during rest periods. The sign of these connections may be reversed to negative due to the gPPI model and mean centering.
6. When averaging the upper and lower elements of the PPI matrix, report its symmetry:

a. High correlation between the upper and lower elements of the PPI matrix suggests that the results are reliable;
b. Low correlation indicates insufficiency or low quality of data for TMFC analysis.
7. Draw conclusions about the direction of information flow from PPI results with caution:

a. Use the deconvolution procedure. Without deconvolution, PPI methods do not provide information about directional information flow.
b. Make sure your data has low HRF variability, a high enough SNR, and long scan duration.
c. Support preliminary whole-brain PPI results with confirmatory analysis for several selected ROIs using an advanced effective connectivity method such as DCM.
8. Do not use the LSA method for BSC analysis (especially if mean ISI < 12 s). The BSC-LSS method is preferred, however the BSC-FRR method can be also be used to reduce computation time.
9. If low HRF variability is expected, one can apply task design upsampling prior to convolution with HRF. Otherwise, task design upsampling should not be used (e.g., set the microtime resolution in SPM to one).
10. When TMFC analysis is the main goal:

10.1 Avoid using very rapid event-related designs (mean ISI < 2 s);
10.2 Increase the number of events as much as possible (TMFC analysis requires more events per condition than does the activation analysis);
10.3 Prefer a larger number of events to a long ISI;
10.4 Use a longer event duration (if possible);
10.5 Fast fMRI data acquisition is preferable (TR < 1 s)
10.6 If high HRF variability is expected, consider using an event-related design a large number of events instead of a block design.

We make all simulated time series and code publicly available, along with Jupyter notebooks for replication of our results (https://github.com/IHB-IBR-department/TMFC_simulations). These simulated time-series can be used in future studies for validation and comparison of new TMFC or TMEC methods. The code for TMFC simulation is also available as a separate Python module *TMFC_simulator* (https://github.com/IHB-IBR-department/TMFC_simulator). *TMFC_simulator* can be used not only to compare TMFC methods, but also to facilitate the design of fMRI experiments. The most effective task design can be selected from various possible design options based on simulations with a given signal-to-noise ratio and haemodynamic variability.

We also provide a user-friendly SPM12-based toolbox with GUI and parallel computing capability for voxel-based and ROI-to-ROI TMFC, called *TMFC_toolbox* (https://github.com/IHB-IBR-department/TMFC_toolbox). It implements two of the most sensitive methods covering all types of fMRI designs: BSC-LSS and gPPI with deconvolution. It also allows to perform FIR task regression along with TMFC analysis. Additionally, we provide MATLAB and Python functions for performing deconvolution based on ridge regression that reproduce results of deconvolution implemented in SPM12 (https://github.com/IHB-IBR-department/BOLD_deconvolution). These functions can be useful for performing PPI analysis independently of SPM12 and/or MATLAB.

There are several issues that were not addressed in the present study, providing opportunities for future research. We manipulated synaptic weights between neural masses, which caused modulations of gamma-band neuronal oscillations and synchronisation. This, in turn, led to cascaded modulation of ultra-slow gamma-band envelope fluctuations, haemodynamic fluctuations, and BOLD-signal fluctuations. We assessed how different TMFC methods estimate these modulations depending on the task design, TR, SNR, sample size, presence of co-activations, and haemodynamic response variability across brain regions and subjects. However, we did not consider a number of factors that may also affect TMFC estimates.

First, neuronal synchronisation or coherence can be observed not only for oscillations, measured by local field potential (LFP), but also for spiking activity across multiple cortical areas, measured by single-unit activity (SUA) or multi-unit activity (MUA), which also influence BOLD-signals^58,59^. Moreover, concurrent non-oscillatory aperiodic activity can be misinterpreted as oscillatory activity and confound measures of oscillatory activity^60^. One important extension of this study would be simulation of fast aperiodic neuronal activity and its influence on neuronal synchronization and TMFC estimates.

Second, neuronal synchrony can by modulated not only by short-term plasticity (e.g., synaptic facilitation/depression or spike-timing dependent plasticity), but also by other mechanisms. For example, synchronisation between downstream regions can be influenced by the temporal coordination of spiking activity in source region^61^. Synchronisation also depends on the precise timing of the sender and receiver regions, namely, a sending group of neurons will have the highest impact on a receiving group, if its inputs consistently arrive when synaptic gain is high^62^. The rapid changes in balance between excitatory and inhibitory activity continuously modulate neuronal synchrony depending on stimulus and behavioral state^63,64^. Regional stimulation modulates synchrony depending on the brain network’s collective dynamical state^25,65,66^. Besides, synchronization can be controlled by astrocytic modulation, which is rarely taken into account in computational models^67^.

Third, ultra-slow arteriole diameter fluctuations can depend not only on ultra-slow envelopes of higher-frequency bands, but also on ultra-slow aperiodic neuronal activity, such as the slow cortical potential^68,69^. Although a number of studies have shown that an electrophysiological signal with spectral components below 1 Hz can have a non-neuronal origin and themselves caused by cerebral vasomotion^70,71,72^.

Fourth, neuronal activity and ultra-slow hemodynamic fluctuations can depend on neuromodulatory input from subcortex (e.g., cholinergic or noradrenaline modulatory centers) and arousal level^71,72,73^. Fifth, cardiorespiratory processes influence arteriole diameter fluctuations and BOLD signal fluctuations (cerebral blood flow and oxygen concertation). TMFC estimation can be affected by imperfect denoising procedures and aliasing of physiological rhythms^71,74^. Fast fMRI techniques not only improve TMFC estimates, as shown in the present study, but can also help avoid aliasing effects^75^. Although future studies should also introduce a penalty of reduced SNR per scan due to reduced longitudinal magnetisation recovery for fast fMRI acquisition. Finally, TMFC estimates can be affected by head motion, especially if motion is correlated with task performance^74,76,77,78^.

It will also be important for future research to investigate how variability in neural mass model parameters affect TMFC estimates. In our simulations, we used a fixed parameters for the large-scale neural mass model. To account for hierarchical heterogeneity in local circuit properties across cortical areas (e.g., excitatory-inhibitory balance), neural mass model can be parameterizing with T1w/T2w myelin gradients^79^, RSFC gradients^80^, and/or transcriptional variations in excitatory and inhibitory receptor gene expression^81^. In addition, we chose parameters such that each neural mass generated gamma-band oscillations. Although gamma-band power has been shown to be the closest electrophysiological correlate of spontaneous and evoked BOLD signals^58,69,72,82,83,84,85^, oscillations in other frequency bands can also influence BOLD-signals^86,87^. Future research should consider oscillations in other frequency bands^86^ and cross-frequency coupling^64,88,89^ with respect to TMFC estimation.

Other remaining issues that could be addressed in future research are the influence of ROI selection (parcellation scheme) on TMFC estimates^90,91,92,93^, the best choice of deconvolution method for PPI analysis (e.g., consideration of blind deconvolution methods, that do not assume a canonical HRF shape), as well as region- and subject-specific HRF selection for the PPI and BSC methods. Finally, the proposed simulation approach would be useful for comparison of the whole-brain task-modulated effective connectivity methods (TMEC), such as rDCM, Granger causality, structural equation modelling, transfer entropy, Bayesian nets, Patel’s pairwise conditional probability, and others^94,95^. In the present study, the rDCM method was used to demonstrate that the direction of information flow can in principle be estimated for our simulations if SNR is high and HRF has a canonical shape.

## Methods

### Empirical data

We analysed preprocessed fMRI data for two block design tasks (working memory and social cognition tasks, N = 100) from the HCP dataset^41,43^, two event-related tasks (stop-signal and task-switching tasks, N = 115) from the CNP dataset^42,44^ and resting-state data from both datasets. For details about the scanning parameters and task designs, see Supplementary Information 2. The HCP and CNP preprocessing pipelines included realignment, spatial artefact/distortion correction, co-registration between functional and structural images, and normalisation to Montreal Neurological Institute (MNI) space. The CNP pipeline also included a slice-timing correction. Additionally, we smoothed the functional data with a 4 mm Gaussian kernel using SPM12 (https://www.fil.ion.ucl.ac.uk/spm/software/spm12).

To extract region-wise time series, we used a set of 300 functionally-defined ROIs published by Seiztman et al.^45^ The full set of functional ROIs consists of 239 cortical ROIs (most part of them taken from Power et al.^96^), 34 subcortical, and 27 cerebellar ROIs. Each volumetric ROI represents a sphere with a radius of 4 or 5 mm. We discarded ROIs for which data were incomplete for at least one subject. As a result, we utilised 239 ROIs for the HCP dataset and 246 ROIs for the CNP dataset.

### Large-scale neural mass model

The simulation procedure included five steps. First, we simulated gamma-band oscillatory neuronal activity for 100 interconnected brain regions using Wilson-Cowan equations^26,27^ and manipulated the synaptic weights depending on the task conditions to control the ground-truth TMFC. Transient activity-dependent modulation of synaptic strength, lasting from tens of milliseconds to several minutes, is referred to as short-term plasticity^29,30,31,32^. Second, we independently simulated co-activations using box-car functions to evaluate their impact on spurious inflation of TMFC estimates. Third, we applied the Balloon-Windkessel haemodynamic model^28^ to convert oscillatory neuronal activity and co-activations into BOLD signals. Fourth, we downsampled the BOLD signal to different time resolutions to assess the potential benefits of fast data acquisition for TMFC estimation. Fifth, we added white Gaussian noise to model scanner measurement error.

Our simulation approach expands previous TMFC simulation studies in several ways. First, we used a large-scale neural mass model instead of delta or boxcar functions^13,15,22,23^. Second, we applied biophysically realistic simulations not only to block designs^24,25^ but also to different event-related designs. Third, previous biophysically realistic simulations have indirectly modulated FC by injecting task stimulation into brain regions^24,25^. In the current work, we directly manipulated ground-truth FC by changing synaptic weights between neuronal units depending on the task context, which corresponds to short-term synaptic plasticity^29,30,31,32^. Finally, we ensured that the neural mass model generated spontaneous oscillations in the gamma band. In contrast, the most recent TMFC simulation study by Cole et al.^25^ used a large-scale neural mass model without inhibitory subpopulations, where limit-cycle oscillations cannot emerge under any model parameters (Supplementary Fig. S13). For a more detailed overview of the results and limitations of previous TMFC simulation studies, see Supplementary Table S2.

The Wilson–Cowan neural mass model achieves a good balance between biophysical realism and mathematical abstraction^27,87,97^. According to this model, a single neuronal population can be represented as synaptically coupled excitatory and inhibitory subpopulations described by two ordinary differential equations with a non-linear saturation function^26^. Each population can produce self-sustained limit-cycle oscillations as a result of feedback between coupled excitatory and inhibitory subpopulations. The Wilson-Cowan units were connected through the excitatory subpopulations^27,66,97^ with a signal transmission delay of 25 ms^98^. In line with previous large-scale neural mass simulations^27,66,97^, we chose to set the model parameters such that each coupled Wilson-Cowan unit produced gamma-band oscillations (≈ 40 Hz) due to the following considerations.

A large body of literature demonstrates that gamma-band oscillations and synchronisation are linked to sensory processing^99,100^, motor acts^101^, and cognitive processes^102,103^ and are thought to underlie information processing and transmission^64,89,104,105,106^. The spectral power (or envelope) of gamma-band oscillations fluctuates very slowly with time, and brain regions with shared function demonstrate co-fluctuation of gamma-band envelopes^107,108^. At the same time, a multitude of animal and human studies have shown that local field potential power in the gamma band is the closest electrophysiological correlate of spontaneous and evoked BOLD signals^58,69,72,82,83,84,85^. Moreover, trial-by-trial BOLD fluctuations are positively correlated with trial-by-trial fluctuation in gamma power during task performance^109^. The mechanisms for the relationship between neuronal activity and BOLD signals have not yet been fully determined. However, a recent study by Mateo et al.^46^ elucidated these mechanisms using optogenetic manipulations and concurrently measuring local field potential, arteriole diameter and blood oxygenation in awake mice. They provided direct evidence that an increase in gamma-band power leads to an increase in arteriole diameter, and an increase in arteriole diameter leads to an increase in blood oxygenation. This chain of processes can be described by a coupled oscillator model (Fig. 4a). For details on the applied model and its parameters, see Supplementary Information 3.1 and Supplementary Table S9. Synaptic activity was calculated as the sum of all inputs to excitatory and inhibitory subpopulations of the Wilson-Cowan unit and was considered a proxy for the local field potential^24,110^.

The code for TMFC simulation is available as a Python module *TMFC_simulator* (https://github.com/IHB-IBR-department/TMFC_simulator), which is an extension of the *neurolib* software (https://github.com/neurolib-dev/neurolib). *neurolib* is a Python library that provides a computationally efficient framework for whole-brain resting-state functional connectivity (RSFC) simulations^111^. The *TMFC_simulator* extension allows to manipulate synaptic weights depending on the task condition, simulate co-activations, change repetition time (TR), vary Balloon-Windkessel parameters, and convert excitatory and inhibitory activity into synaptic activity.

### Ground-truth synaptic weight matrices

The construction of the synaptic weight matrices involved three steps. First, we drew synaptic weights for each subject from a Gaussian distribution with a mean of one and a standard deviation of 0.1. Then, we multiplied the synaptic weights within and between functional modules by weighting factors that determined the network structure (Supplementary Table S10). Finally, we normalised the synaptic weights so that all inputs to each region summed to one, following previous simulation studies^25,66^.

### Analysis of simulated synaptic activity

We additionally analysed simulated synaptic activity before converting it into the BOLD signal. Initially, we downsampled the raw time series from a temporal resolution of 0.1 ms to 5 ms and bandpass filtered it in a narrow carrier frequency range [*f_carrier_* – 2 Hz, *f_carrier_* + 2 Hz], where *f_carrier_* = 40 Hz^86^. Next, we employed the Hilbert transform to obtain the instantaneous amplitudes and phases of the narrowband signal. The instantaneous amplitudes, or amplitude envelopes, were further cross-correlated with the simulated BOLD signals, while the instantaneous phases were utilised to estimate gamma-band neuronal synchronisation based on the phase-locking value method^112^.

### BOLD signal generation

The synaptic activity was converted to the BOLD signal using the Balloon-Windkessel haemodynamic model^28^. For more information about the model, see Supplementary Information 3.2. For simulations with the fixed HRF shape, we used the standard parameters taken from Friston et al.^28^ (Supplementary Table S11), which have previously been used in whole-brain RSFC simulation studies^27,66,97,113^. For simulations with variable HRF shape, we randomly sampled Balloon-Windkessel parameters for each ROI and subject from Gaussian distributions that provide time-to-peak variability of 3–7 s (Supplementary Table S12), consistent with empirical data^51^.

The BOLD signal generated from fast oscillatory activity corresponded to task-independent spontaneous fluctuations and task-modulated fluctuations during the rest and task conditions. To simulate the BOLD signal related to simple co-activations, we used boxcar activation functions as input to the Balloon-Windkessel model. Boxcar activation functions were set equal to one during task conditions and equal to zero at all other times. We normalised the BOLD signal related to co-activations such that the ratio between the standard deviations of the oscillatory-related signal and co-activation-related signal was determined by the scaling factor: SF = *σ_oscill_*/*σ_coact_*. In all simulations with co-activations, we set SF = 1. After the FIR task regression, the SF value does not influence the sensitivity and specificity of TMFC methods (Supplementary Fig. S14).

Next, we downsampled the raw BOLD signals from 0.1 ms to distinct time resolutions corresponding to different TRs and added white Gaussian noise as measurement error. The SNR was defined as the ratio between the standard deviations of the signal and noise, SNR = *σ_signal_*/*σ_noise_*, which is commonly used in DCM studies^52,53^. For simulations with fixed HRF shape, we varied the SNR between 0.3 and 0.5, and the default SNR was 0.4. For simulations with variable HRF shape, we varied the SNR between 0.4 and 0.7, since haemodynamic variability significantly reduced sensitivity of all TMFC methods. To compare the gPPI and rDCM methods, we increased the SNR to 5, as the rDCM method requires a very high SNR.

### Simulation experiments and task designs

By default, we considered symmetric synaptic weight matrices (Fig. 4c). The block design included 10 blocks each for the “A” and “B” conditions, alternating with “Rest” blocks. Each block lasted for 20 s. The total duration of the block design was ≈ 13 minutes. The default event-related design included 100 events for the “A” and “B” conditions interleaved by “Rest” periods. Each event lasted for 1 s. The ISI was randomly jittered between 4–8 s (mean ISI = 6 s). The ISI was defined as the interval between the end of one event and the start of the next event. The total duration of the default event-related design was ≈ 24 minutes. In other event-related designs, we separately varied event duration = [100 ms, 250 ms, 500 ms, 1 s, 2 s, 4 s], number of events per condition = [20, 40, 60, 80, 100], and mean ISI = [2 s, 4 s, 6 s, 8 s, 12 s]. Stimulus onset timings were determined by Chris Rorden’s fMRI Design software (https://github.com/neurolabusc/fMRI-Simulator), which improves the statistical efficiency of the task design. By default, we used typical slow fMRI data acquisition, TR = 2 s. In a separate experiment, we varied TR = [500 ms, 700 ms, 1 s, 2 s, 3 s]. In a separate experiment, we considered asymmetric synaptic weight matrices (Supplementary Fig. S10). Simulations were performed for N = 100 subjects unless otherwise stated. We also performed simulations with task designs identical to the working memory and social cognition tasks from the HCP dataset, as well as the stop-signal and task-switching tasks from the CNP dataset.

### Resting-state, background and task-state functional connectivity

To calculate the empirical RSFC, BGFC and TSFC matrices, we used the CONN toolbox release 21.a (www.nitrc.org/projects/conn)^114^. The mean time series extracted from functional ROIs were filtered using a bandpass filter of 0.01 Hz to 0.1 Hz and corrected for head motion and physiological noise using 24 motion regressors^115^ and 6 anatomical component-based noise correction (aCompCorr) regressors^116^. Pearson’s r correlation coefficients were calculated for each pair of regions and then converted to Fisher’s Z. Empirical RSFC and TSFC matrices were calculated based on the whole time series of denoised resting-state and task-state BOLD signals, respectively. To calculate the BGFC matrix for the block design task, we regressed out the stimulus-evoked haemodynamic responses (co-activations) from the time series using an FIR basis set with 64 post-stimulus time bins of 0.72 s duration^9^. For the event-related task, we used an FIR basis set with 32 post-stimulus time bins of 1 s duration.

### Task-modulated functional connectivity

The TMFC matrices for the empirical and simulated data were constructed in a similar way. For all TMFC analyses, we considered differences between two task conditions: (1) “2-back” and “0-back” for the working memory task, (2) “Social interaction” and “Random interaction” for the social cognition task, (3) “Go” and “Stop” for the stop-signal task, (4) “Switch” and “No switch” for the task-switching task, and (5) “Cond A” and “Cond B” for simulated data. All TMFC matrices were calculated without and with FIR task regression prior to TMFC analysis. The FIR basis sets were the same as for the BGFC analysis.

### Direct correlation difference

To calculate the direct correlation difference between task blocks, we removed the first six seconds in each block to account for transient haemodynamic effects^8,11^. We then concatenated the time series for each task condition, calculated Pearson’s r correlation coefficients, and converted them to Fisher’s Z. Finally, we calculated the Fisher’s Z difference between the two conditions.

### Psychophysiological interaction

The PPI approach applies a general linear model (GLM) to reveal the relationships between the BOLD signal in the “target” region and several explanatory variables: (1) expected task-evoked haemodynamic responses (psychological regressor, reflecting co-activations), (2) BOLD signal in the “seed” region (physiological regressor, reflecting task-independent connectivity, similar to BGFC), (3) element-by-element product of psychological and physiological variables (psychophysiological interaction regressor, reflecting TMFC), and (4) non-neuronal nuisance variables (head motion and physiological noise). The standard PPI (sPPI) method includes one PPI regressor and one task regressor for the difference between task conditions^12^. The generalised PPI (gPPI) approach includes multiple PPI terms and task regressors for each task condition, which enables us to consider more than two task conditions in a single GLM^13^. To account for the fact that psycho-physiological interactions occur at the neuronal level, it was proposed to deconvolve the BOLD signal into underlying neuronal activity, calculate the element-by-element product of the estimated neuronal activity and unconvolved psychological regressor^15^, and then reconvolve this interaction term with HRF. For more details about different PPI approaches, see Supplementary Information 4.

For deconvolution, we used the SPM12 approach, which is based on representing unknown neuronal activity in the frequency domain as a linear combination of a full-rank cosine basis set^15^. Since parameter estimates based on a full-rank basis set are highly unstable (in particular for high frequencies, because the HRF selectively attenuates high frequencies), the estimates must be constrained or regularised^15^. For this purpose, SPM12 uses a Parametric Empirical Bayes (PEB) approach with Gaussian priors^15^. Alternatively, regularisation can be achieved using ridge regression^117^. In the main text, we provided results for sPPI and gPPI methods without and with deconvolution implemented in SPM12 (*spm_peb_ppi.m* function). Additionally, we provided MATLAB and Python functions for deconvolution based on ridge regression, independent of the SPM12 software (https://github.com/IHB-IBR-department/BOLD_deconvolution). Both PEB and ridge regression deconvolution approaches produce very similar results (see Supplementary Information 5, Fig. S15, S16). Importantly, these deconvolution methods do not require any knowledge about the timing of the task design, but do assume the canonical shape of the HRF.

All results for the sPPI and gPPI methods were obtained using mean centering of the task design regressor prior to PPI term calculation, as previously recommended by Di et al.^11^ In the supplementary, we provided the detailed explanation of the mean centering procedure and comparison of results obtained with or without mean centering (see Supplementary Information 6, Fig. S17, S18, S19). In short, mean centering did not change PPI parameter estimates without deconvolution, but did change them when deconvolution is applied. Primary, differences could be seen between nodes that exhibit high connectivity during rest periods in PPI matrices comparing the single task condition to the baseline. Without deconvolution, the PPI estimates between these nodes are negative, regardless of mean centering (Fig. S17a-b and S18a-b). The negative PPI estimates for the strongest task-independent connections are due to the gPPI model simultaneously fitting task-independent (physiological) and task-modulated (PPI) regressors. In the model without the physiological regressor, the negative PPI estimates arise from mean centering, which reverses the rest periods in the PPI regressor.

When deconvolution is applied, the PPI estimates between these nodes are positive without mean centering and negative with mean centering (Fig. S17c-d and S18c-d). If the psychological variable is non-centered with a constant component, the constant component will add a physiological variable to the PPI term^11^. After deconvolution and reconvolution, this physiological component is no longer exactly the same as the original physiological variable. In this case, the gPPI model fits the original physiological regressor and PPI regressor with a smoother physiological component. Since these physiological components are not identical, the sign reversal does not occur.

When considering the relative connectivity changes in one task compared with others (“Condition A vs. Condition B”, i.e., the main goal of TMFC analysis), mean centering did not affect PPI results for well-balanced task designs. In unbalanced designs (e.g., the stop-signal and task-switching tasks from the CNP dataset), the lack of mean centering led to false positive results between nodes that exhibit high connectivity during rest periods if the sPPI method is used (Fig. S19b, d). False positives could be avoided by applying mean centering to the sPPI method or using the gPPI method regardless of mean centering (Fig. S19a, c).

The PPI matrices consist of the beta estimates for the PPI regressors. The beta estimates change when the “target” and “seed” regions are swapped. However, some authors suppose that the directionality of the PPI approach is arbitrary, as it is based on the a priori designation of regions as either sources or targets rather than a dynamic model of causal effects posed at the neuronal level combined with a forward biophysical model linking neuronal dynamics with haemodynamic responses^34^. The correlational PPI (cPPI) approach is a modification of the sPPI approach that avoids arbitrary directionality assumptions^14^. It is based on the partial correlation between PPI terms, removing the variance associated with the psychological and physiological regressors. To calculate the group-mean cPPI matrices, we converted partial correlations to Fisher’s Z.

For all PPI analyses, we used SPM’s volume of interest function to extract time series from the ROIs. Representative ROI time series were extracted as first eigenvariates after signal pre-whitening (using an autoregressive model of order one, AR(1) model), high-pass filtering (cut-off of 128 s), and adjustment for effects of no interest (session-specific constant terms, head motion and aCompCorr regressors). Regression models for the sPPI and gPPI methods included task regressors regardless of whether FIR task regression was performed before TMFC analysis. All task regressors were convolved with canonical HRF implemented in SPM12.

### Beta-series correlation

The BSC approach is based on the correlations between beta estimates of stimulus-evoked haemodynamic responses for individual task trials. According to the originally proposed BSC approach, all events are modelled by separate regressors in a single GLM^16^. This BSC approach, also known as the “Least-Squares All” approach (LSA), was initially developed for slow event-related designs. However, the LSA approach is unsuitable for rapid event-related designs for two reasons. First, beta estimates can become unreliable due to correlations between regressors for individual trials. Shorter ISIs lead to higher correlations between regressors. This multicollinearity results in highly unstable (noisy) estimates due to a limited amount of variability unique to an individual event^17^. Second, rapid event-related designs consist of many trial repetitions due to the short ISI. As the number of individual trial regressors approaches the number of time points, the design matrix becomes close to singular (e.g., the stop-signal task consists of 126 trials and 182 time points) and cannot be reliably inverted using ordinary least squares^118^.

The alternative approach, called the “Least-Squares Separate” approach (LSS), solves this problem by modelling each individual event against all other events with separate GLMs^17^. This approach has two main drawbacks. First, the LSS approach shrinks noisy parameter estimates toward more stable values uniformly across all brain regions. Such approach ignores the fact that different brain regions may require different levels of shrinkage^19^. Second, the LSS approach is computationally expensive. For example, the stop-signal task consists of 128 trials and requires estimation of 128 GLMs. A possible solution maybe to use a new method called fractional ridge regression^18,19^. Previously, it was used to estimate regularised single-trial parameter estimates for brain decoding^119,120^, brain encoding^121,122,123^, reconstruction of viewed images^120,124^ and representational similarity analysis^125^. Here, we propose to apply FRR to BSC analysis (BSC-FRR).

The FRR procedure is a computationally efficient approach based on singular value decomposition of the design matrix and reparameterisation of the ridge regression in terms of the fraction between the L2-norms of the regularised and unregularised parameters^18,19^. It allows to determine the optimal level of shrinkage fraction (regularisation) for each voxel or brain region from the full range of regularisation levels in a fast and automated way^18^. For example, the BSC-LSS analysis took 4 hours 50 minutes for the stop-signal task dataset with parallel computations on an Intel Core i9-12900F 2.40 Hz (16-core) with 64 GB RAM, a 64-bit Windows 10 operating system, and MATLAB R2021b. On the same workstation and dataset, the BSC-FRR analysis took 2 hours 58 minutes. We used the FRR procedure implemented in the GLMsingle toolbox for MATLAB (https://glmsingle.org/)^18^. We selected the canonical HRF for the BSC-FRR analysis, as in all other TMFC analyses, and did not apply data-derived nuisance regressors.

We considered the BSC-LSA, BSC-LSS and BSC-FRR approaches. To obtain the BSC matrices, we calculated Pearson’s r correlations between the mean beta values extracted from ROIs separately for each task condition, converted them to Fisher’s Z and calculated the difference between Fisher’s Z values. For more details about different BSC approaches, see Supplementary Information 7.

### Task-modulated effective connectivity: regression dynamic causal modelling

The most popular EC method is DCM, which employs an explicit forward or generative model of how observed haemodynamic signals emerge from hidden neuronal and biophysical states^28^. Originally, DCM was introduced for fMRI data to describe changes in hidden neuronal states via a bilinear differential equation. It enables us to estimate the effective strength of task-independent (intrinsic) synaptic connections (*A* matrix), task-modulated (extrinsic) synaptic connections (*B* matrix), and the direct influence of driving inputs that cause activations (*C* matrix). The problem is that the original DCM is limited to parameter estimation of small brain networks (approximately ten nodes), as the model inversion becomes ill-posed and computationally demanding for large-scale (whole-brain) networks. To solve this problem, the regression DCM (rDCM) approach, which applies several simplifications to the original DCM, was proposed^52^. For more information about the rDCM approach, see Supplementary Information 8.

The task-modulated effective connectivity (TMEC) matrix cannot be directly obtained using the rDCM approach because it is based on the linear neural state equation without the *B* matrix. If we feed the entire time series of the resting-state or task-state BOLD signal into rDCM, the *A* matrix will reflect the resting-state effective connectivity (RSEC) and task-state effective connectivity (TSEC) matrices, respectively (cf. Fig. 1). In the latter case, the *A* matrix will depend on both spontaneous (intrinsic) and task-modulated (extrinsic) fluctuations. To calculate the TMEC matrix, we propose calculating two *A* matrices for concatenated “Cond A” and “Cond B” block time series after removing the first six seconds of each block. The difference between these matrices will subtract spontaneous (intrinsic) EC and result in TMEC^53^. The driving input *C* matrix was set to zero, as we removed the transition periods.

### Comparison of functional connectivity matrices

For the empirical data, we considered unthresholded weighted FC matrices. To evaluate similarity between the group-mean RSFC, TSFC, and TMFC matrices produced by different methods, we calculated the Pearson’s r correlations between the lower diagonal elements. To evaluate the asymmetry of the raw (i.e., before symmetrisation procedure) group-mean and individual gPPI matrices, we calculated the Pearson’s r correlations between the upper and lower diagonal elements.

For the simulated data, we compared the ground-truth TMFC matrix and thresholded binary TMFC matrices to evaluate the sensitivity and specificity of distinct TMFC methods (Supplementary Information 1). To obtain thresholded binary matrices, we determined significant FC differences between “Cond A” and “Cond B” using a two-sided one-sample t test. The Benjamini–Hochberg procedure^126^ was used to control for the false discovery rate (FDR) at the 0.001 level.

Additionally, we used Bayesian analysis to provide evidence for no difference between the sPPI and gPPI methods. Bayesian comparisons were based on the default t test with a Cauchy (0, r = 1/2^0.5^) prior implemented in JASP software (https://jasp-stats.org).

## Ethics declaration

This study involves an analysis of simulated fMRI data and open-access fMRI datasets. The HCP data were acquired using protocols approved by the Washington University institutional review board. The CNP procedures were approved by the Institutional Review Boards at UCLA and the Los Angeles County Department of Mental Health.

## Data availability

The HCP data are available at https://www.humanconnectome.org/study/hcp-young-adult/document/1200-subjects-data-release. The CNP data are available at https://openfmri.org/dataset/ds000030. Simulated data are available at https://github.com/IHB-IBR-department/TMFC_simulations.

## Code availability

Python and MATLAB code for task-modulated functional connectivity simulations, along with user-friendly Jupyter notebooks, are available at https://github.com/IHB-IBR-department/TMFC_simulations. Python module for TMFC simulations is available at https://github.com/IHB-IBR-department/TMFC_simulator. The SPM12-based TMFC toolbox with graphical user interface (GUI) for task-modulated functional connectivity analysis is available at https://github.com/IHB-IBR-department/TMFC_toolbox. MATLAB and Python functions for performing deconvolution based on ridge regression (https://github.com/IHB-IBR-department/BOLD_deconvolution).

## Acknowledgements

The current study was supported by the program of fundamental studies of IHB RAS (theme number 122041500046-5). Data were provided [in part] by the Human Connectome Project, WU-Minn Consortium (Principal Investigators: David Van Essen and Kamil Ugurbil; 1U54MH091657) funded by the 16 NIH Institutes and Centers that support the NIH Blueprint for Neuroscience Research and by the McDonnell Center for Systems Neuroscience at Washington University. Another part of the data was provided by the Consortium for Neuropsychiatric Phenomics (NIH Roadmap for Medical Research grants UL1-DE019580, RL1MH083268, RL1MH083269, RL1DA024853, RL1MH083270, RL1LM009833, PL1MH083271, and PL1NS062410).

## Author contributions

RM: Conceptualisation, Methodology, Software, Formal analysis, Writing – original draft, review & editing, Visualisation. IK: Methodology, Software, Formal analysis, Writing – review & editing, Visualisation. AK: Writing – review & editing. DC: Writing – review & editing, Funding acquisition. MK: Conceptualisation, Resources, Writing – review & editing, Supervision, Project administration, Funding acquisition.

## Competing interests

The authors declare no competing interests.

## Notes

https://github.com/IHB-IBR-department/TMFC_simulations

### Competing Interest Statement

The authors have declared no competing interest.

### Summary of Updates

Major revisions: 1) Performed new simulations with variable haemodynamic response parameters; 2) Added a new TMFC method to the comparison, namely beta-series correlation based on fractional ridge regression (Prince et al., 2022); 3) Considered the influence of mean centering of the task design regressors on the estimation of the sPPI/gPPI matrices; 4) Analysed the influence of amplitude differences on the asymmetry of gPPI matrices; 5) Provided software to implement deconvolution outside of SPM12 (MATLAB and Python code for deconvolution based on ridge regression); 6) Performed new simulations using task designs from HCP and CNP datasets; 7) Added new illustrations that show the drawbacks of the cPPI method; 8) Expanded discussion sections related to the task design parameters and limitations.

