## Supplementary for "Comparison of whole-brain task-modulated functional connectivity methods for fMRI task connectomics"

##### Supplementary Information 1: Sensitivity and specificity

For a thresholded $N \times N$ binary matrix $\boldsymbol{P}$, significant edges were coded as 1 and all other edges were coded as 0. For a ground-truth $N \times N$ binary matrix $\boldsymbol{T}$, the ground-truth edges were coded as 1 and all other edges were coded as 0. The ground-truth edges connected: (1) modules 1 and 2, 3 and 4 for "Condition A"; (2) modules 1 and 4, 2 and 3 for "Condition B". The sensitivity and specificity of task-modulated functional connectivity (TMFC) methods were computed as:

$$False\ Positives\ (FP) = \sum_{i=1}^{N}\sum_{j=1}^{N}((1 - T_{ij}) \times P_{ij}) \tag{S1.1}$$

$$False\ Negatives\ (FN) = \sum_{i=1}^{N}\sum_{j=1}^{N}(T_{ij} \times (1 - P_{ij})) \tag{S1.2}$$

$$True\ Positives\ (TP) = \sum_{i=1}^{N}\sum_{j=1}^{N}(T_{ij} \times P_{ij}) \tag{S1.3}$$

$$True\ Negatives\ (TN) = \sum_{i=1}^{N}\sum_{j=1}^{N}((1 - T_{ij}) \times (1 - P_{ij})) \tag{S1.4}$$

$$Sensitivity = True\ Positive\ Rate\ (TPR) = \frac{TP}{TP+FN} \times 100 \tag{S1.5}$$

$$Specificity = True\ Negative\ Rate\ (TNR) = \frac{TN}{TN+FP} \times 100 \tag{S1.6}$$

#### Supplementary Information 2: Empirical data

##### 2.1. Scanning parameters

For the Human Connectome Project (HCP) dataset, the sample size was N = 100 (unrelated healthy subjects, 54 females, 46 males, mean age = 29.1 ± 3.7). For the Consortium for Neuropsychiatric Phenomics (CNP) dataset, the sample size was N = 115 subjects (healthy subjects, 55 females, 60 males, mean age = 31.7 ± 8.9).

The fMRI data from the HCP dataset were collected using an echo-planar imaging (EPI) sequence on a modified 3T Siemens Skyra with the scanner repetition time, TR = 720 ms, echo time, TE = 33.1 ms, flip angle = 52°, in-plane field of view, FOV = 208 × 180 mm, 72 axial slices, and voxel size = 2 × 2 × 2 mm, with a multi-band acceleration factor of 8 (Glasser et al., 2013). Each HCP task consisted of two sessions: one with right-to-left and the other with left-to-right phase encoding (see duration in section 2.2). The HCP resting-state data consisted of two sessions each lasting ≈ 14.5 min (1200 scans).

The fMRI data from the CNP dataset were collected using an EPI sequence on a 3T Siemens Trio scanner with TR = 2 s, TE = 30 ms, flip angle = 90°, in-plane FOV = 192 × 192 mm, 34 axial slices, slice thickness = 4 mm, and voxel size = 3 × 3 × 4 mm (Poldrack et al., 2016). Each CNP task consisted of one session (see duration in section 2.2). The CNP resting-state data consisted of one session lasting ≈ 5 min (152 scans).

##### 2.2. Task designs

The block design tasks included working memory and social cognition tasks (for more details, see Barch et al., 2013). In the working memory task, each session consisted of eight blocks for two working memory loads (0-back and 2-back) and four stimulus categories (faces, places, tools, body parts). Each block lasted 27.5 s. For each working memory load within a session, two regressors were included in the subject-level model. Each session contained four fixation blocks (15 s). The total scanning time was ≈ 10 min (810 scans). In the social cognition task, each session consisted of two or three blocks for the random movement condition and two or three blocks for the social interaction condition. Each block lasted 20 s. For each condition within a session, two regressors were included in the subject-level model. Each session contained five fixation blocks (15 s). The total scanning time was ≈ 7 min (548 scans).

The event-related tasks were the stop-signal and task-switching tasks (for more details, see Poldrack et al., 2016). The stop-signal task consisted of 96 Go trials and 32 Stop-signal trials. All trials were preceded by a 500 ms fixation cross. Each trial lasted for 1000 ms and was separated by jittered null events ranging from 0.5 to 4 s (mean of 1 s). Two regressors were included in the subject-level model with a fixed event duration of 1.5 s. The total scanning time was ≈ 7 min (184 scans). The task-switching task consisted of 36 "Congruent, No Switch" trials, 12 "Congruent, Switch" trials, 36 "Incongruent, No Switch" trials, and 12 "Incongruent, Switch" trials. Four regressors were included in the subject-level model with a fixed duration of 1 s. The total scanning time was ≈ 6 min (208 scans).

#### Supplementary Information 3: Simulations

#### 3.1. Large-scale neural mass model

Consistent with previous studies (e.g., Papadopoulos et al., 2020), the coarse-grained form of the Wilson-Cowan equations for the *i*-th brain region can be expressed as:

$$\tau_E \frac{dE_i(t)}{dt} = -E_i(t) + \big(1 - E_i(t)\big) f_E\big[w_{EE}E_i(t) - w_{IE}I_i(t) + G\sum_j w_{ji}\, E_j(t-d) + P_E\big] + \xi_i^{ou}(t), \quad \text{(S2.1)}$$

$$\tau_I \frac{dI_i(t)}{dt} = -I_i(t) + \big(1 - I_i(t)\big) f_I[w_{EI}E_i(t) - w_{II}I_i(t) + P_I] + \xi_i^{ou}(t). \quad \text{(S2.2)}$$

The variables $E_i(t)$ and $I_i(t)$ correspond to neuronal activity of the excitatory and inhibitory subpopulations of the *i*-th region (proportion of excitatory and inhibitory cells firing per unit time), $\tau_E$ and $\tau_I$ are the excitatory and inhibitory time constants, and $w_{EE}$, $w_{IE}$, $w_{EI}$ and $w_{II}$ are the connectivity coefficients between excitatory and inhibitory subpopulations representing the average number of excitatory and inhibitory synapses per cell (synaptic weights). The non-linear response functions $f_E$ and $f_I$ for excitatory and inhibitory subpopulations are defined as:

$$f_E(x) = \frac{c_E}{1+e^{-a_E(x-b_E)}}, \quad \text{(S3.1)}$$

$$f_I(x) = \frac{c_I}{1+e^{-a_I(x-b_I)}}, \quad \text{(S3.2)}$$

where $a_E$ and $a_I$ are the slopes, $b_E$ and $b_I$ are the positions of the maximum slope, $c_E$ and $c_I$ are the amplitudes of excitatory and inhibitory response functions.

The $G\sum_j w_{ji}\, E_j(t-d)$ term represents interactions of the *i*-th region with the rest of the network, $G$ is the global coupling parameter, $w_{ji}$ is the synaptic weight from the *j*-th to *i*-th region, and $d$ is the signal transmission delay between brain regions. The delay was $d = 25$ ms, which is a physiologically plausible value for humans as estimated by Ringo et al. (1994). Synaptic weights ($w_{ji}$) were changed according to stimulus onsets. Typically, short-term synaptic plasticity reaches a plateau by the $8^{th}$ – $10^{th}$ excitatory postsynaptic current ($EPSC_{8\text{-}10}$), so at 40 Hz activity, it reaches a plateau at about 200 ms (Dittman et al., 2000; Kreitzer & Regehr, 2000). To account for the fact that synaptic weights do not change instantaneously, we added a synaptic plasticity delay of 200 ms between stimulus onset and synaptic weight change.

The $P_E$ and $P_I$ terms are constant, background drive for excitatory and inhibitory subpopulations. The $\xi_i^{ou}$ term is the background noise modelled as an Ornstein-Uhlenbeck process with zero mean (Uhlenbeck & Ornstein, 1930; Cakan et al., 2021, 2022):

$$\frac{d\xi_i^{ou}(t)}{dt} = -\frac{\xi_i^{ou}(t)}{\tau_{ou}} + \sigma_{ou}\xi_i(t), \quad \text{(S4)}$$

where $\tau_{ou}$ is the time scale, $\sigma_{ou}$ is the standard deviation of the process, and $\xi_i$ is the white Gaussian noise. In the absence of background noise, the Wilson-Cowan units produce unmodulated limit-cycle oscillations. With the addition of background noise, ultra-slow fluctuations in the power of the limit-cycle oscillations can be observed in accordance with empirical observations (Leopold et al., 2003; Nir et al., 2008; Keller et al., 2013; Thompson et al., 2013).

Before converting neuronal activity into the BOLD signal, we calculated synaptic activity as the sum of all inputs to the excitatory and inhibitory subpopulations (Tagamets & Horwitz, 1998; Kim & Horwitz, 2007; Ulloa & Horwitz, 2016):

$$SA_i(t) = w_{EE}E_i(t) + w_{EI}E_i(t) + w_{II}I_i(t) + w_{IE}I_i(t) + \sum_j w_{ji}\, E_j(t). \quad \text{(S5)}$$

Synaptic activity $SA_i(t)$ was considered as a proxy for local field potential (LFP) and used as input to the Balloon-Windkessel model.

All fixed parameters of the Wilson-Cowan model were selected as in Papadopoulos et al. (2020). Three tuning parameters ($G$, $P_E$, $\sigma_{ou}$) were determined based on the maximum similarity between the ground-truth synaptic weight matrix and the task-modulated functional connectivity (TMFC) matrix estimated using the direct correlation difference (CorrDiff) approach for block design time series without scanner measurement error. Similarity between matrices was assessed using Pearson's r correlation. First, we considered combinations of parameters with large steps: $G$ = 2.0 – 3.0 (step = 0.1), $P_E$ = 0.70 – 0.80 (step = 0.1), $\sigma_{ou} = 1.0\times10^{-3} - 6.0\times10^{-3}$ (step = $0.5\times10^{-3}$). Then we used smaller steps: $G$ = 2.60 – 2.75 (step = 0.01), $P_E$ = 0.755 – 0.765 (step = 0.01), $\sigma_{ou} = 3.0\times10^{-3} - 4.5\times10^{-3}$ (step = $0.1\times10^{-3}$). The selected set of tuning parameters for all simulations was as follows: $G = 2.63$, $P_E = 0.758$, $\sigma_{ou} = 3.5\times10^{-3}$. For the full set of parameters, see Supplementary Table S9.

Numerical integration of the system of ordinary differential equations was performed using the Euler-Maruyama method with a time step of $dt$ = 0.1 ms, implemented in the *neurolib* software (https://github.com/neurolib-dev/neurolib). *Neurolib* is a Python library that provides a computationally efficient framework for whole-brain resting-state functional connectivity

(RSFC) simulations (Cakan et al., 2021). In the current study, we modified the *neurolib* software to perform task-modulated functional connectivity (TMFC) simulations for a given task design and temporal resolution. To facilitate replication of our TMFC simulations, we share the simulated time-series, as well as MATLAB (R2021b) and Python code (versions 3.7-3.10) and user-friendly Jupyter notebooks (https://github.com/IHB-IBR-department/TMFC_simulations). The code for TMFC simulation is also available as a separate Python module *TMFC_simulator* (https://github.com/IHB-IBR-department/TMFC_simulator).

#### 3.2. The Balloon-Windkessel haemodynamic model

This model consists of three parts (Friston et al., 2003). The first describes relationship between synaptic activity and regional cerebral blood flow (rCBF). It is assumed that changes in synaptic activity cause an exponentially decaying vasodilatory signal $S_i(t)$, which is subject to autoregulatory feedback depending on the blood flow $f_i(t)$ it induces:

$$\frac{dS_i(t)}{dt} = SA_i(t) - \kappa S_i(t) - \gamma(f_i(t) - 1), \quad \text{(S6.1)}$$

$$\frac{df_i(t)}{dt} = S_i(t), \quad \text{(S6.2)}$$

where $\kappa$ is the rate of vasodilatory signal decay, and $\gamma$ is the rate of flow-dependent autoregulatory feedback.

The second part describes how change in blood flow causes changes in blood volume $v_i(t)$ and deoxyhaemoglobin content $q_i(t)$ within the post-capillary venous compartment, analogous to an expandable “balloon” or “windkessel" (air chamber):

$$\tau\frac{dv_i(t)}{dt} = f_i(t) - v_i^{1/\alpha}(t), \quad \text{(S7.1)}$$

$$\tau\frac{dq_i(t)}{dt} = f_i(t)\frac{1-(1-\rho)^{1/f_i(t)}}{\rho} - v_i^{1/\alpha}(t)\frac{q_i(t)}{v_i(t)}, \quad \text{(S7.2)}$$

where $\tau$ is the haemodynamic transit time (mean transit time of venous blood), $\alpha$ is the Grubb’s exponent (stiffness of the venous “balloon”), and $\rho$ is the resting net oxygen extraction fraction. Blood flow, blood volume and deoxyhaemoglobin content are expressed in normalised form relative to resting values.

The third part derives the BOLD signal $Y_i(t)$ from blood volume and deoxyhaemoglobin content by a static nonlinear function:

$$Y_i(t) = V_0\left[k_1\big(1 - q_i(t)\big) + k_2\left(1 - \frac{q_i(t)}{v_i(t)}\right) + k_3\big(1 - v_i(t)\big)\right], \quad \text{(S8)}$$

where $V_0$ is the resting volume fraction, $k_1 = 7\rho$, $k_2 = 2$, and $k_3 = 2\rho - 0.2$. The first term of Eq. (S8) describes the intrinsic extravascular signal, the second term describes the intravascular signal, and the third term describes the effect of changing the balance between them (Buxton et al., 1998).

#### Supplementary Information 4: Psycho-physiological interactions

The psychophysiological interaction (PPI) approach adapted for the ROI-to-ROI analysis is based on a general linear model of the following form:

$$Y_i = \beta_{PSY_i} X_{PSY} + \beta_{PHYS_{ij}} X_{PHYS_j} + \beta_{PPI_{ij}} X_{PPI_j} + \beta_{nuis_i} X_{nuis} + \beta_{0_i} + \varepsilon_i, \quad \text{(S9)}$$

where $Y_i$ is the BOLD signal in the target region ($i$-th ROI); $X_{PSY}$ is the psychological variable(s), modelling task-evoked haemodynamic responses; $X_{PHYS_j}$ is the physiological variable, BOLD signal in the "seed" region ($j$-th ROI); $X_{PPI_j}$ is the psychophysiological interaction variable(s); $X_{nuis}$ are the non-neuronal nuisance variables (e.g. head motion regressors and aCompCorr regressors); $\beta_{0_i}$ is the constant term; and $\varepsilon_i$ is the residual error. The beta values $\beta_{PSY_i}$, $\beta_{PHYS_{ij}}$, $\beta_{PPI_{ij}}$, and $\beta_{nuis_i}$ reflect the contributions to the target ROI of co-activations, spontaneous fluctuations, task-modulated fluctuations (TMFC) and non-neuronal noise, respectively.

The PPI term represents the element-by-element product of psychological and physiological variables. According to the standard PPI (sPPI) approach proposed by Friston et al. (1997), the general linear model includes one psychological and one PPI term for a difference between task conditions:

$$X_{PSY} = X_{PSY(A-B)} = (Z_{TASK(A)} - Z_{TASK(B)}) \otimes \text{HRF}, \quad \text{(S10.1)}$$

$$X_{PPI_j} = X_{PPI(A-B)_j} = X_{PHYS_j} \times ((Z_{TASK(A)} - Z_{TASK(B)}) \otimes \text{HRF}), \quad \text{(S10.2)}$$

where $Z_{TASK(A)}$ and $Z_{TASK(B)}$ are the task design regressors (box-car or delta functions) for conditions A and B, respectively; $\otimes$ is the convolution operator; and HRF is the haemodynamic response function.

According to the generalised PPI (gPPI) approach proposed by McLaren et al. (2012), the general linear model includes multiple psychological and PPI terms for each task condition:

$$X_{PSY} = X_{PSY(A)} + X_{PSY(B)} = Z_{TASK(A)} \otimes \text{HRF} + Z_{TASK(B)} \otimes \text{HRF}, \quad \text{(S11.1)}$$

$$X_{PPI_j} = X_{PPI(A)_j} + X_{PPI(B)_j} = X_{PHYS_j} \times (Z_{TASK(A)} \otimes \text{HRF}) + X_{PHYS_j} \times (Z_{TASK(B)} \otimes \text{HRF}). \quad \text{(S11.2)}$$

To account for the fact that psycho-physiological interactions occur at the neuronal level, Gitelman et al. (2003) proposed to deconvolve the BOLD signal in the seed ROI and calculate the element-by-element product of the deconvolved physiological regressor $Z_{PHYS_j}$ (estimated neuronal activity) and the task design regressor $Z_{TASK}$ (i.e. psychological variable before convolution with HRF). The product is then convolved with HRF:

$$X_{PHYS_j} = Z_{PHYS_j} \otimes \text{HRF} + \varepsilon, \quad \text{(S12.1)}$$

$$X^{*}_{PPI_j} = (Z_{PHYS_j} \times Z_{TASK}) \otimes \text{HRF}. \quad \text{(S12.2)}$$

We calculated PPI terms both with and without the deconvolution step. For more details on the deconvolution procedure, see Supplementary Information 5.

If the deconvolution step is applied, it is necessary to centre the task design regressor $Z_{TASK}$ before multiplication with the deconvolved psychological regressor $Z_{PHYS_j}$. When the PPI term is calculated with deconvolution but without centring, the PPI approach produces spurious results that resemble RSFC rather than TMFC (Di et al., 2017). Mean centring of $Z_{TASK}$ is the default setting in SPM12 since revision r6556. It is important to caution that the gPPI toolbox (McLaren et al., 2012) (https://www.nitrc.org/projects/gppi), one of the most popular tools for PPI analysis, does not use centering. In the main text, we calculated all PPI terms with $Z_{TASK}$ mean centering. For more details on the mean centering of the task design regressors, see Supplementary Information 6.

The correlational PPI (cPPI) approach proposed by Fornito et al. (2012) is based on the partial correlation between PPI terms for $i$-th and $j$-th ROIs, $r_{X_{PPI_i}, X_{PPI_j} \cdot C}$, eliminating the variance associated with confounds $C = \{X_{PHYS_i}, X_{PHYS_j}, X_{PSY}, X_{nuis}\}$. The cPPI approach uses PPI terms for the difference between task conditions, $r_{X_{PPI(A-B)_i}, X_{PPI(A-B)_j} \cdot C}$, as well as sPPI.

All MATLAB scripts for different types of PPI analysis are available on GitHub (https://github.com/IHB-IBR-department/TMFC_simulations). We also provide a user-friendly SPM12-based toolbox with GUI and parallel computing capability for voxel-based and ROI-to-ROI gPPI analysis with deconvolution, called *TMFC_toolbox* (https://github.com/IHB-IBR-department/TMFC_toolbox).

#### Supplementary Information 5: Deconvolution of the BOLD signal

In the main text, we used deconvolution approach proposed by Gitelman et al. (2002) and implemented in SPM12 (*spm_peb_ppi.m* function). Let's express convolution from Eq. (S12.1) in matrix form:

$$X_{PHYS} = HZ_{PHYS} + \varepsilon, \tag{S13}$$

where $H$ is the HRF in Toeplitz matrix form.

We can assume that the observed BOLD signal represents a neuronal signal convolved with a canonical HRF with measurement noise $\varepsilon$. The goal of BOLD signal deconvolution is to estimate this underlying unknown neuronal activity. To do this, we can expand the underlying neuronal activity in terms of a cosine basis set, so that the estimation of the temporal time series is transformed into the frequency domain. We can rewrite the neuronal signal as a linear combination of cosine functions (discrete cosine series, $X_{DCS}$) multiplied by unknown $\beta$ parameters:

$$Z_{PHYS} = X_{DCS}\beta. \tag{S14}$$

We now need to estimate $\beta$ parameters of the linear deconvolution model:

$$X_{PHYS} = HX_{DCS}\beta + \varepsilon. \tag{S15}$$

We can estimate parameters $\beta$ using the ordinary least-squares (OLS) solution:

$$\hat{\beta}^{OLS} = (X_{DCS}^T H^T HX_{DCS})^{-1} X_{DCS}^T H^T X_{PHYS}, \tag{S16}$$

where $X_{DCS}^T H^T$ is a transpose of $HX_{DCS}$.

To recover the neuronal signal from $\beta$ parameters, we need to multiply them by unconvolved cosine basis set.

However, if deconvolution is performed with least squares estimation, parameter estimates will be highly variable due to the use of the full-rank basis set and multicollinearity. This is particularly so for the coefficients controlling high frequencies, since HRF attenuates them. Therefore, some type of regularisation or prior constraints should be introduced to suppress this confounding effect. Gitelman et al. (2002) proposed to use Parametric Empirical Bayes (PEB) approach with Gaussian priors. SPM12 deconvolution suppresses high frequency components in the signals. The reconvolved BOLD signal represents a smoother version of the original BOLD signal.

A practical disadvantage of the PEB approach is that it is difficult to implement outside of the SPM12 package. As an alternative to the PEB approach, we can use ridge regression (Gaudes et al., 2010), which is readily available in most software packages for data analysis. Ridge regression (Tikhonov regularisation or L2-norm regularisation) penalizes higher parameter estimates and prevents model overfitting to noise by introducing a regularisation parameter $\alpha$:

$$\hat{\beta}^{RR} = (X_{DCS}^T H^T HX_{DCS} + \alpha I)^{-1} X_{DCS}^T H^T X_{PHYS}, \tag{S17}$$

where $I$ is the identity matrix (Hoerl & Kennard, 1970; Tikhonov & Arsenin, 1977).

When $\alpha = 0$, the penalty term has no effect, and ridge regression reduces to a least squares estimation. The higher $\alpha$ the more regularisation is applied. Increasing regularisation parameter $\alpha$ results in a smoother estimated neuronal signal, while decreasing $\alpha$ leads to overfitting.

To ensure the reproducibility of our results, we developed MATLAB and Python functions for ridge regression deconvolution, independent of the SPM12 software (https://github.com/IHB-IBR-department/BOLD_deconvolution). A comparison of the results produced by these functions and the *spm_peb_ppi.m* function can be found on GitHub (https://github.com/IHB-IBR-department/TMFC_simulations/tree/main/deconvolution). Here, our goal was not to determine the best deconvolution method, but to reproduce as closely as possible the results obtained by the *spm_peb_ppi.m* function without SPM12.

Consider, for example, ridge regression implemented in Python for a simulated BOLD signal in a block design with ten 20 s blocks per condition and a signal-to-noise ratio (SNR) of 0.4. The correlation between neuronal activity estimated by SPM12 PEB and ridge regression deconvolution was r = 0.998 at $\alpha$ = 0.005 (see Supplementary Fig. S15, S16a-c). The correlation between convolved PPI terms calculated using SPM12 PEB and ridge regression at $\alpha$ = 0.005 was r = 0.936 (see Supplementary Fig. S16d). Similar results were obtained for the simulated event-related task and empirical block and event-related tasks (https://github.com/IHB-IBR-department/TMFC_simulations/tree/main/deconvolution/python). For a simulated event-related task with one hundred 1 s events per condition and ISI = 6 s, the correlation between convolved PPI terms was r = 0.963 at $\alpha$ = 0.005. For the empirical block design task (working memory task), the correlation between convolved PPI terms was r = 0.985 at $\alpha$ = 0.002. For the empirical event-related task (stop-signal task) the correlation between convolved PPI terms was r = 0.952 at $\alpha$ = 0.003. Similar results were also obtained using ridge regression implemented in MATLAB (https://github.com/IHB-IBR-department/TMFC_simulations/tree/main/deconvolution/matlab). Therefore, ridge regression allows us to obtain similar estimates of neuronal activity as SPM12 PEB with the regularisation parameter $\alpha$ = 0.002-0.005.

#### Supplementary Information 6: Mean centering of the task design regressor prior to PPI term calculation

Without deconvolution, TMFC estimates obtained using PPI analysis with or without mean centering will be equivalent. Di et al. (2017) demonstrated it with empirical data and provided mathematical proof of this. For convenience, we present the mathematical proof here.

Consider the general linear model for PPI analysis from Eq. (S9). Let's remove the subscripts denoting ROI indices and add superscripts "1" for the model with a centered psychological regressor $X^1_{PSY}$:

$$Y = \beta^1_{PSY}X^1_{PSY} + \beta^1_{PHYS}X_{PHYS} + \beta^1_{PPI}X^1_{PPI} + \beta^1_{nuis}X_{nuis} + \beta^1_o + \varepsilon. \tag{S18}$$

We can rewrite the PPI term calculated with mean centering and without deconvolution as:

$$X^1_{PPI} = X^1_{PSY}X_{PHYS}. \tag{S19}$$

Put Eq. (S19) into Eq. (S18):

$$Y = \beta^1_{PSY}X^1_{PSY} + \beta^1_{PHYS}X_{PHYS} + \beta^1_{PPI}X^1_{PSY}X_{PHYS} + \beta^1_{nuis}X_{nuis} + \beta^1_o + \varepsilon. \tag{S20}$$

For a model without mean centering of the psychological regressor, add superscripts "2":

$$Y = \beta^2_{PSY}X^2_{PSY} + \beta^2_{PHYS}X_{PHYS} + \beta^2_{PPI}X^2_{PSY}X_{PHYS} + \beta^2_{nuis}X_{nuis} + \beta^2_o + \varepsilon. \tag{S21}$$

The non-centered psychological variable $X^2_{PSY}$ can be expressed as:

$$X^2_{PSY} = X^1_{PSY} + c, \tag{S22}$$

where *c* is a constant value.

Put Eq. (S22) into Eq. (S21):

$$Y = \beta^2_{PSY}(X^1_{PSY} + c) + \beta^2_{PHYS}X_{PHYS} + \beta^2_{PPI}(X^1_{PSY} + c)X_{PHYS} + \beta^2_{nuis}X_{nuis} + \beta^2_o + \varepsilon. \tag{S23}$$

And match terms in Eq. (S23) and Eq. (S20):

$$Y = \beta^2_{PSY}X^1_{PSY} + (\beta^2_{PHYS} + c\beta^2_{PPI})X_{PHYS} + \beta^2_{PPI}X^1_{PSY}X_{PHYS} + \beta^2_{nuis}X_{nuis} + (\beta^2_o + c\beta^2_{PSY}) + \varepsilon. \tag{S24}$$

Comparing Eq. (S24) and Eq. (20), it can be seen that:

$$\beta^1_{PSY} = \beta^2_{PSY}, \tag{S25.1}$$

$$\beta^1_{PHYS} = (\beta^2_{PHYS} + c\beta^2_{PPI}), \tag{S25.2}$$

$$\beta^1_{PPI} = \beta^2_{PPI}, \tag{S25.3}$$

$$\beta^1_{nuis} = \beta^2_{nuis}, \tag{S25.4}$$

$$\beta^1_o = \beta^2_o + c\beta^2_{PSY}. \tag{S25.5}$$

Therefore, without deconvolution, the parameter estimates of the psychological, PPI and nuisance regressors are exactly the same with or without mean centering. For TMFC analysis, we use the parameter estimates of the PPI regressors (Eq. S25.3). The only parameter estimates that change are those for the physiological regressor and the constant term (Eqs. S25.2 and S25.5).

We also used simulations to demonstrate that PPI results without deconvolution are exactly the same with or without mean centering (see Supplementary Fig. S17a-b and S18a-b).

However, when we apply deconvolution, the PPI matrices calculated for each of the task conditions (i.e., "Condition vs. Baseline") become different with or without mean centering (Fig. S17c-d and S18c-d). FC estimates between nodes that exhibit high connectivity during rest periods (reflecting task-unrelated spontaneous fluctuations) are positive when we do not apply mean centering (Fig. S17d and S18d) and negative when we apply mean centering (Fig. S17c and S18c). Without deconvolution, the FC estimates for these functional modules are also negative, regardless of mean centering (Fig. S17a-b and S18a-b). Similar to Di et al. (2017), we can see in simulations that the PPI matrices calculated without deconvolution (with or without mean centering) are more similar to the PPI matrices calculated with deconvolution and mean centering.

When we use deconvolution, we cannot match the model terms with and without mean centering as in Eqs. (S25.1-5). If the psychological variable is non-centered with a constant component, the constant component will add a physiological variable to the PPI term (Di et al., 2017). After deconvolution and reconvolution, this physiological component is no longer exactly the same as the original physiological variable. Consider the PPI term without deconvolution and without centering:

$$\beta^2_{PPI}(X^1_{PSY} + c)X_{PHYS} = \beta^2_{PPI}X^1_{PPI} + c\beta^2_{PPI}X_{PHYS}. \tag{S26}$$

We can combine the terms $c\beta^2_{PPI}X_{PHYS}$ and $\beta^2_{PHYS}X_{PHYS}$, since both contain the same variable $X_{PHYS}$, to obtain Eq. (S25.2) and Eq. (S23.3).

Now, consider the PPI term with deconvolution and without centering:

$$\beta^2_{PPI}[((Z^1_{PSY} + c)Z_{PHYS})\otimes\text{HRF}] = \beta^2_{PPI}[(Z^1_{PSY}Z_{PHYS})\otimes\text{HRF}] + \beta^2_{PPI}[(cZ_{PHYS})\otimes\text{HRF}]. \tag{S27}$$

We cannot combine the terms $\beta^2_{PPI}[(cZ_{PHYS})\otimes\text{HRF}]$ and $\beta^2_{PHYS}X_{PHYS}$ and obtain equations similar to Eqs. (S25.1-5), since the original physiological signal is not equal to the physiological signal reconvolved from the estimated neuronal signal:

$$X_{PHYS} \neq Z_{PHYS} \otimes \text{HRF}. \tag{S28}$$

Meanwhile, the main goal of studying task-modulated functional connectivity is not to study connectivity only in one task condition but the relative connectivity changes in one task compared with others ("Condition A vs. Condition B"). For the balanced task designs, the sPPI and gPPI matrices showing the difference in FC between conditions are still the same with or without mean centering (Fig. S17c-d and S18c-d). In well-balanced designs, the effects related to spontaneous fluctuations are similar in the condition of interest and the control condition, so they canceled out for the "Condition A vs. Condition B" comparison. However, it may be not the case if the spontaneous fluctuation component has different weights for two task conditions, for example, the two task conditions have different numbers of blocks or events (Di et al., 2017).

To demonstrate this, we performed simulations with task designs identical to the stop-signal task and task-switching task from the CNP dataset, as an examples of unbalanced event-related designs. The stop-signal task consists of 96 Go trials and 32 Stop-signal trials. The task-switching task consists of 72 "No Switch" trials and 24 "Switch" trials. For both task designs, we can see a significant difference in FC between conditions within functional modules (i.e., FC associated with task-unrelated spontaneous fluctuations) when using the sPPI method without mean centering (see Supplementary Fig. S19). This is a false positive result since we did not change synaptic weights within functional modules during the task. This false positive result can be avoided if we apply mean centering to the sPPI method or use the gPPI method with or without mean centering.

#### Supplementary Information 7: Beta-series correlation

The beta-series correlation "Least-Squares All" (BSC-LSA) approach is based on a single general linear model with all trials represented by separate regressors (Rissman et al., 2004):

$$Y = \beta_{A1} X_{TASK(A1)} + \cdots + \beta_{AN} X_{TASK(AN)} + \beta_{B1} X_{TASK(B1)} + \cdots + \beta_{BN} X_{TASK(BN)} + \beta_{NOISE} X_{NOISE} + \beta_0 + \varepsilon \qquad \text{(S29)}$$

where $\{X_{TASK(A1)}, \ldots, X_{TASK(AN)}\}$ are regressors for individual trials {1, …, N} for task condition A; $\{X_{TASK(B1)}, \ldots, X_{TASK(BN)}\}$ are regressors for individual trials {1, …, N} for task condition B; $\{\beta_{A1}, \ldots, \beta_{AN}\}$ and $\{\beta_{B1}, \ldots, \beta_{BN}\}$ are beta parameters for individual trials for task conditions A and B, respectively. For rapid event-related results with short interstimulus intervals, the correlation between individual trial regressors becomes high (multicollinearity problem), resulting in noisy beta estimates obtained by the ordinary least squares.

The BSC "Least-Squares Separate" (BSC-LSS) approach is based on separate general linear models for each individual trial, avoiding multicollinearity problem (Mumford et al., 2012):

$$Y = \beta_{A1} X_{TASK(A1)} + \beta_{A\{2:N\}B\{1:N\}} X_{TASK(A\{2:N\}B\{1:N\})} + \beta_{NOISE} X_{NOISE} + \beta_0 + \varepsilon \qquad \text{(S30.1)}$$

…

$$Y = \beta_{AN} X_{TASK(AN)} + \beta_{A\{1:N-1\}B\{1:N\}} X_{TASK(A\{1:N-1\}B\{1:N\})} + \beta_{NOISE} X_{NOISE} + \beta_0 + \varepsilon \qquad \text{(S30.2)}$$

$$Y = \beta_{B1} X_{TASK(B1)} + \beta_{A\{1:N\}B\{2:N\}} X_{TASK(A\{1:N\}B\{2:N\})} + \beta_{NOISE} X_{NOISE} + \beta_0 + \varepsilon \qquad \text{(S30.3)}$$

…

$$Y = \beta_{BN} X_{TASK(BN)} + \beta_{A\{1:N\}B\{1:N-1\}} X_{TASK(A\{1:N\}B\{1:N-1\})} + \beta_{NOISE} X_{NOISE} + \beta_0 + \varepsilon \qquad \text{(S30.4)}$$

where $\{X_{TASK(A\{2:N\}B\{1:N\})}, \ldots, X_{TASK(A\{1:N-1\}B\{1:N\})}\}$ are regressors for all trials except for the {1, …, N}-th individual trial for task condition A and all trials for condition B; $\{X_{TASK(A\{1:N\}B\{2:N\})}, \ldots, X_{TASK(A\{1:N\}B\{1:N-1\})}\}$ are regressors for all trials for task condition A and all trials except for the {1, …, N}-th individual trial for condition B.

In this paper, we also propose to apply fractional ridge regression (FRR) for BSC analysis. FRR approach is based on the LSA model, but uses fractional ridge regression instead of least-squares estimation to regularize single-trial beta estimates for each voxel or region of interest (Rokem & Kay, 2020; Prince et al., 2022). FRR is a reparameterisation of the ridge regression in terms of the ratio $\gamma$ between the L2-norms of the regularised and unregularised parameters (Rokem & Kay, 2020):

$$\gamma = \frac{\left\| \tilde{\beta}^{RR} \right\|_2}{\left\| \tilde{\beta}^{OLS} \right\|_2}, \qquad \text{(S31)}$$

where $\left\| \tilde{\beta}^{RR} \right\|_2$ and $\left\| \tilde{\beta}^{OLS} \right\|_2$ are L2-norms of the regularised and unregularised (ordinary least squares) solutions in the low-dimensional space obtained by singular value decomposition (SVD) of the design matrix *X*. The fraction value $\gamma = 1$ means no regularisation, and $\gamma = 0$ means full regularisation (shrinking all parameters to $\beta = 0$).

One of the challenges of using ridge regression is the choice of the regularisation parameter $\alpha$. FRR is fast and scalable approach to determine the degree of regularisation that yields the best solution, designed to handle large-scale data. Computational redundancies are avoided because solutions with different fraction value $\gamma$ are guaranteed to be different. FRR approach explores the full range of effects of regularisation on $\beta$, thus ensuring that the best possible solution is within the range of solutions explored (Rokem & Kay, 2020).

The LSS approach can be thought as an extreme regularisation approach applied uniformly across all brain regions, while the FRR approach can be thought as a tunable regularisation where regularisation is applied only if the data need it. In the current work, the regularisation for each region of interest was determined via split-half cross-validation on single-trial betas. The first step of the cross-validation procedure is to analyze all data using the ordinary least squares ($\gamma = 1$). The second step is to perform a grid search over fraction value $\gamma$ ranging from 0.05 to 1.00 in increments of 0.05. For each fraction value $\gamma$, we assess how well the beta estimates calculated on one half (training) generalize to the other half (testing). Squared errors between the regularised beta estimates from the training half and the unregularised beta estimates from the validation half are computed. Next, this procedure is repeated for the swapped halves, and then the errors are summed up. Finally, the optimal fraction value $\gamma$ is applied to the full dataset.

To calculate regularised single-trial beta estimates, we used the FRR approach implemented in the GLMsingle toolbox for MATLAB (Prince et al., 2022) (https://glmsingle.org/). We used the canonical HRF for the BSC-FRR analysis, as in all other TMFC analyses, and did not use data-derived nuisance regressors (i.e., GLMdenoise). For the BSC-FRR analysis, we did not perform upsampling of the task design matrix prior to convolution with HRF, since the GLMsignle toolbox requires that the task design have the same temporal resolution as the data to be convolved.

In the current work, beta values for individual trials were calculated using LSA, LSS and FRR approaches. Pearson's r correlations were calculated between the mean beta values extracted from the *i*-th and *j*-th ROIs, separately for task conditions A and B ($\{\beta_{A1}, \ldots, \beta_{AN}\}$ and $\{\beta_{B1}, \ldots, \beta_{BN}\}$). When Pearson's r correlations for task conditions A and B were converted to Fisher's Z and subtracted.

All MATLAB scripts for different types of BSC analysis are available on GitHub (https://github.com/IHB-IBR-department/TMFC_simulations). We also provide a user-friendly SPM12-based toolbox with GUI and parallel computing capability for voxel-based and ROI-to-ROI BSC-LSS analysis, called *TMFC_toolbox* (https://github.com/IHB-IBR-department/TMFC_toolbox).

#### Supplementary Information 8: Regression dynamic causal modelling

Dynamic causal modelling (DCM) is a generative modelling framework for inferring effective connectivity from neuroimaging data (Friston et al., 2003). Originally, DCM was introduced for fMRI data and described changes in hidden neuronal states by a bilinear differential equation:

$$\frac{dx}{dt} = \left(A + \sum_{k=1}^{m} u_k B^{(k)}\right)x + Cu, \quad \text{(S32)}$$

where $x$ is the neural state, $u_k$ is the *k-th* experimental manipulation, $A$ is the matrix of task-independent (endogenous) connection strengths, $B^{(k)}$ is the matrix representing connection strengths modulated by *k-th* experimental manipulation (task-modulated or exogenous effective connectivity, TMEC), and $C$ is the matrix representing the direct influence of driving inputs (co-activations). Conceptually, the $\{A, B, C\}$ parameters can be matched with the $\{\beta_{PHYS}, \beta_{PPI}, \beta_{PSY}\}$ parameters from the PPI approach (see Eq. S9).

The combination of the neuronal state equation (Eq. S32) and the set of the Balloon-Windkessel haemodynamic model equations (Eqs. S6-8) constitutes a generative (forward) model. The model inversion is aimed at finding the parameters that enable the model to best explain the data and is based on Variational Bayes under the Laplace assumption (Friston et al., 2007). The problem is that the original DCM is limited to parameter estimation of small brain networks (about ten nodes), since the model inversion becomes ill-posed and computationally demanding for large-scale (whole-brain) networks. To address this problem, a regression DCM (rDCM) approach has been proposed (Frassle et al., 2017).

This approach applies several simplifications and modifications to the original DCM that include: (1) replacing the bilinear equation with a linear equation (removing the $\sum_{k=1}^{m} u_k B^{(k)} x$ term, related to TMEC); (2) translation from the time domain to the frequency domain using the Fourier transformation; (3) replacing the non-linear haemodynamic model (the Balloon-Windkessel model) with a fixed, linear haemodynamic response function (HRF); (4) ignoring dependencies among parameters affecting distinct brain regions; and (5) replacing the log-normal prior on noise variance with a Gaussian prior. These modifications reformulate the model inversion as a special case of Bayesian linear regression and significantly improve computational efficiency for large-scale networks (Frassle et al., 2017). We used the rDCM method implemented in the TAPAS software collection (Frassle et al., 2021).

#### Supplementary Information 9: Asymmetry of the regression coefficients due to amplitude differences

One possible reason for the PPI matrices asymmetry could be differences in the BOLD signal amplitudes, rather than causal directionality (Cole et al., 2016). When using linear regression to estimate the statistical relationship between two time series, the difference in time-series amplitudes typically makes the regression coefficients asymmetric with respect to the substitution of dependent and independent variables. This asymmetry does not reflect real causality between time series.

Here, we can demonstrate it with two simple "toy" examples (https://github.com/IHB-IBR-department/TMFC_simulations/tree/main/amplitude_difference_toy_examples). We generated correlated time series for 30 ROIs and 100 subjects from a multivariate normal distribution with a mean of 100 and a covariance matrix with main diagonal elements equal 0.5 and off-diagonal elements equal 0.1. Each time series consisted of 400 time points. We then calculated the dependencies between each pair of ROIs using Pearson's r correlation and linear regression. In this case, the connectivity matrices obtained by correlation and regression methods were similar (see Supplementary Fig. S11a). The regression-based connectivity matrix was close to symmetric, since the correlation between the upper and lower diagonal elements was very high, $r_{(up,down)}$ = 0.90. The amplitude differences were small relative to the mean amplitude of the time series (mean amplitude difference across ROIs and subjects, $\overline{amp}_{diff}$ = -0.0001). The correlation between the absolute amplitude differences and differences in beta coefficients from the upper and lower diagonal was close to zero (mean correlation across subjects, $\bar{r}_{(|amp_{diff}|,|\beta_{diff}|)}$ = -0.001).

Next, we split 30 ROIs on two "functional modules". The first 15 ROIs were assigned to Module #1 (M1), and the last 15 ROIs were assigned to Module #2 (M2). We considered two examples with amplitude differences.

In the first example, we added uncorrelated time series to the M1 (Fig. S11b). The uncorrelated time series were generated from a normal distribution with a mean of one and a standard deviation of 0.5. The mean amplitude difference across ROIs and subjects, $\overline{amp}_{diff}$ = 0.5, is relatively high compared to the mean amplitude of the time series of 4.7. In this example, we observed a clear difference between correlation-based and regression-based connectivity matrices. In the correlation-based matrix, we saw the largest connectivity decrease within M1, a medium decrease between M1 and M2, and no decrease within M2. We also observed a decrease within M1 and no decrease within M2 in the regression-based matrix. However, there was a decrease in connectivity estimates from M1 to M2 (M1→M2, M1 – seed, M2 – target) and no decrease from M2 to M1 (M2→M1, M2 – seed, M1 – target). This made the regression-based matrix noticeably asymmetric ($r_{(up,down)}$ = 0.33). The correlation between the absolute amplitude differences and differences in beta coefficients from the upper and lower diagonal was high, $\bar{r}_{(|amp_{diff}|,|\beta_{diff}|)}$ = 0.80. Cole et al. (2016) previously obtained a similar result showing that adding an unshared signal can decrease source (or seed) betas and not change target betas.

In the second example, we added correlated time series to the M1 (Fig. S11c). The correlated time series were generated from a multivariate normal distribution with a mean of one and a covariance matrix with main diagonal elements equal 0.5 and off-diagonal elements equal 0.15. In the correlation-based matrix, we observed a connectivity increase within M1, a decrease between M1 and M2, and no decrease within M2. In the regression-based matrix, we also observed an increase within M1 and no decrease within M2. However, as in the previous example, connectivity estimates decreased from M1 to M2 and did not decrease from M2 to M1. In this example, there was also a high correlation between the absolute amplitude differences and differences in beta coefficients from the upper and lower diagonal, $\bar{r}_{(|amp_{diff}|,|\beta_{diff}|)}$ = 0.82.

To summarize, introducing amplitude differences by adding uncorrelated and correlated signals into some functional module decreased the connectivity estimates from that module to other modules and did not change estimates from other modules to that module. Within-module connectivity estimates increased after adding correlated signals and decreased after adding uncorrelated signals.

In the large-scale neural mass simulations, the time series for all ROIs had the same amplitudes for the rest condition. However, there were two potential sources of differences in amplitudes during task conditions. The first was the change of synaptic weights, and the second was the addition of co-activations. We verified that the change of synaptic weight did not change the amplitude differences between ROIs. However, the addition of co-activations leaded to amplitude differences and asymmetry of PPI matrices (in particular, the sPPI matrices). After FIR task regression, amplitude differences and the asymmetry of PPI matrices associated with these differences were eliminated. Below we consider several simulations in which we independently manipulated synaptic weights and co-activations.

***(A) Simulation with task-related synaptic weight changes and without adding co-activations.***

For the block design with ten 20 s blocks per condition, the mean amplitude difference during task conditions across 100 ROIs and 100 subjects was $\overline{amp}_{diff}$ = 0.004 without adding co-activations, which is relatively small compared to the mean amplitude of 12.5. The gPPI matrix was close to symmetric, $r_{(up,down)}$ = 0.97. The correlations between the absolute amplitude differences during task conditions and absolute differences in beta coefficients from the upper and lower diagonal of the "Cond A-B" matrix were close to zero. For condition A, $\bar{r}_{(|amp_{diff,A}|,|\beta_{diff,A-B}|)}$ = -0.002. For condition B,

$\bar{r}_{(|amp_{diff,B}|,|\beta_{diff,A-B}|)}$ = -0.001. Therefore, the change of synaptic weight did not induce the amplitude differences between ROIs.

***(B) Simulation without task-related synaptic weight changes and with adding co-activations.***

In the previous case, we changed the synaptic weights and did not add co-activations. Now, we consider simulation without synaptic weights changes, but with adding co-activations (see Supplementary Fig. S12). Co-activations were added to modules 1 (M1) and 3 (M3) in condition A and to modules 2 (M2) and 4 (M4) in condition B (Fig. S12b). The mean amplitude difference during task conditions across 100 ROIs and 100 subjects was $\overline{amp}_{diff}$ = 0.6, which is comparable to the mean amplitude of 12.8. Adding co-activations is analogous to adding correlated signals, as in the second "toy" example described above (Fig. S11c). Therefore, we can expect an increase in beta parameters within and between M1 and M3 for condition A, as well as M2 and M4 for condition B. More importantly, we can expect a decrease in M1→M2, M1→M4, M3→M2, M3→M4 for condition A and decrease in M2→M1, M2→M3, M4→M1, M4→M3 for condition B (Fig. S12b). This should result in an asymmetric TMFC matrix ("Cond A-B"). Indeed, we can see the expected pattern of the asymmetry when using the sPPI method without FIR task regression (Fig. S12c). In this case, the asymmetry of regression coefficients was due differences in the BOLD signal amplitudes. The correlations between the absolute amplitude differences during task conditions and absolute differences in beta coefficients from the upper and lower diagonal of the "Cond A-B" matrix across subjects were high. For condition A, $\bar{r}_{(|amp_{diff,A}|,|\beta_{diff,A-B}|)}$ = 0.736. For condition B, $\bar{r}_{(|amp_{diff,B}|,|\beta_{diff,A-B}|)}$ = 0.737. The sPPI method failed to eliminate the effects related to co-activations. We observed significant FC differences between all regions after FDR thresholding, which were false positives. Meanwhile, when we used the gPPI method without FIR task regression, all false positives were eliminated (Fig. S12d). The asymmetry due to amplitude differences was also eliminated: $\bar{r}_{(|amp_{diff,A}|,|\beta_{diff,A-B}|)}$ = -0.004 and $\bar{r}_{(|amp_{diff,B}|,|\beta_{diff,A-B}|)}$ = -0.003. Furthermore, when we performed FIR task regression before TMFC analysis, the sPPI method did not produce false positives or demonstrate asymmetry due to amplitude differences: $\bar{r}_{(|amp_{diff,A}|,|\beta_{diff,A-B}|)}$ = -0.004, $\bar{r}_{(|amp_{diff,B}|,|\beta_{diff,A-B}|)}$ = -0.003 (Fig. S12e). The same was true for the gPPI method with FIR task regression: $\bar{r}_{(|amp_{diff,A}|,|\beta_{diff,A-B}|)}$ = -0.003, $\bar{r}_{(|amp_{diff,B}|,|\beta_{diff,A-B}|)}$ = -0.002 (Fig. S12f). Therefore, the gPPI method (with or without FIR task regression) as well as the sPPI method with FIR task regression protects against co-activation effects and regression coefficient asymmetry due to amplitude differences.

***(C) Simulations with task-related synaptic weight changes and co-activations.***

All simulations presented in the main text were performed with the change of synaptic weights during task conditions and addition of co-activations, unless otherwise stated. To deal with co-activations, we performed FIR task regression before TMFC analysis. Thus, the asymmetry of the regression coefficients due to the amplitude differences was eliminated. For the block design simulation with task-related synaptic weight changes and co-activations, the correlations between the absolute amplitude differences during task conditions and absolute differences in beta coefficients from the upper and lower diagonal of the "Cond A-B" matrix were close to zero: $\bar{r}_{(|amp_{diff,A}|,|\beta_{diff,A-B}|)}$ = 0.003 and $\bar{r}_{(|amp_{diff,B}|,|\beta_{diff,A-B}|)}$ = -0.009.

***(D) Simulations with asymmetric ground-truth synaptic weights and without co-activations***

In the last section of the main text, we considered simulations with asymmetric ground-truth matrices (see Supplementary Fig. S10) without co-activations to test whether the gPPI method could *in principle* reveal true causal directionality. Here, we consider the best-case scenario: block design simulation with twenty 20 s blocks per condition (twice as much as in the default block design), high SNR = 5 and no HRF variability. The gPPI method was used with deconvolution and without a symmetrisation procedure.

The mean amplitude difference during task conditions across 100 ROIs and 100 subjects was $\overline{amp}_{diff}$ = 0.002 without adding co-activations, which is relatively small compared to the mean amplitude of 4.9. The correlations between the absolute amplitude differences during task conditions and absolute differences in beta coefficients from the upper and lower diagonal of the "Cond A-B" matrix were close to zero. For condition A, $\bar{r}_{(|amp_{diff,A}|,|\beta_{diff,A-B}|)}$ = 0.051. For condition B, $\bar{r}_{(|amp_{diff,B}|,|\beta_{diff,A-B}|)}$ = 0.039. Therefore, the asymmetry of the gPPI matrix was not related to the amplitude differences between ROIs. The gPPI asymmetry reflected the asymmetric ground-truth synaptic weights. The correlation between the the group-mean asymmetric gPPI matrix and asymmetric ground-truth "Cond A-B" matrix was 0.93. The percent of correct direction estimated (correct sign rate, CSR) was 100%. CSR was defined as the ratio between correctly identified sign of connections to the total number of non-zero ground-truth connections for the "Cond A-B" contrast:

$$Correct\ Sign\ Rate\ (CSR) = \frac{\sum_{i=1}^{N}\sum_{j=1}^{N}((|T_{ij}^{Asymm} \times S_{ij}| + (T_{ij}^{Asymm} \times S_{ij}))/2)}{\sum_{i=1}^{N}\sum_{j=1}^{N}|T_{ij}^{Asymm}|}, \quad (S33)$$

where $S_{ij}$ is a signed $N\times N$ matrix representing the sign of the difference between the connectivity estimates for conditions A and B ("Cond A-B" contrast). $T_{ij}^{Asymm}$ is a $N\times N$ signed matrix representing the asymmetric ground-truth signs for the

“Cond A-B” contrast. In “Cond A”, synaptic weights were increased for M4→M3, M3→M2, M2→M1 and M1→M4 (coded as 1 in $T_{ij}^{Asymm}$). In “Cond B”, synaptic weights were increased in the opposite direction M1→M2, M2→M3, M3→M4 and M4→M1 (coded as -1 in $T_{ij}^{Asymm}$). All other connections were coded as 0 in the $T_{ij}^{Asymm}$ matrix (see Fig. 10a in the main text). CSR of 50% means the sign was detected by chance. CSR of 100% means that the signs of all connections presented in the ground truth were correctly identified. CSR of 0% means the opposite of 100% (all signs are flipped).

Thus, the gPPI method with deconvolution was able to reveal the true causal directionality in the best-case scenario. However, there are two important notes to make. First, even though CSR = 100%, none of the connections survived the FDR-corrected threshold of 0.001. Second, when we shortened the scan duration, reduced the SNR, and, most importantly, introduced HRF variability, the ability of the gPPI method to correctly identify the direction of information flow was reduced to almost zero.

### Supplementary Figures

#### Supplementary Figure S1

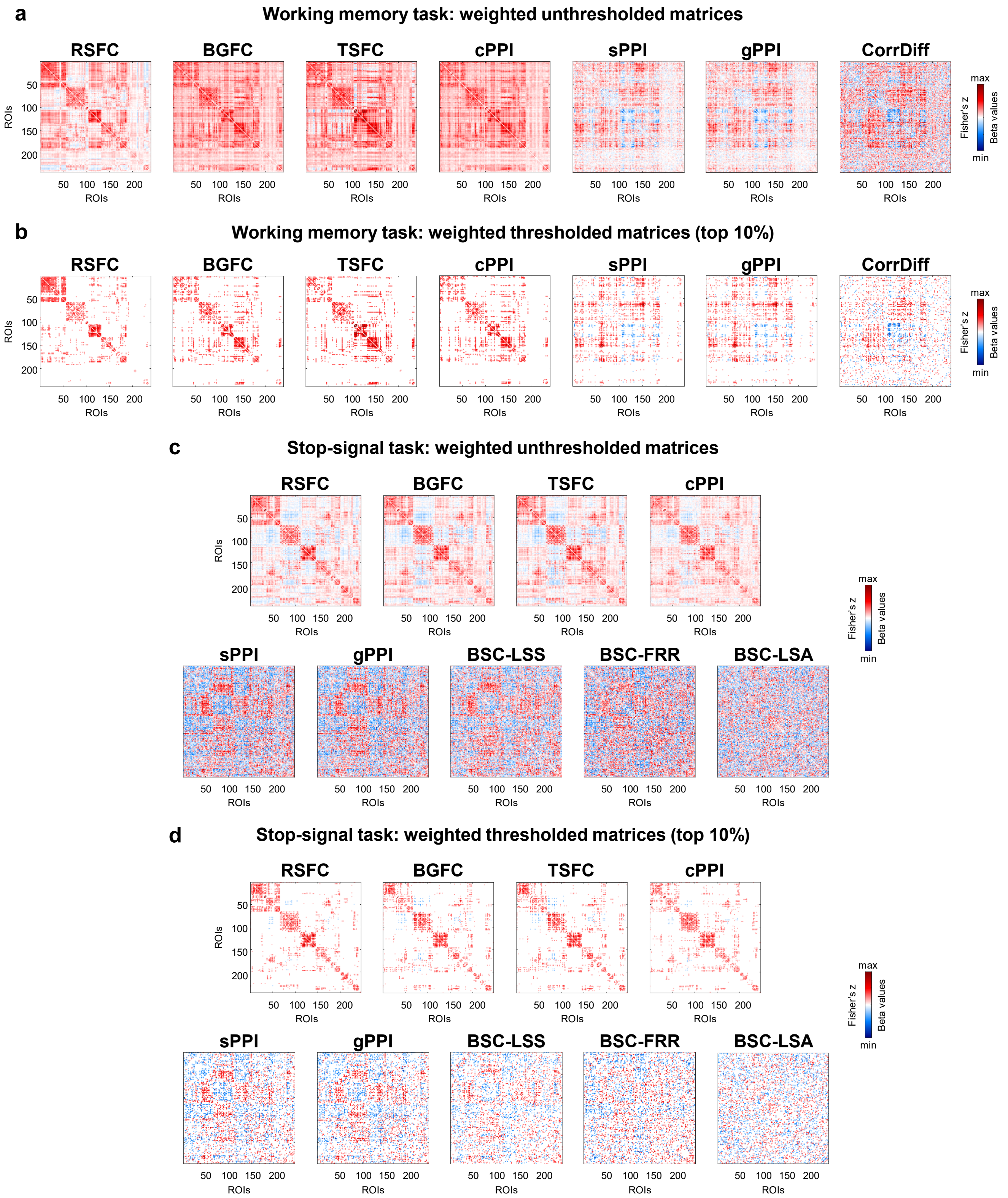


**Figure S1. RSFC, BGFC and TSFC matrices as well as FC matrices obtained by different TMFC methods.** sPPI and gPPI matrices were symmetrised. All PPI terms were calculated with the deconvolution step. **(a, b)** Results for the block design: working memory task ("2-back > 0-back"). **(c, d)** Results for the event-related design: stop-signal task ("Stop > Go"). **(a, c)** Unthresholded weighted group-mean matrices. **(b, d)** Thresholded weighted group-mean matrices representing the top 10% connections.

**Supplementary Figure S2**

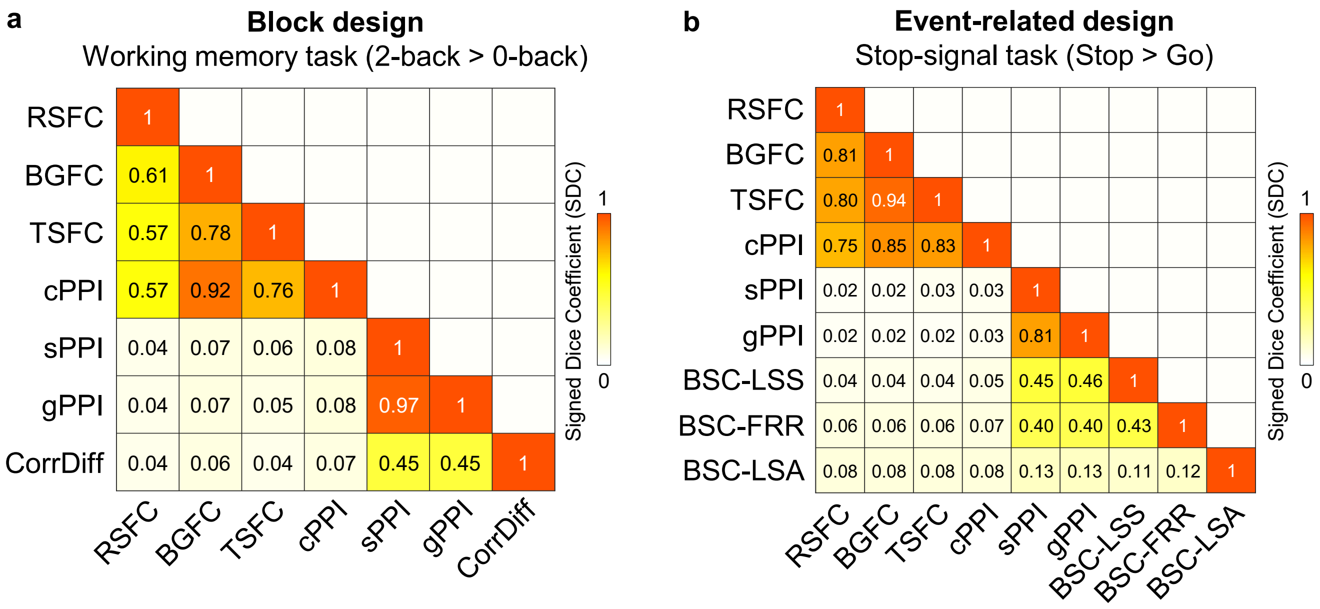


**Figure S2. Overlap between thresholded RSFC, BGFC and TSFC matrices and FC matrices obtained by different TMFC methods.** sPPI and gPPI matrices were symmetrised. All PPI terms were calculated with the deconvolution step. **(a)** Results for the block design: working memory task. **(b)** Results for the event-related design: stop-signal task. The overlap was calculated between thresholded weighted group-mean matrices representing the top 10% connections. Overlap was defined as the ratio between overlapping connections with the same sign to the number of all connections in the two thresholded weighted (signed) matrices. We refer to this index as the Signed Dice Coefficient (SDC), which is similar to the original Dice Coefficient but takes into account the sign of connections.[1] SDC of 0 means no overlap between thresholded signed matrices. SDC of 1 means complete overlap between positive and negative connections of two thresholded signed matrices.

---

[1] Dice Coefficient can be defined as:

$$Dice\ Coefficient\ (DC) = \frac{2\sum_{i=1}^{N}\sum_{j=1}^{N}|W_{ij}^{(1)} \times W_{ij}^{(2)}|}{\sum_{i=1}^{N}\sum_{j=1}^{N}|W_{ij}^{(1)}| + \sum_{i=1}^{N}\sum_{j=1}^{N}|W_{ij}^{(2)}|},$$

where $W_{ij}^{(1)}$ and $W_{ij}^{(2)}$ are thresholded signed $N \times N$ matrices (consisting of 1, 0, -1) calculated by method (1) and method (2). When the Signed Dice Coefficient can be defined as:

$$Signed\ Dice\ Coefficient\ (SDC) = \frac{2\sum_{i=1}^{N}\sum_{j=1}^{N}((|W_{ij}^{(1)} \times W_{ij}^{(2)}| + (W_{ij}^{(1)} \times W_{ij}^{(2)}))/2)}{\sum_{i=1}^{N}\sum_{j=1}^{N}|W_{ij}^{(1)}| + \sum_{i=1}^{N}\sum_{j=1}^{N}|W_{ij}^{(2)}|}.$$

#### Supplementary Figure S3

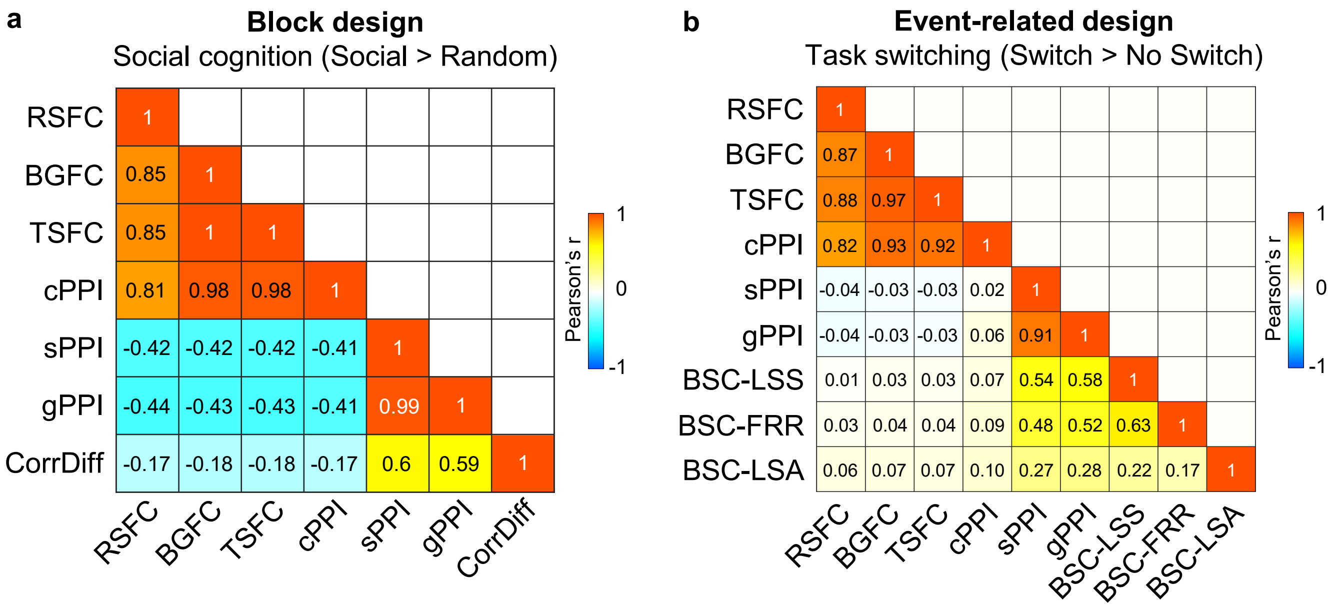


**Figure S3. Correlations between unthresholded RSFC, BGFC and TSFC matrices and FC matrices obtained by different TMFC methods.** To evaluate the similarity between the raw FC matrices, we calculated Pearson's r correlations between lower diagonal elements. sPPI and gPPI matrices were symmetrised. All PPI terms were calculated with the deconvolution step. **(a)** Results for the block design: social cognition task. **(b)** Results for the event-related design: task-switching task.

**Supplementary Figure S4**

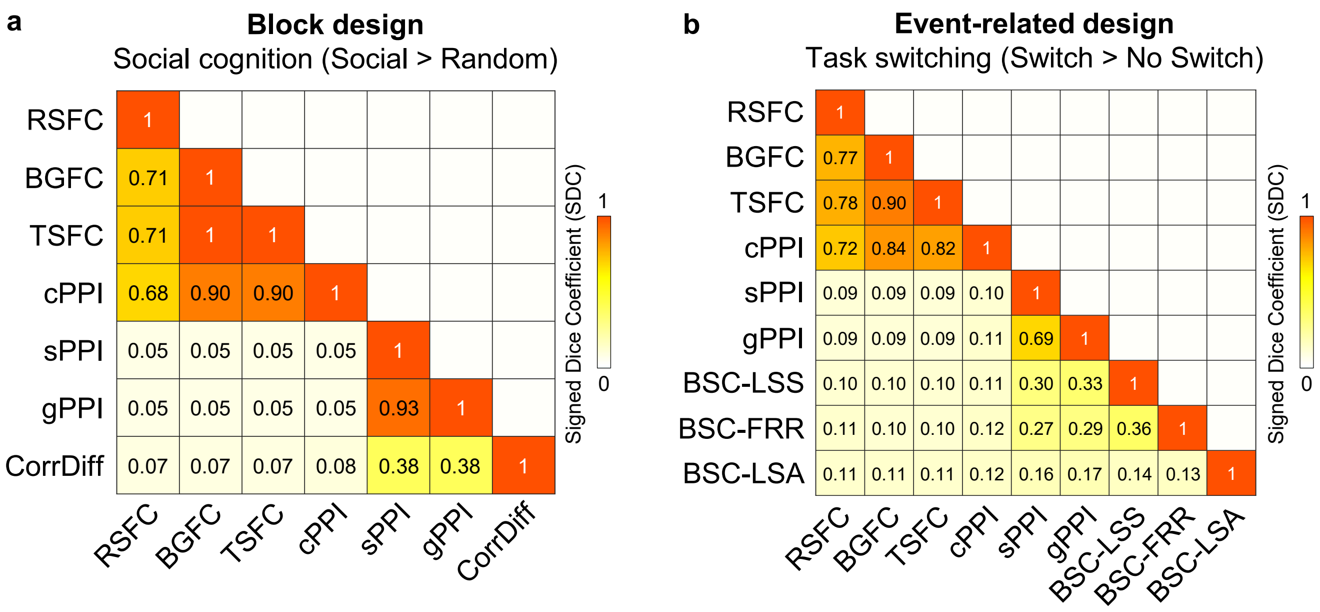


**Figure S4. Overlap between thresholded RSFC, BGFC and TSFC matrices and FC matrices obtained by different TMFC methods.** sPPI and gPPI matrices were symmetrised. All PPI terms were calculated with the deconvolution step. **(a)** Results for the block design: social cognition task. **(b)** Results for the event-related design: task-switching task. The overlap was calculated between thresholded weighted group-mean matrices representing the top 10% connections. Overlap was defined as the ratio between overlapping connections with the same sign to the number of all connections in the two thresholded weighted (signed) matrices. We refer to this index as the Signed Dice Coefficient (SDC), which is similar to the original Dice Coefficient but takes into account the sign of connections. SDC of 0 means no overlap between thresholded signed matrices. SDC of 1 means complete overlap between positive and negative connections of two thresholded signed matrices.

#### Supplementary Figure S5

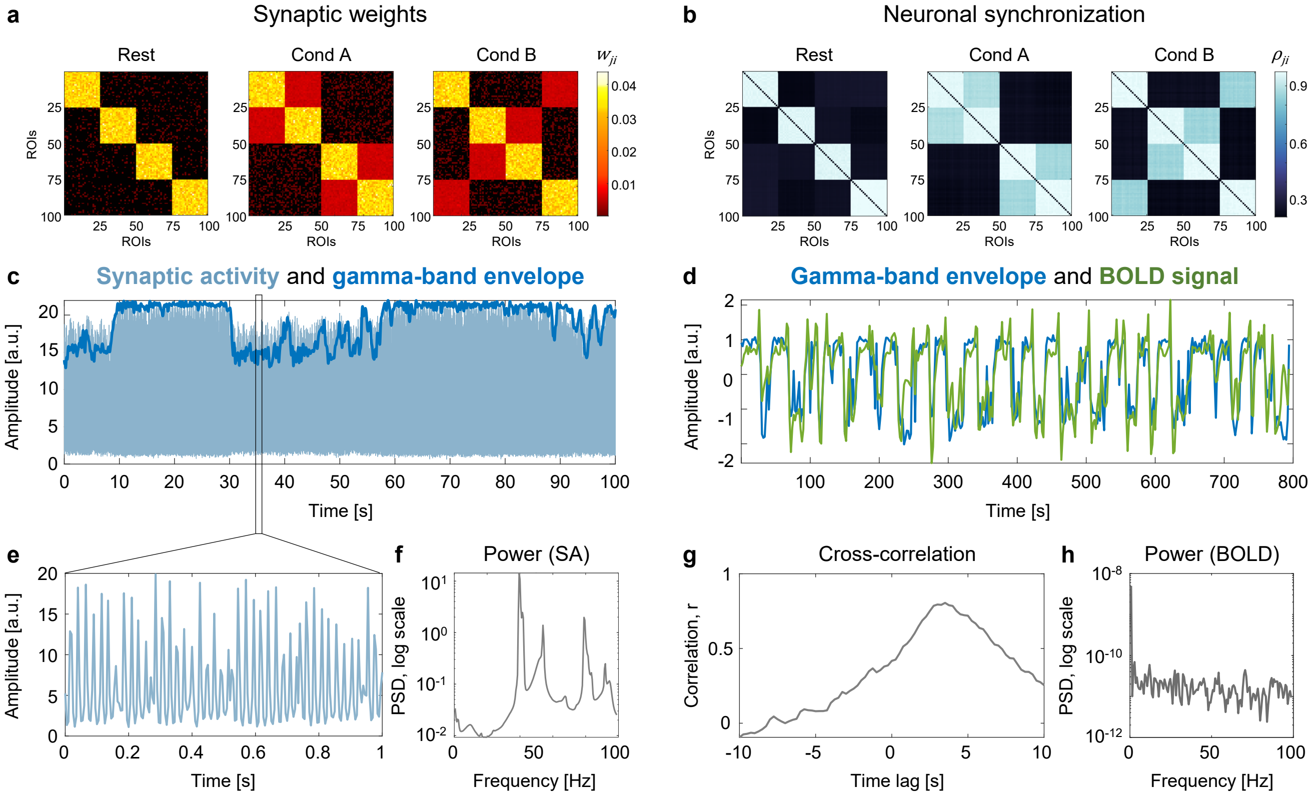


**Figure S5. The emergence of ultra-slow BOLD signal fluctuations from ultra-slow modulations of fast oscillatory activity (block design simulation). (a)** Ground-truth symmetric synaptic weight ($w_{ji}$) matrices for a single subject. For the long-range connections between Wilson-Cowan units ($w_{ji}$), we used three synaptic weight matrices corresponding to two task conditions ("Cond A" and "Cond B") and interim "Rest" periods. Each synaptic weight matrix consisted of 100 brain regions and 4 functional modules. **(b)** Gamma-band neuronal synchronisation estimated by the phase-locking value ($\rho_{ji}$) for a single subject. The simulation was performed for the block design with ten 20 s blocks per condition. **(c)** Example of simulated synaptic activity (SA) and gamma-band envelope for one of 100 connected brain regions (ROIs). **(d)** The time series of the gamma-band envelope and BOLD signal generated by the Balloon-Windkessel model based on simulated SA. We used the standard parameters of the haemodynamic model (Friston et al., 2003). **(e)** One second of simulated SA. **(f)** Power spectral density (PSD) of simulated SA. The main peak at 40 Hz (gamma-band oscillations). **(g)** Cross-correlation between the gamma-band envelope and BOLD signal. The maximum correlation $r = 0.81$ corresponds to a time lag of 3.5 seconds. **(h)** Power spectral density of the BOLD signal.

#### Supplementary Figure S6

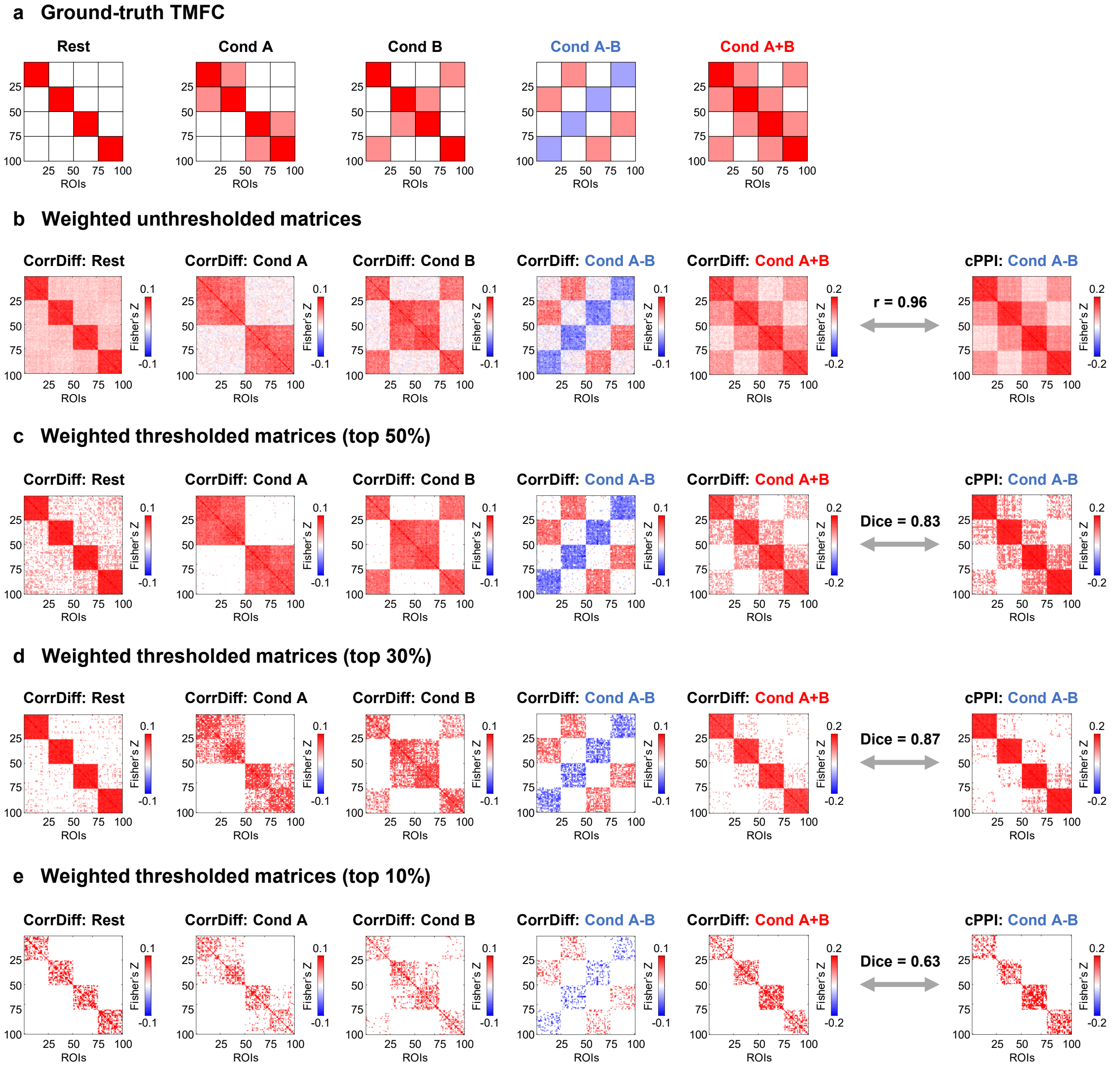


**Figure S6. The cPPI method fails to estimate TMFC.** Simulation results for the block design with ten 20 s blocks per condition, sample size N = 100, SNR = 0.4, and TR = 2 s. **(a)** Expected FC matrices based on ground-truth synaptic weight matrices. **(b)** Relationship between weighted unthresholded FC matrices obtained by the correlation difference (CorrDiff) and correlational PPI (cPPI) methods. To evaluate the similarity between unthresholded matrices, we calculated Pearson's r correlation. **(c-e)** Relationship between weighted thresholded FC matrices obtained by the CorrDiff and cPPI methods. To evaluate similarity between thresholded matrices, we calculated Dice coefficients. In this case, the original Dice coefficients were numerically identical to the signed Dice coefficients (SDC). Matrices were thresholded to the **(c)** top 50% connections, **(d)** top 30% connections, and **(e)** top 10% connections. In all cases, the cPPI matrices calculated for the "Cond A-B" contrast were similar to the CorrDiff matrices calculated for the "Cond A+B" contrast. **(c)** The top 50% and top 30% of connections of the cPPI matrix reflect summary FC in both task conditions and resting state. **(e)** The top 10% of connections of the cPPI matrix reflect resting-state (task-independent) connections. The PPI terms for the "Cond A-B" contrast were calculated using deconvolution. The color scales were adjusted for each matrix based on the maximum absolute value and were assured to be positive and negative symmetrical.

#### Supplementary Figure S7

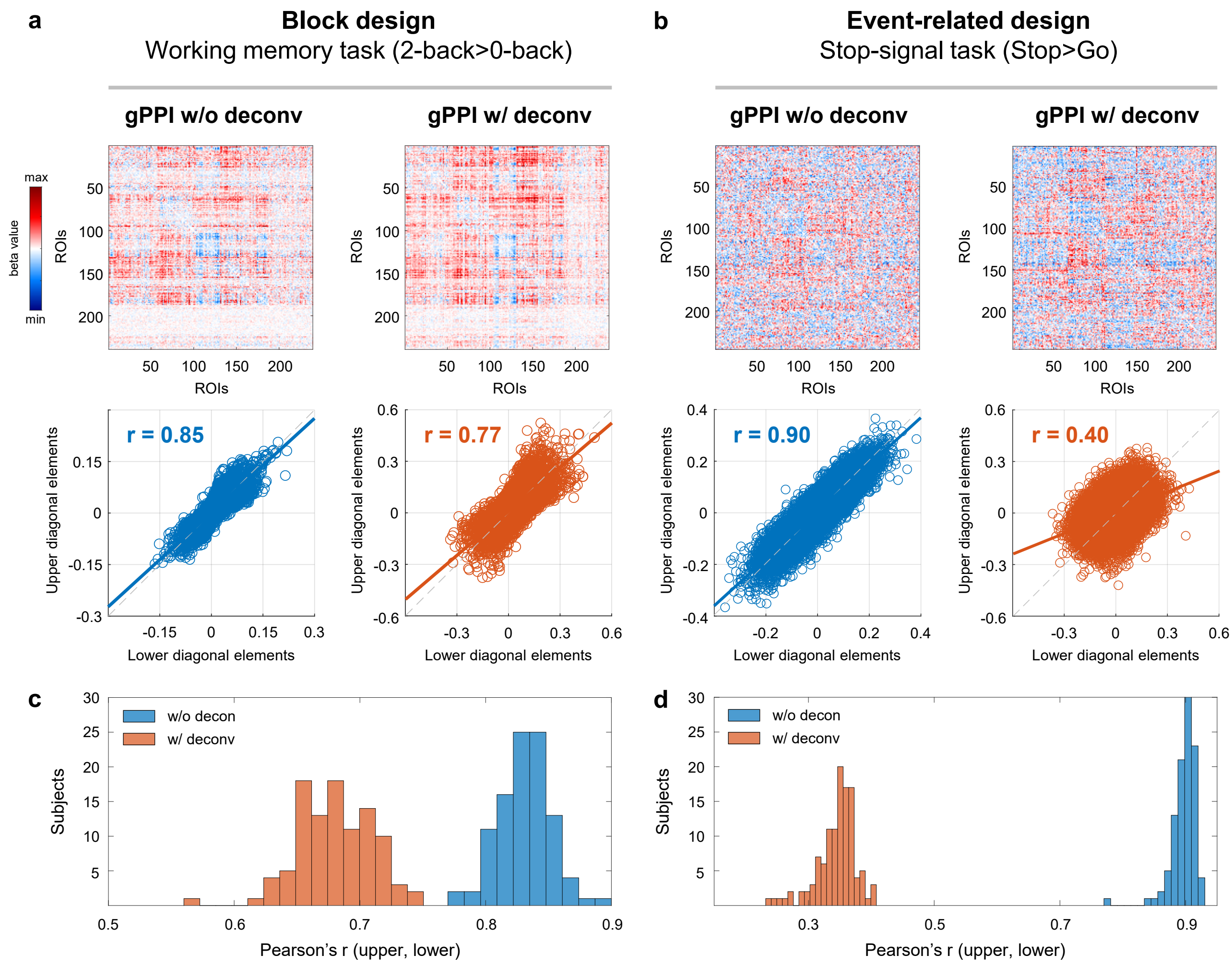


**Figure S7. Asymmetry of gPPI matrices in block and event-related designs.** Results for the working memory task and the stop-signal task. **(a, b)** Group-mean gPPI matrices without symmetrisation procedure for the block and event-related design tasks (upper panels). Scatterplots for the upper and lower diagonal elements of the corresponding gPPI matrices (lower panels). **(c, d)** Histograms of correlations between the upper and lower diagonal elements of individual subjects' matrices. Blue and red colours represent gPPI results without (w/o) and with (w/) the deconvolution step, respectively. The color scales were adjusted for each matrix based on the maximum absolute value and were assured to be positive and negative symmetrical.

#### Supplementary Figure S8

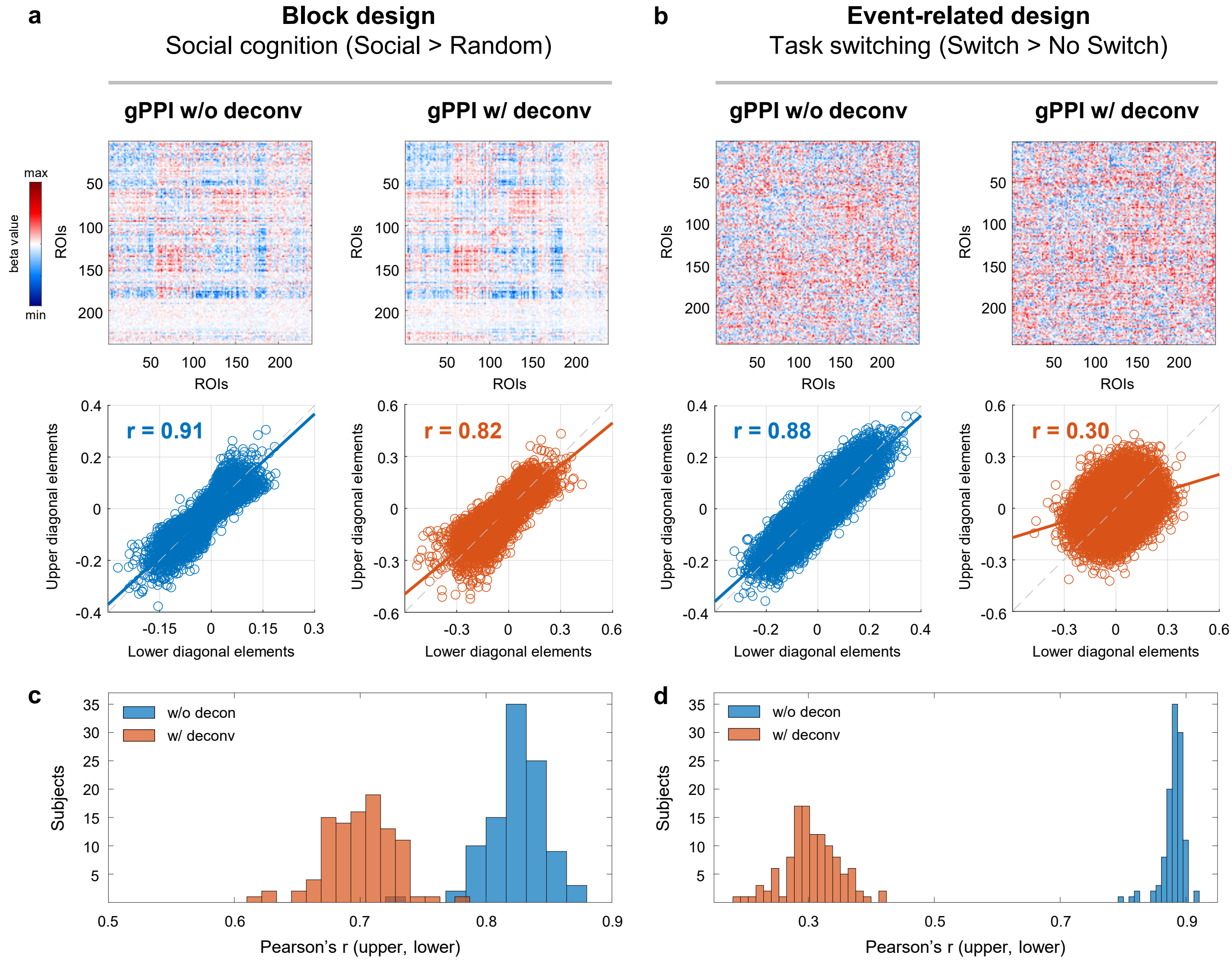


**Figure S8. Asymmetry of gPPI matrices in block and event-related designs.** Results for the social cognition task and the task switching task. **(a, b)** Group-mean gPPI matrices without symmetrisation procedure for the block and event-related design tasks (upper panels). Scatterplots for the upper and lower diagonal elements of the corresponding gPPI matrices (lower panels). **(c, d)** Histograms of correlations between the upper and lower diagonal elements of individual subjects' matrices. Blue and red colours represent gPPI results without (w/o) and with (w/) the deconvolution step, respectively. The color scales were adjusted for each matrix based on the maximum absolute value and were assured to be positive and negative symmetrical.

#### Supplementary Figure S9

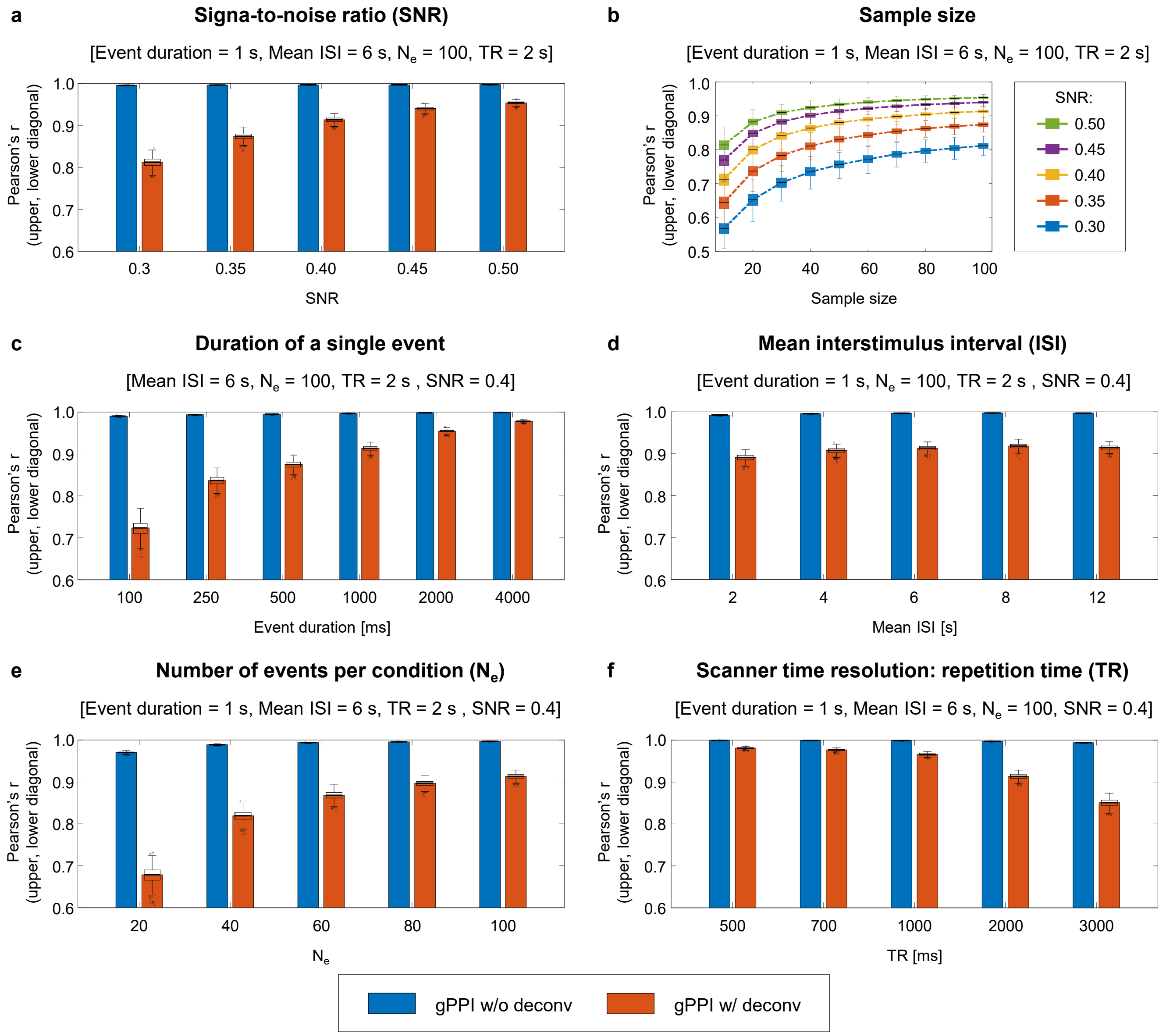


**Figure S9. Spurious asymmetry of gPPI matrices**. Here, we considered simulations with symmetric ground-truth matrices. We calculated correlations between the upper and lower diagonal elements of the group-mean gPPI matrices to assess matrix asymmetry. By default, we considered the event-related design to consist of one hundred 1 s events per condition, mean ISI = 6 s, TR = 2 s, and sample size N = 100. **(a)** Dependence of gPPI matrix asymmetry on SNR. **(b)** Asymmetry depending on sample size. Here, PPI terms were calculated using deconvolution. Without deconvolution, the asymmetry does not arise. **(c)** Asymmetry depending on the duration of a single event. **(d)** Asymmetry depending on the mean interstimulus (ISI) interval. **(e)** Asymmetry depending on the number of events per condition ($N_e$). **(f)** Asymmetry depending on the repetition time (TR) for a fixed scan time (23.6 minutes). Boxplots whiskers are drawn within the 1.5 interquartile range (IQR), computed from 1000 random resamplings with replacement. Blue and red bars represent gPPI matrix symmetry without (w/o) and with (w/) the deconvolution step, respectively.

#### Supplementary Figure S10

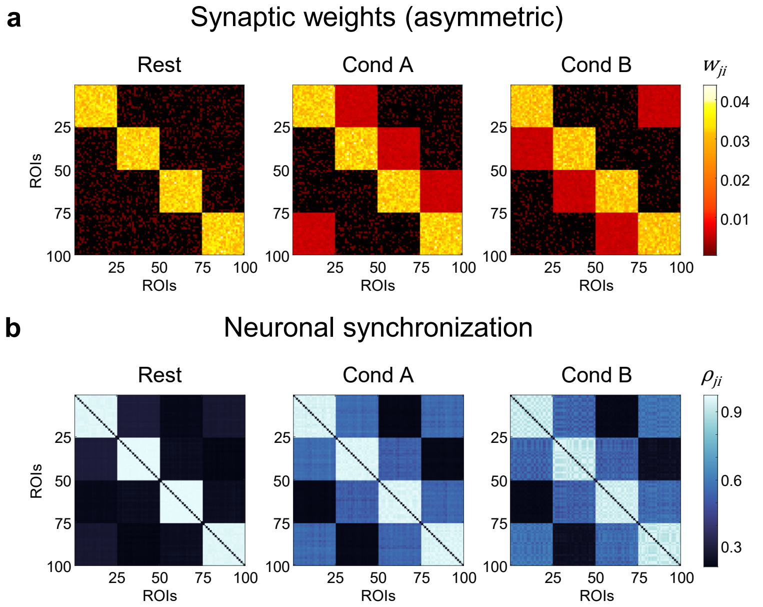


**Figure S10. Asymmetric ground-truth matrices. (a)** Ground-truth asymmetric synaptic weight ($w_{ji}$) matrices for a single subject. For the long-range connections between Wilson-Cowan units ($w_{ji}$), we used three synaptic weight matrices corresponding to two task conditions ("Cond A" and "Cond B") and interim "Rest" periods. Each synaptic weight matrix consisted of 100 brain regions and 4 functional modules. In "Cond A", synaptic weights were increased from module №1 to №4, from №4 to №3, from №3 to №2, and from №2 to №1. In "Cond B", synaptic weights were increased in the opposite direction. **(b)** Gamma-band neuronal synchronisation estimated by the phase-locking value ($\rho_{ji}$) for a single subject. The simulation was performed for the block design with ten 20 s blocks per condition. Note that phase-locking value (PLV) is a functional connectivity method that cannot reflect the direction of information flow and produces symmetrical matrices.

#### Supplementary Figure S11

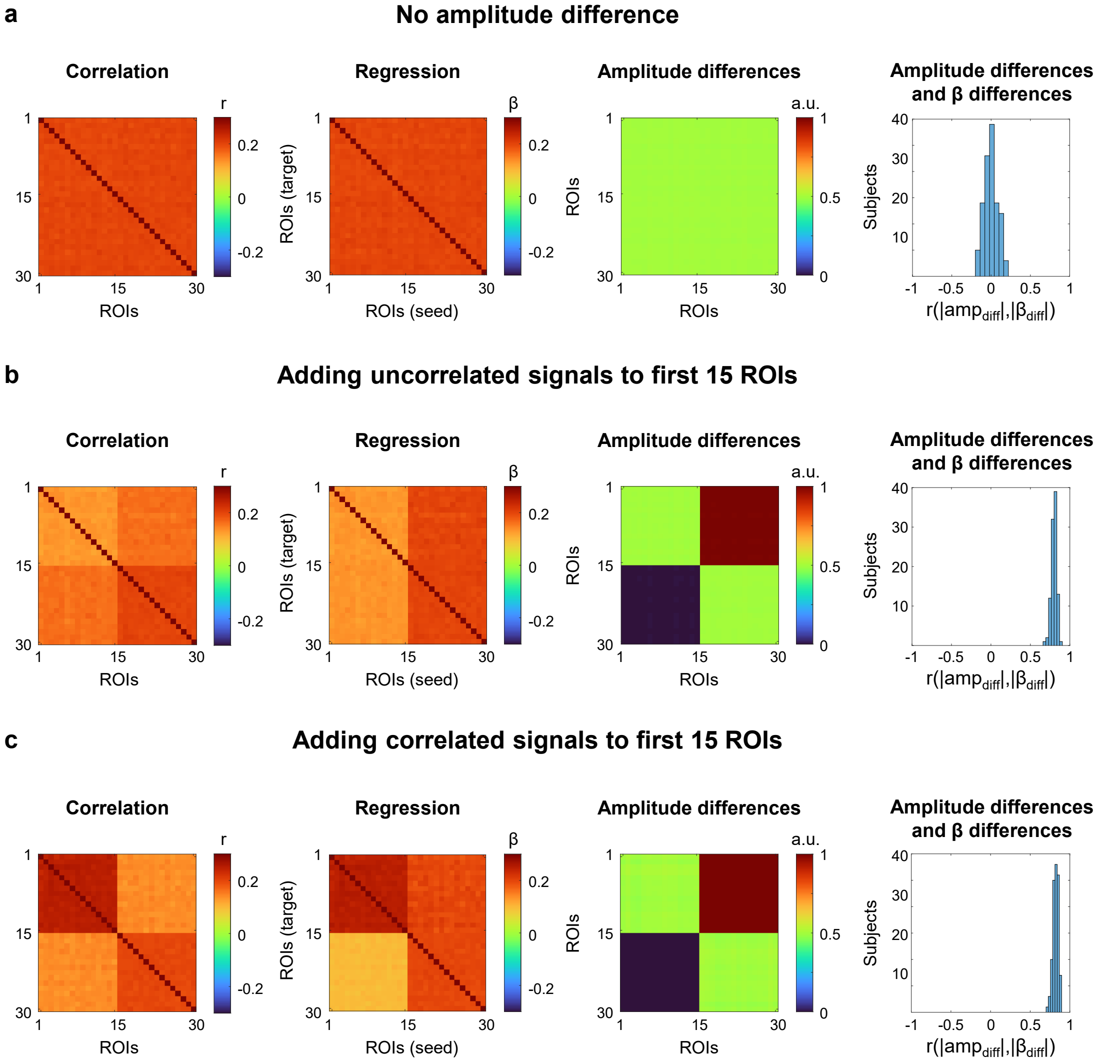


**Figure S11. Asymmetry of the regression coefficients due to amplitude differences.** To demonstrate the impact of amplitude differences on the regression coefficients asymmetry, we performed simple "toy" example simulations. **(a)** Correlated time series for 30 ROIs and 100 subjects were generated from a multivariate normal distribution with a mean of 100 and a covariance matrix with main diagonal elements equal 0.5 and off-diagonal elements equal 0.1. Each time series consisted of 400 time points. Correlation analysis performed using Pearson's r correlation. Regression analysis performed using simple linear regression. Histograms show the distribution of correlation between the absolute amplitude differences and beta coefficients differences (upper diagonal minus lower diagonal) for each subject. Mean correlation across subjects $\bar{r}_{(|amp_{diff}|,|\beta_{diff}|)}$ = -0.001. **(b)** Example #1. Adding uncorrelated signals to the first 15 ROIs (Module #1, M1), but not to the last 15 ROIs (Module #2, M2). The uncorrelated time series were generated from a normal distribution with a mean of one and a standard deviation of 0.5. Correlation and regression-based approaches demonstrate FC decrease within M1 and no decrease within M2. Regression-based approach show FC decrease from M1 to M2 (M1→M2, M1 – seed, M2 – target) and no decrease from M2 to M1 (M2→M1, M2 – seed, M1 – target). $\bar{r}_{(|amp_{diff}|,|\beta_{diff}|)}$ = 0.80. **(c)** Example #2. Adding correlated signals to the first 15 ROIs (Module #1, M1), but not to the last 15 ROIs (Module #2, M2). The correlated time series were generated from a multivariate normal distribution with a mean of one and a covariance matrix with main diagonal elements equal 0.5 and off-diagonal elements equal 0.15. Correlation and regression-based approaches demonstrate FC increase within M1 and no change within M2. As in previous example, regression-based approach show FC decrease from M1 to M2 and no decrease from M2 to M1. $\bar{r}_{(|amp_{diff}|,|\beta_{diff}|)}$ = 0.82.

#### Supplementary Figure S12

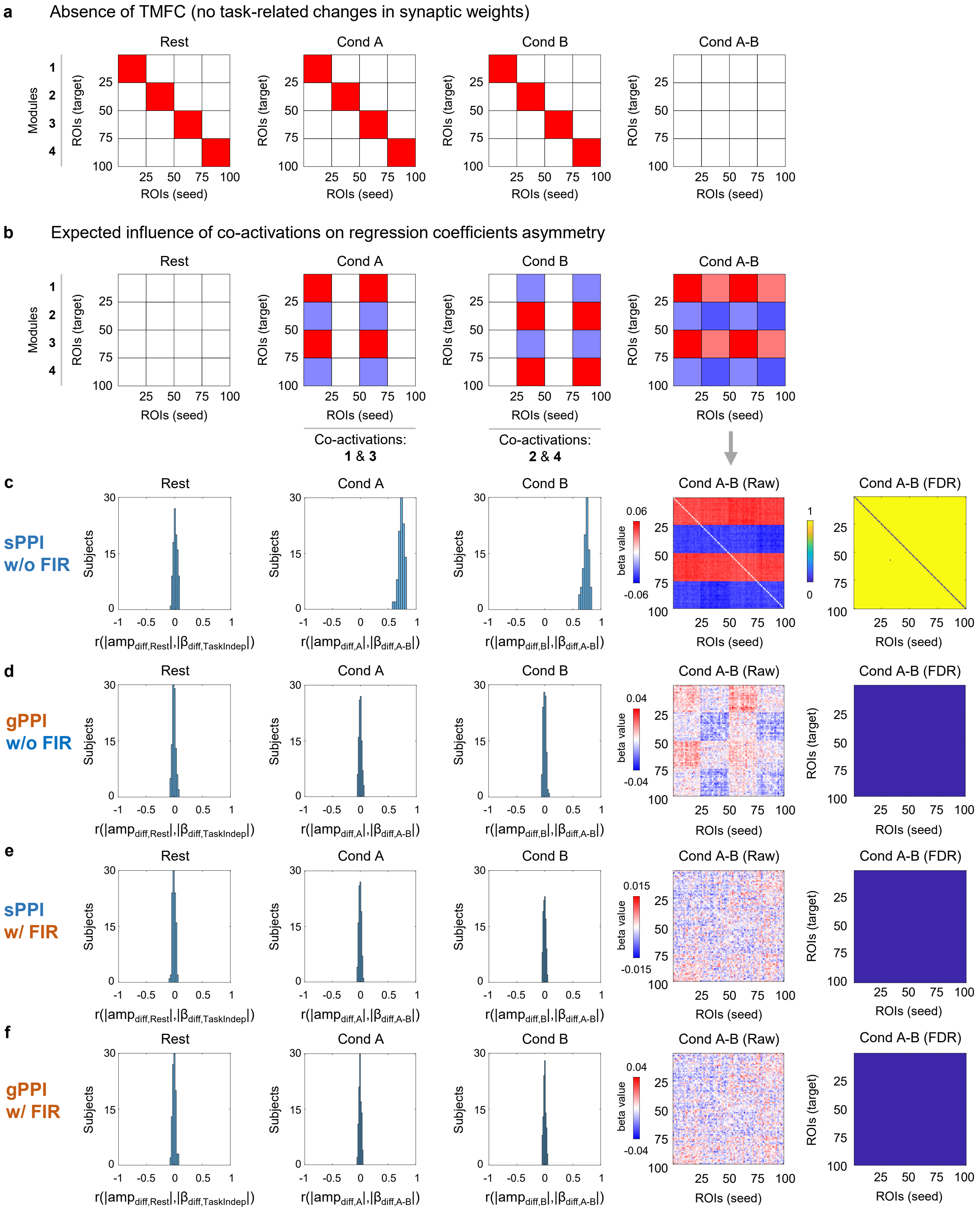


**Figure S12. Asymmetry of the regression coefficients due to co-activation.** In this example, we simulated BOLD signals without changing synaptic weights during task conditions, but adding co-activations. The simulation was performed for the block design with ten 20 s blocks per condition. SNR = 0.4, TR = 2 s. **(a)** Synaptic weights were the same for the resting condition, task condition A (Cond A) and condition B (Cond B) blocks. True TMFC is absent. Any significant difference in FC between the two task conditions (Cond A-B) will be a false positive. **(b)** Co-activations were added to modules 1 (M1) and 3 (M3) in condition A and to modules 2 (M2) and 4 (M4) in condition B. We can expect an increase in beta parameters within and between M1 and M3 for condition A, as well as M2 and M4 for condition B. We also can expect a decrease in M1→M2, M1→M4, M3→M2, M3→M4 for condition A and decrease in M2→M1, M2→M3, M4→M1, M4→M3 for condition B. **(c)** We obtained the expected asymmetric regression coefficients when using the sPPI method without FIR task regression. In this case, the sPPI method failed to eliminate the effects related to co-activations. We see significant FC differences between all regions after thresholding. All of them are false positives. The asymmetry of regression coefficients is due differences in the BOLD signal amplitudes, rather than causal directionality. Histograms show the distribution of correlations between the absolute amplitude differences during each task condition (Rest, Cond A, Cond B) and beta coefficients differences (difference between the upper and lower diagonal elements of the "Task Independent" matrix and "Cond A-B" matrix) for each subject. For the rest condition, mean correlation across subjects was close to zero, $\bar{r}_{(|amp_{diff,Rest}|,|\beta_{diff,TaskIndep}|)}$ = 0.013. However, correlations were high for task conditions. For condition A, $\bar{r}_{(|amp_{diff,A}|,|\beta_{diff,A-B}|)}$ = 0.736. For condition B, $\bar{r}_{(|amp_{diff,B}|,|\beta_{diff,A-B}|)}$ = 0.737. **(d)** When we used the gPPI method without FIR task regression, all false positives were eliminated. The asymmetry due to amplitude differences was also eliminated: $\bar{r}_{(|amp_{diff,Rest}|,|\beta_{diff,TaskIndep}|)}$ = -0.005, $\bar{r}_{(|amp_{diff,A}|,|\beta_{diff,A-B}|)}$ = -0.004, $\bar{r}_{(|amp_{diff,B}|,|\beta_{diff,A-B}|)}$ = -0.003. **(e)** When we performed FIR task regression before TMFC analysis, the sPPI method did not produce false positives or demonstrate asymmetry due to amplitude differences, $\bar{r}_{(|amp_{diff,Rest}|,|\beta_{diff,TaskIndep}|)}$ = -0.009, $\bar{r}_{(|amp_{diff,A}|,|\beta_{diff,A-B}|)}$ = -0.004, $\bar{r}_{(|amp_{diff,B}|,|\beta_{diff,A-B}|)}$ = -0.003. **(f)** The same was true for the gPPI method with FIR task regression: $\bar{r}_{(|amp_{diff,Rest}|,|\beta_{diff,TaskIndep}|)}$ = -0.006, $\bar{r}_{(|amp_{diff,A}|,|\beta_{diff,A-B}|)}$ = -0.003, $\bar{r}_{(|amp_{diff,B}|,|\beta_{diff,A-B}|)}$ = -0.002. All PPI terms were calculated with deconvolution and with mean centering. sPPI and gPPI matrices were thresholded at $\alpha = 0.001$ (two-sided one-sample t test, false discovery rate (FDR) correction). sPPI and gPPI matrices were not symmetrised. The color scales were adjusted for each matrix based on the maximum absolute value and were assured to be positive and negative symmetrical.

#### Supplementary Figure S13

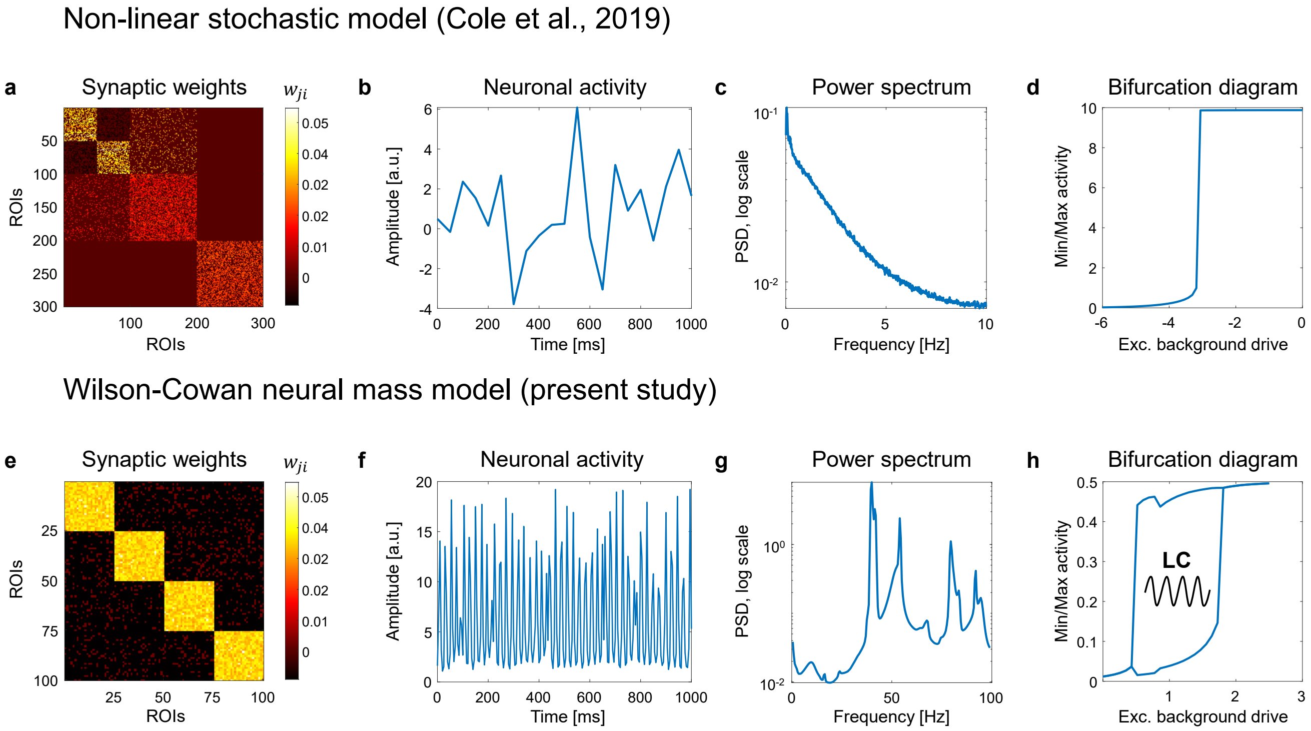


**Figure S13. Dynamics of neuronal activity produced by a non-linear stochastic model and the Wilson-Cowan neural mass model.** A simulation study by Cole et al. (2019) used a large-scale neural mass model with 300 regions obeying simplified Wilson-Cowan type equations. The authors sought to optimize the model simultaneously for both simplicity and biological plausibility. To simplify the model, they removed the inhibitory subpopulations and allowed the remaining populations to have both positive and negative synaptic weights. Similar neural mass reductions have also been used in several previous studies (Galan, 2008; Cabral et al., 2012; Stern et al., 2014; Messe et al., 2015). We reproduced the Cole et al. (2019) simulations, varying the model parameters, and did not observe limit-cycle oscillations. This may indicate that the dynamics of the Cole et al. (2019) model are driven only by the injected Gaussian noise. In the absence of background noise, the neuronal activity simulated by this non-linear stochastic model converge to a constant value, similar to the linear stochastic model (Galan, 2008; Cabral et al., 2012). Therefore, this non-linear stochastic model (NLSM) neglects the fast oscillatory nature of neuronal activity and does not produce the dynamics observed in electrophysiological data. In the current study, we used the Wilson-Cowan model (WCM) with separate excitatory and inhibitory subpopulations, which is able to produce self-sustained limit-cycle (LC) oscillations. **(a, e)** Synaptic weights between regions for NLSM and WCM, respectively. **(b, f)** One second of neuronal activity simulated by NLSM and WCM. **(c, g)** Power spectral density (PSD) of neuronal activity simulated by NLSM and WCM. NSLM gives rise to a 1/f power spectrum. At the same time, the power spectrum for the WCM has a main peak at 40 Hz (gamma-band oscillations). **(d, h)** Bifurcation diagrams for NLSM and WCM. They represent the relationship between the minimum/maximum neuronal activity in one of many coupled brain regions and the excitatory background drive in the absence of background noise. Excitatory background drive ($P_E$) corresponds to the bias parameter in Cole et al. (2019). NLSM has no bifurcation points and does not produce to LC oscillations. WCM has two bifurcation points and produces LC oscillations.

#### Supplementary Figure S14

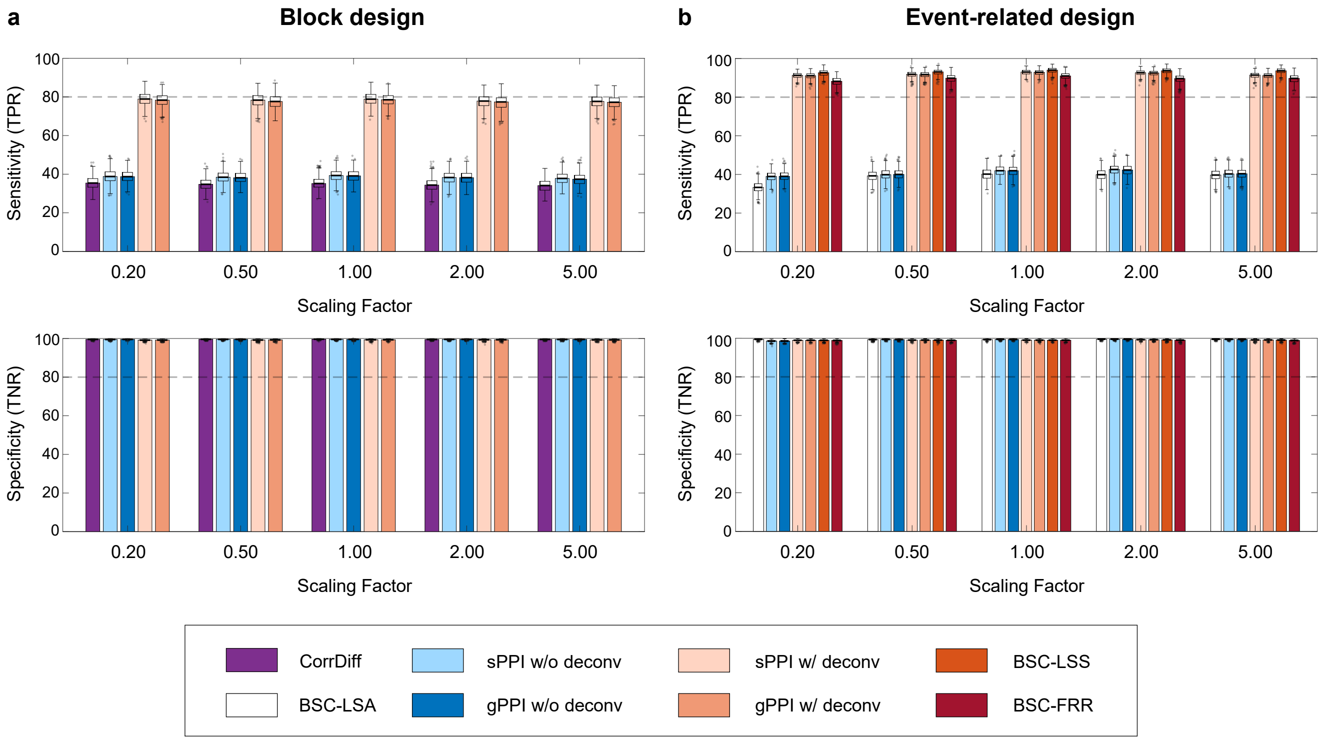


**Figure S14. Sensitivity and specificity of TMFC methods do not depend on the scaling factor after FIR task regression**. Simulation results for sample size N = 100, SNR = 0.4 and TR = 2 s. **(a)** Sensitivity (true positive rate, TPR) and specificity (true negative rate, TNR) of different TMFC methods for the block design with ten 20 s blocks per condition. **(b)** TPR and TNR for the event-related design with one hundred 1 s events per condition and mean ISI = 6 s. All TMFC matrices were calculated with FIR task regression. All PPI terms were calculated with and without the deconvolution step. sPPI and gPPI matrices were symmetrised. Boxplots whiskers are drawn within the 1.5 interquartile range (IQR), computed from 1000 random resamplings with replacement.

**Supplementary Figure S15**

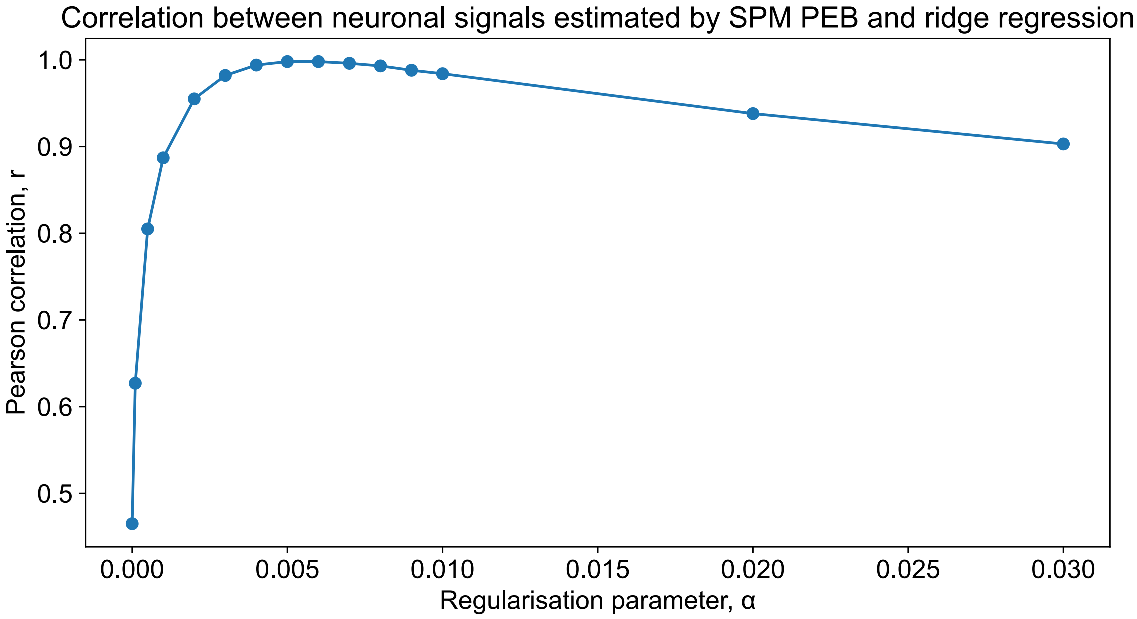


**Figure S15. Correlation between neuronal signals estimated by SPM12 PEB and ridge regression.** Results for the block design simulation. SNR = 0.4, TR = 2 s. The maximum correlation r = 0.998 is achieved with a regularisation parameter $\alpha$ = 0.005.

#### Supplementary Figure S16

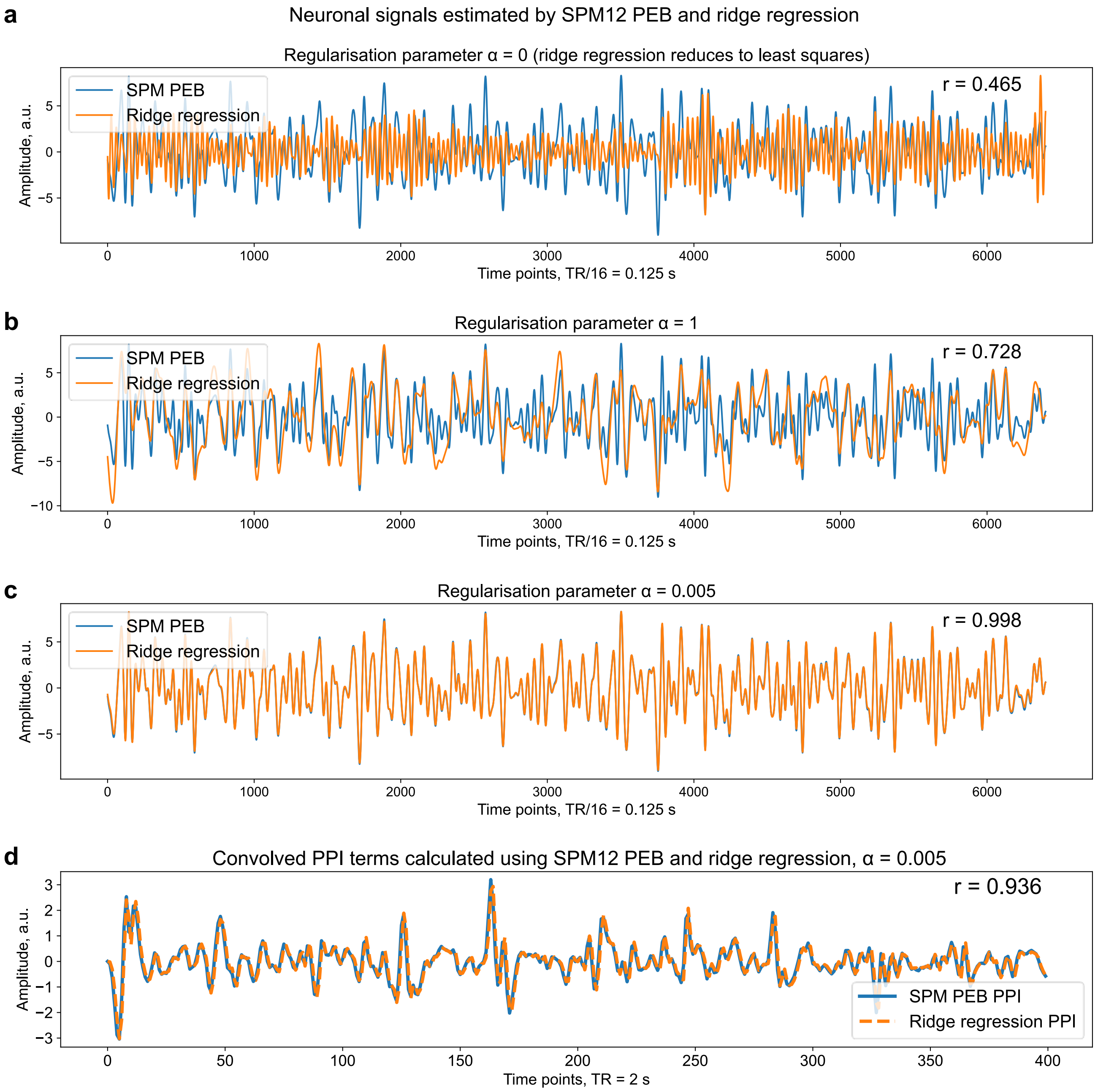


**Figure S16. Comparison of neuronal signals and PPIs estimated by SPM12 PEB and ridge regression.** Results for the block design simulation. SNR = 0.4, TR = 2 s. **(a-c)** Example of neuronal signals estimated using SPM12 PEB and ridge regression with different values of the regularisation parameter $\alpha$. **(a)** When $\alpha = 0$, ridge regression reduces to a least squares estimation. When the regularisation parameter is close to zero, there is a risk of overfitting. **(b)** Increasing regularisation parameter $\alpha$ results in a smoother estimated neuronal signal. **(c)** The maximum correlation between neuronal signals estimated by SPM12 PEB and ridge regression is achieved with a regularisation parameter $\alpha = 0.005$. **(d)** At $\alpha = 0.005$, the correlation between convolved PPI regressors calculated using SPM12 PEB and ridge regression is r = 0.936.

#### Supplementary Figure S17

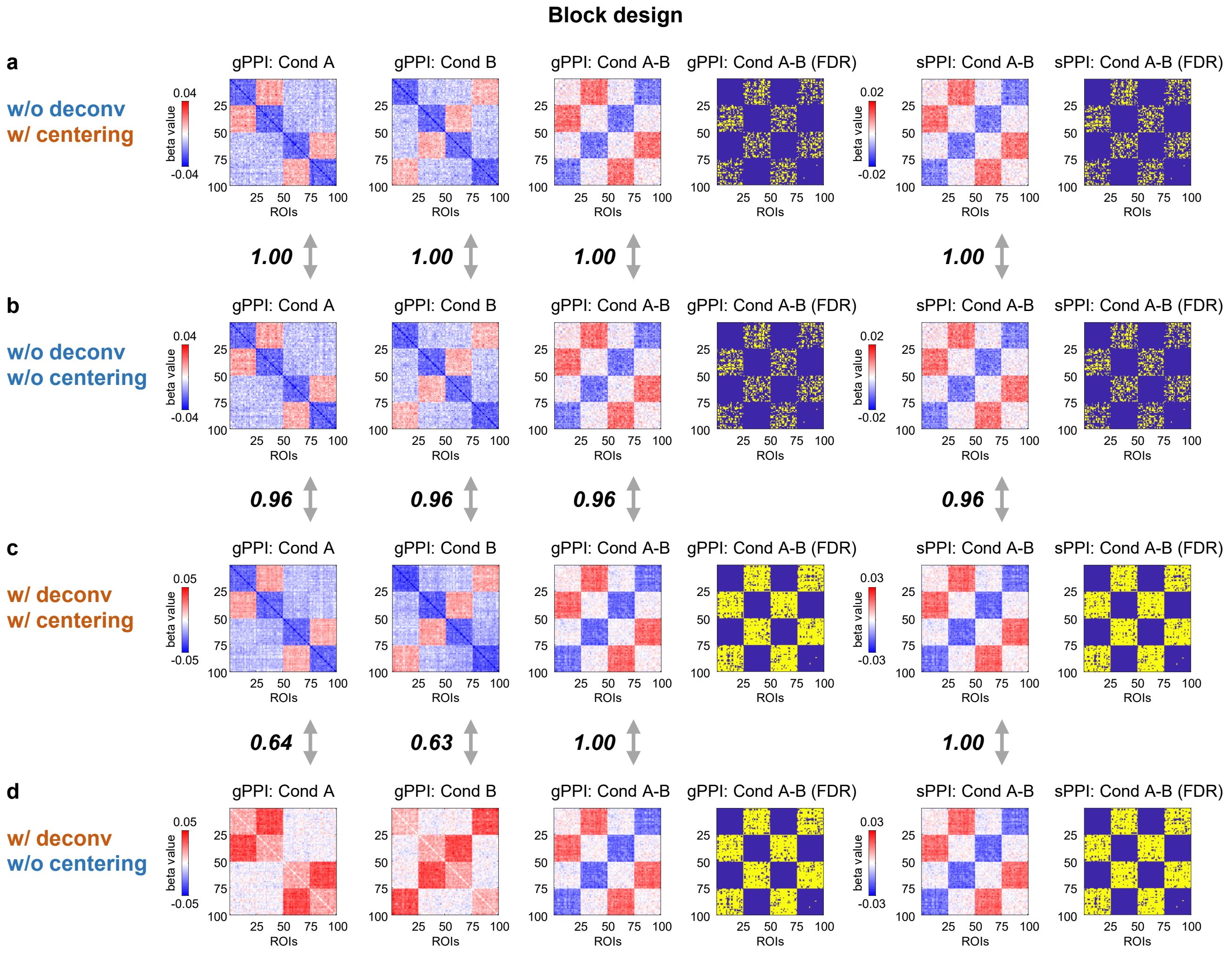


**Figure S17. Comparison PPI matrices estimated with and without mean centering of the task design regressor prior to PPI term calculation (block design simulation).** The simulation was performed for the block design with ten 20 s blocks per condition. SNR = 0.4, TR = 2 s. **(a)** PPI terms were calculated without (w/o) deconvolution and with (w/) mean centering. **(b)** PPI terms were calculated without (w/o) deconvolution and without (w/o) mean centering. **(c)** PPI terms were calculated with (w/) deconvolution and with (w/) mean centering. **(d)** PPI terms were calculated with (w/) deconvolution and without (w/o) mean centering. sPPI and gPPI matrices were thresholded at α = 0.001 (two-sided one-sample t test, false discovery rate (FDR) correction). To evaluate the similarity between matrices, we used Pearson's r correlation. sPPI and gPPI matrices were symmetrised. The color scales were adjusted for each matrix based on the maximum absolute value and were assured to be positive and negative symmetrical.

#### Supplementary Figure S18

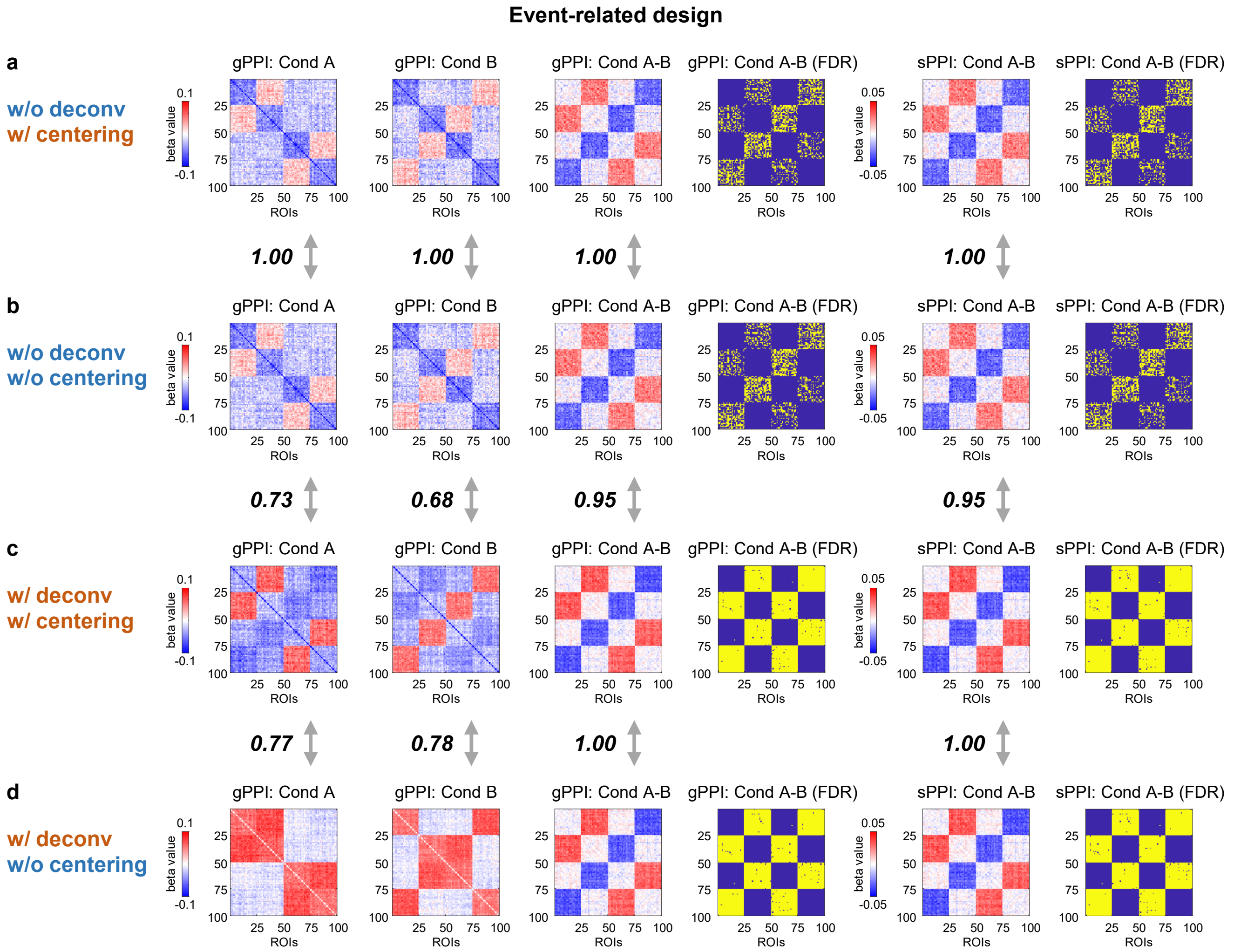


**Figure S18. Comparison PPI matrices estimated with and without mean centering of the task design regressor prior to PPI term calculation (event-related design simulation).** The simulation was performed for the event-related design with one hundred 1 s events per condition and mean interstimulus interval (ISI) = 6 s. SNR = 0.4, TR = 2 s. **(a)** PPI terms were calculated without (w/o) deconvolution and with (w/) mean centering. **(b)** PPI terms were calculated without (w/o) deconvolution and without (w/o) mean centering. **(c)** PPI terms were calculated with (w/) deconvolution and with (w/) mean centering. **(d)** PPI terms were calculated with (w/) deconvolution and without (w/o) mean centering. sPPI and gPPI matrices were thresholded at $\alpha = 0.001$ (two-sided one-sample t test, false discovery rate (FDR) correction). To evaluate the similarity between matrices, we used Pearson's r correlation. sPPI and gPPI matrices were symmetrised. The color scales were adjusted for each matrix based on the maximum absolute value and were assured to be positive and negative symmetrical.

#### Supplementary Figure S19

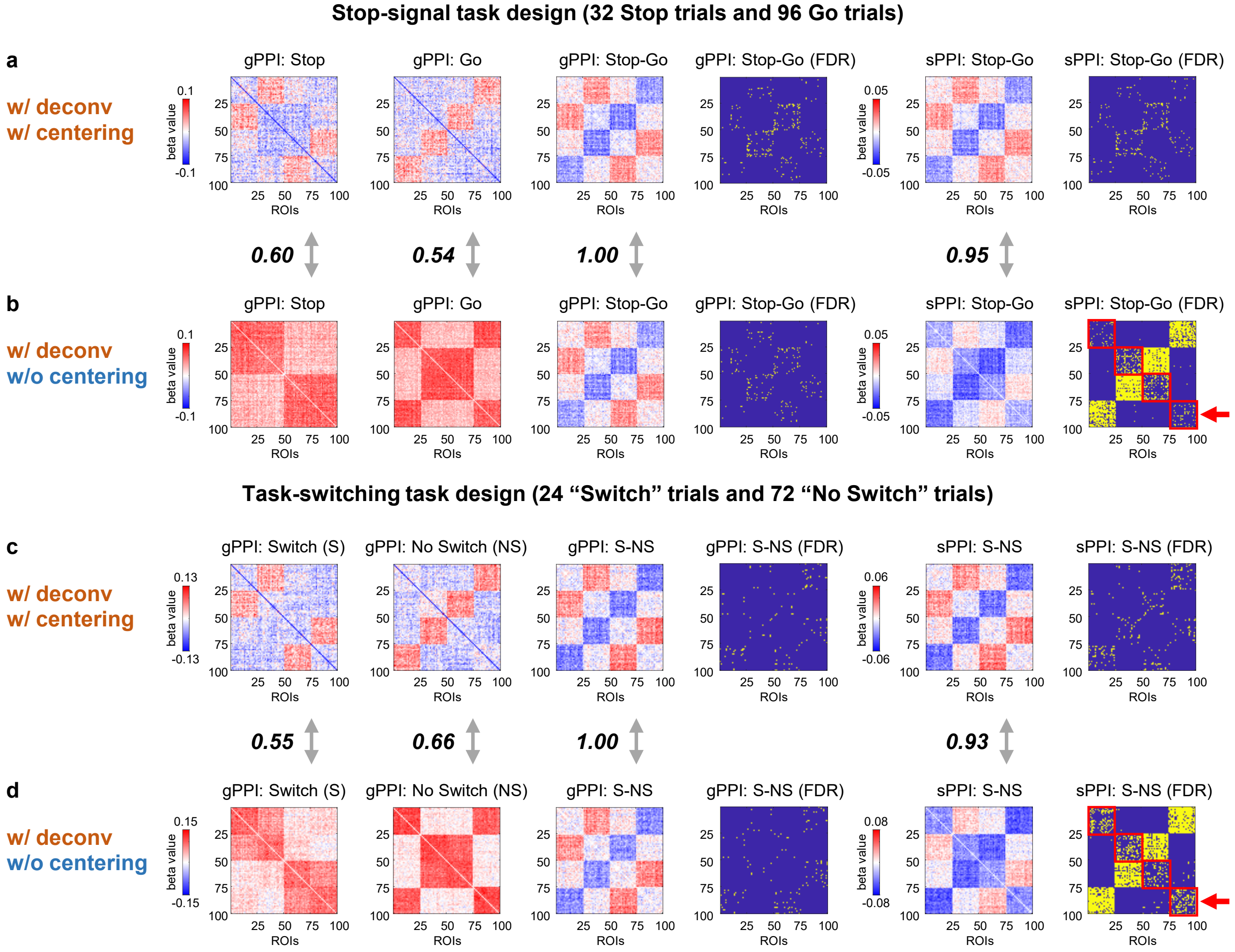


**Figure S19. Comparison PPI matrices estimated with and without mean centering of the task design regressor prior to PPI term calculation (unbalanced event-related designs from the CNP dataset).** The simulations were performed for the unbalanced event-related designs taken from the CNP dataset. The sample size was the same as for the CNP dataset, N = 115 subjects. SNR = 0.4, TR = 2 s. **(a-b)** The stop-signal task consisted of 32 Stop-signal trials and 96 Go trials lasted for 1.5 s. Mean ISI = 1 s. The contrast of interest is “Stop-Go”. **(c-d)** The task-switching task consisted of 24 “Switch” trials and 72 “No Switch” lasted for 1 s. Mean ISI = 3 s. **(a, c)** PPI terms were calculated (w/) deconvolution and with (w/) mean centering. **(b, d)** PPI terms were calculated with (w/) deconvolution and without (w/o) mean centering. **(a-d)** The gPPI method produces similar TMFC matrices (“Stop-Go” and “Switch-NoSwitch” or “S-NS”) w/ and w/o mean centering. **(a, c)** The sPPI method produces TMFC matrices similar to the gPPI method, only w/ mean centering. **(b, d)** W/o mean centering, the sPPI method shows greater sensitivity to FC modulations caused by task condition for which considerably more trials are available (Go condition for the stop-signal task and “No Switch” condition for the task-switching task, see connections between functional modules 1 and 4, 2 and 3). However, we also see significant FC differences within each functional module that are associated with task-unrelated spontaneous fluctuations when using sPPI w/o mean centering. False positive results are highlighted with red squares and red arrows. Task-unrelated spontaneous fluctuations are present both in the rest condition and task conditions (high connectivity within each functional module). Therefore, in each condition there are both task-related and task-unrelated effects. The difference between the two well-balanced conditions should subtract task-unrelated effects (here, effects within each functional module). However, in unbalanced designs this may not be the case. Using sPPI w/o mean centering, we see higher connectivity within each functional module in the condition with a large number of trials compared to the condition with a small number of trials. This false positive result can be avoided if we apply mean centering to the sPPI method or use the gPPI method w/ or w/o mean centering. sPPI and gPPI matrices were thresholded at α = 0.001 (two-sided one-sample t test, false discovery rate (FDR) correction). To evaluate the similarity between matrices, we used Pearson’s r correlation. sPPI and gPPI matrices were symmetrised. The color scales were adjusted for each matrix based on the maximum absolute value and were assured to be positive and negative symmetrical.

#### Supplementary Tables

##### Supplementary Table S1

Terms used to refer to different types of FC in this work and previous studies

| Definition of FC type | Terms used in previous studies | Terms used in this work |
|---|---|---|
| Correlation of the whole time series of the preprocessed resting-state BOLD signal | • **Rest FC** (Cole et al., 2014, 2019)<br>• **Rest-based FC** (Krienen et al., 2014; Greene et al., 2018)<br>• **Resting-state FC** (Rehme et al., 2013; Cole et al., 2014, 2019; Krienen et al., 2014; Watanabe et al., 2014; Greene et al., 2018; Lynch et al., 2018; Di & Biswal, 2019; Gao et al., 2019; Yang et al., 2020; Zhang et al., 2023; Zhao et al., 2023) | Resting-state FC (RSFC) |
| Correlation of the whole time series of the preprocessed task-state BOLD signal | • **Task FC** (Gao et al., 2019; Yang et al., 2020)<br>• **Task-based FC** (Rehme et al., 2013; Greene et al., 2018; Gao et al., 2019; Zhao et al., 2023)<br>• **Task-state FC** (Rehme et al., 2013; Lynch et al., 2018; Zhang et al., 2023) | Task-state FC (TSFC) |
| Correlation of the whole residual time series after task regression | • **Background FC** (Al-Aidroos et al., 2012; Norman-Haignere et al., 2012; Cordova et al., 2016)<br>• **Task-residualized FC** (Raud et al., 2023)<br>• **Task-model-residual FC** (Zhao et al., 2023) | Background FC (BGFC) |
| Correlation of the residual time series after task regression restricted to particular task condition | • **Task FC** (Cole et al., 2014, 2019; Krienen et al., 2014; Gratton et al., 2016)<br>• **Task-based FC** (Krienen et al., 2014)<br>• **Task-state FC** (Cole et al., 2019)<br>• **Background FC** (Griffis et al., 2015) | Correlation difference (CorrDiff) approach after FIR task regression (FC matrices calculated separately for “Condition A”, “Condition B” and “Rest” blocks) |
| Changes in FC measures (correlation or regression coefficients) during one condition compared to another, eliminating the influence of spontaneous task-independent fluctuations and co-activations | • **Task-based FC** (Poon et al., 2022; Doganci et al., 2023)<br>• **Task-state FC** (Cole et al., 2013, 2019)<br>• **Task-modulated FC** (Di & Biswal, 2019; Yang et al., 2020)<br>• **Task-related FC** (Fornito et al., 2012; Doganci et al., 2023)<br>• **Task-evoked FC** (Soreq et al., 2019)<br>• **Task-specific FC** (Watanabe et al., 2014)<br>• **Task-dependent FC** (Harrison et al., 2016; Xu et al., 2017; Poon et al., 2022)<br>• **Context-dependent FC** (Dodel et al., 2005; Cole et al., 2013; Harrison et al., 2016)<br>• **Context-modulated FC** (Cisler et al., 2014)<br>• **Context-specific FC** (Yin et al., 2021) | Task-modulated FC (TMFC) (“Condition A vs Condition B” contrast) |

#### Supplementary Table S2

Overview of previous TMFC simulation studies.

| Study | Neural activity simulation | Task design, time repetition (TR) | Objectives | Results | Limitations |
|---|---|---|---|---|---|
| **1) Gitelman et al. (2003)** | • Pair of regions<br>• Delta functions | • Event-related<br>• Single TR | • Introduce deconvolution step in sPPI | • Deconvolution step allow to model interactions at neuronal level | • Biophysically simplified model<br>• Single TR |
| **2) Kim & Horwitz (2008)** | • Large-scale (8 regions)<br>• Wilson-Cowan model | • Block<br>• Distinct TR | • Compare correlation difference and sPPI approaches<br>• Validate deconvolution step<br>• Investigate how TR and haemodynamic delay affect sPPI performance | • PPI better reflect TMFC than simple correlation difference<br>• sPPI with and without deconvolution produces similar results<br>• TR and haemodynamic delay do not have much of an impact on TMFC | • Block design only |
| **3) McLaren et al. (2012)** | • Pairs of regions<br>• Boxcar functions | • Event-related<br>• Single TR | • Introduce gPPI approach<br>• Compare sPPI and gPPI approaches | • gPPI is suitable for designs with more than two conditions<br>• gPPI estimate TMFC better than sPPI | • Biophysically simplified model<br>• Single TR |
| **4) Cisler et al. (2013)** | • Pair of regions<br>• Boxcar functions | • Block<br>• Event-related<br>• Single TR | • Compare sPPI, gPPI and BSC approaches for different task designs | • gPPI and BSC are more powerful than sPPI<br>• gPPI better for block designs<br>• BSC better for event-related designs with more trial repetitions | • Biophysically simplified model<br>• Single TR |
| **5) Abdulrahman & Henson (2016)** | • Pair of regions<br>• Delta functions | • Event-related<br>• Single TR | • Compare BSC based on Least squares all (LSA) and Least Squares Separate (LSA) approaches | • LSS better than LSA when scanner noise is higher than trial-by-trial variability (especially for rapid designs) | • Biophysically simplified model<br>• Single TR |
| **6) Cole et al. (2019)** | • Large-scale (300 regions)<br>• Non-linear stochastic model | • Block<br>• Single TR | • Evaluate FC inflation induced by task co-activations | • TMFC estimation by gPPI is inflated by task co-activations | • Block design only<br>• Limited biophysical interpretability<br>• Single TR |

#### Supplementary Table S3

Marginal mean comparison of the sensitivity of TMFC methods for the block design.

| Comparison | | Mean | SE | t | $p_{bonf}$ | Prior Odds | Posterior Odds | $Log(BF_{10,U})$ |
|---|---|---|---|---|---|---|---|---|
| CorrDiff | gPPI (w/ dec) | -29.757 | 0.053 | -559.255 | < .001 | 0.320 | ∞ | 1215.642 |
| | gPPI (w/o dec) | -2.659 | 0.053 | -49.975 | < .001 | 0.320 | 772.877 | 7.791 |
| | sPPI (w/ dec) | -30.126 | 0.053 | -566.197 | < .001 | 0.320 | ∞ | 1248.660 |
| | sPPI (w/o dec) | -2.901 | 0.053 | -54.513 | < .001 | 0.320 | 6437.832 | 9.911 |
| **gPPI (w/ dec)** | gPPI (w/o dec) | 27.098 | 0.053 | 509.281 | < .001 | 0.320 | ∞ | 994.462 |
| | **sPPI (w/ dec)** | **-0.369** | **0.053** | **-6.942** | **< .001** | **0.320** | **0.009** | **-3.595** |
| | sPPI (w/o dec) | 26.856 | 0.053 | 504.742 | < .001 | 0.320 | ∞ | 973.814 |
| **gPPI (w/o dec)** | sPPI (w/ dec) | -27.467 | 0.053 | -516.222 | < .001 | 0.320 | ∞ | 1024.142 |
| | **sPPI (w/o dec)** | **-0.241** | **0.053** | **-4.539** | **< .001** | **0.320** | **0.008** | **-3.700** |
| sPPI (w/ dec) | sPPI (w/o dec) | 27.226 | 0.053 | 511.684 | < .001 | 0.320 | ∞ | 1003.137 |

The marginal means are averaged over the levels of the signal-to-noise ratio. p values were adjusted for 10 t tests using Bonferroni correction. Bayesian comparisons are based on the default t test with a Cauchy (0, $r = 1/2^{0.5}$) prior. The posterior odds have been corrected for multiple testing by fixing the prior probability that the null hypothesis holds across all comparisons to 0.5. The "U" in the Bayes factor denotes that it is uncorrected. PPI terms were calculated with (w/) and without (w/o) the deconvolution step (dec). sPPI and gPPI matrices were symmetrised. Comparisons that show at least "substantial" evidence in favour of the null hypothesis ($LogBF_{10} < 1.1$) are highlighted in bold.

#### Supplementary Table S4

Marginal mean comparison of the sensitivity of TMFC methods for the event-related design.

| Comparison | | Mean | SE | t | $p_{bonf}$ | Prior Odds | Posterior Odds | $Log(BF_{10,U})$ |
|---|---|---|---|---|---|---|---|---|
| BSC-FRR | BSC-LSA | 37.510 | 0.042 | 889.936 | < .001 | 0.219 | ∞ | 2188.752 |
| | BSC-LSS | -3.090 | 0.042 | -73.316 | < .001 | 0.219 | $1.241\times10^{+8}$ | 20.155 |
| | gPPI (w/ dec) | -1.504 | 0.042 | -35.686 | < .001 | 0.219 | 1.061 | 1.578 |
| | gPPI (w/o dec) | 36.654 | 0.042 | 869.630 | < .001 | 0.219 | ∞ | 2102.823 |
| | sPPI (w/ dec) | -1.678 | 0.042 | -39.810 | < .001 | 0.219 | 4.088 | 2.927 |
| | sPPI (w/o dec) | 36.618 | 0.042 | 868.764 | < .001 | 0.219 | ∞ | 2102.619 |
| **BSC-LSA** | BSC-LSS | -40.601 | 0.042 | -963.775 | < .001 | 0.219 | ∞ | 2622.547 |
| | gPPI (w/ dec) | -39.015 | 0.042 | -926.125 | < .001 | 0.219 | ∞ | 2372.147 |
| | **gPPI (w/o dec)** | **-0.856** | **0.042** | **-20.317** | **< .001** | **0.219** | **0.017** | **-2.556** |
| | sPPI (w/ dec) | -39.188 | 0.042 | -930.251 | < .001 | 0.219 | ∞ | 2397.846 |
| | **sPPI (w/o dec)** | **-0.892** | **0.042** | **-21.183** | **< .001** | **0.219** | **0.019** | **-2.447** |
| BSC-LSS | gPPI (w/ dec) | 1.586 | 0.042 | 37.650 | < .001 | 0.219 | 3.273 | 2.704 |
| | gPPI (w/o dec) | 39.745 | 0.042 | 943.458 | < .001 | 0.219 | ∞ | 2530.145 |
| | sPPI (w/ dec) | 1.412 | 0.042 | 33.524 | < .001 | 0.219 | 0.879 | 1.390 |
| | sPPI (w/o dec) | 39.708 | 0.042 | 942.592 | < .001 | 0.219 | ∞ | 2530.481 |
| **gPPI (w/ dec)** | gPPI (w/o dec) | 38.159 | 0.042 | 905.808 | < .001 | 0.219 | ∞ | 2283.728 |
| | **sPPI (w/ dec)** | **-0.174** | **0.042** | **-4.126** | **< .001** | **0.219** | **0.005** | **-3.718** |
| | sPPI (w/o dec) | 38.122 | 0.042 | 904.942 | < .001 | 0.219 | ∞ | 2283.728 |
| **gPPI (w/o dec)** | sPPI (w/ dec) | -38.332 | 0.042 | -909.934 | < .001 | 0.219 | ∞ | 2309.017 |
| | **sPPI (w/o dec)** | **-0.037** | **0.042** | **-0.867** | **1.000** | **0.219** | **0.005** | **-3.790** |
| sPPI (w/ dec) | sPPI (w/o dec) | 38.296 | 0.042 | 909.068 | < .001 | 0.219 | ∞ | 2309.050 |

The marginal means are averaged over the levels of the signal-to-noise ratio. p values were adjusted for 21 t tests using Bonferroni correction. Bayesian comparisons are based on the default t test with a Cauchy (0, $r = 1/2^{0.5}$) prior. The posterior odds have been corrected for multiple testing by fixing the prior probability that the null hypothesis holds across all comparisons to 0.5. The "U" in the Bayes factor denotes that it is uncorrected. PPI terms were calculated with (w/) and without (w/o) the deconvolution step (dec). sPPI and gPPI matrices were symmetrised. Comparisons that show at least "substantial" evidence in favour of the null hypothesis ($LogBF_{10} < 1.1$) are highlighted in bold.

**Supplementary Table S5**

Pairwise comparison of the sensitivity of TMFC methods for event-related designs with different event durations.

| Comparison | | Mean | SE | t | $p_{bonf}$ | Prior Odds | Posterior Odds | Log($BF_{10,\,U}$) |
|---|---|---|---|---|---|---|---|---|
| | | | | **Event duration = 100 ms** | | | | |
| BSC-FRR | BSC-LSA | 12.238 | 0.100 | 122.217 | < .001 | 0.047 | ∞ | 2118.210 |
| | BSC-LSS | -1.757 | 0.100 | -17.551 | < .001 | 0.047 | $9.592\times10^{+29}$ | 72.087 |
| | gPPI (w/ dec) | 0.270 | 0.100 | 2.700 | 1.000 | 0.047 | 0.023 | -0.721 |
| | sPPI (w/ dec) | 0.294 | 0.100 | 2.937 | 1.000 | 0.047 | 0.034 | -0.323 |
| BSC-LSA | BSC-LSS | -13.995 | 0.100 | -139.768 | < .001 | 0.047 | ∞ | 2358.555 |
| | gPPI (w/ dec) | -11.968 | 0.100 | -119.517 | < .001 | 0.047 | ∞ | 2489.800 |
| | sPPI (w/ dec) | -11.944 | 0.100 | -119.280 | < .001 | 0.047 | ∞ | 2471.492 |
| BSC-LSS | gPPI (w/ dec) | 2.028 | 0.100 | 20.251 | < .001 | 0.047 | $5.426\times10^{+49}$ | 117.569 |
| | sPPI (w/ dec) | 2.052 | 0.100 | 20.488 | < .001 | 0.047 | $3.707\times10^{+50}$ | 119.491 |
| gPPI (w/ dec) | sPPI (w/ dec) | 0.024 | 0.100 | 0.237 | 1.000 | 0.047 | 0.002 | -2.968 |
| | | | | **Event duration = 250 ms** | | | | |
| BSC-FRR | BSC-LSA | 48.364 | 0.100 | 482.999 | < .001 | 0.047 | ∞ | 4134.168 |
| | BSC-LSS | -6.152 | 0.100 | -61.441 | < .001 | 0.047 | $6.244\times10^{+211}$ | 490.728 |
| | gPPI (w/ dec) | 1.111 | 0.100 | 11.099 | < .001 | 0.047 | $2.477\times10^{+7}$ | 20.077 |
| | sPPI (w/ dec) | 1.010 | 0.100 | 10.089 | < .001 | 0.047 | 347039.912 | 15.809 |
| BSC-LSA | BSC-LSS | -54.516 | 0.100 | -544.440 | < .001 | 0.047 | ∞ | 4464.545 |
| | gPPI (w/ dec) | -47.253 | 0.100 | -471.900 | < .001 | 0.047 | ∞ | 4389.013 |
| | sPPI (w/ dec) | -47.354 | 0.100 | -472.911 | < .001 | 0.047 | ∞ | 4364.078 |
| BSC-LSS | gPPI (w/ dec) | 7.264 | 0.100 | 72.540 | < .001 | 0.047 | ∞ | 724.413 |
| | sPPI (w/ dec) | 7.162 | 0.100 | 71.529 | < .001 | 0.047 | $2.074\times10^{+303}$ | 701.464 |
| gPPI (w/ dec) | sPPI (w/ dec) | -0.101 | 0.100 | -1.011 | 1.000 | 0.047 | 0.003 | -2.757 |
| | | | | **Event duration = 500 ms** | | | | |
| BSC-FRR | BSC-LSA | 58.066 | 0.100 | 579.889 | < .001 | 0.047 | ∞ | 4492.852 |
| | BSC-LSS | -4.311 | 0.100 | -43.048 | < .001 | 0.047 | $6.935\times10^{+169}$ | 394.125 |
| | gPPI (w/ dec) | 0.290 | 0.100 | 2.898 | 1.000 | 0.047 | 0.02 | -0.85 |
| | sPPI (w/ dec) | -0.013 | 0.100 | -0.133 | 1.000 | 0.047 | 0.002 | -2.986 |
| BSC-LSA | BSC-LSS | -62.376 | 0.100 | -622.937 | < .001 | 0.047 | ∞ | 4834.299 |
| | gPPI (w/ dec) | -57.776 | 0.100 | -576.991 | < .001 | 0.047 | ∞ | 4648.812 |
| | sPPI (w/ dec) | -58.079 | 0.100 | -580.022 | < .001 | 0.047 | ∞ | 4700.323 |
| BSC-LSS | gPPI (w/ dec) | 4.601 | 0.100 | 45.946 | < .001 | 0.047 | $3.355\times10^{+216}$ | 501.620 |
| | sPPI (w/ dec) | 4.297 | 0.100 | 42.915 | < .001 | 0.047 | $3.986\times10^{+200}$ | 464.951 |
| gPPI (w/ dec) | sPPI (w/ dec) | -0.304 | 0.100 | -3.032 | 1.000 | 0.047 | 0.042 | -0.126 |
| | | | | **Event duration = 1000 ms** | | | | |
| BSC-FRR | BSC-LSA | 50.693 | 0.100 | 506.258 | < .001 | 0.047 | ∞ | 4675.406 |
| | BSC-LSS | -3.012 | 0.100 | -30.080 | < .001 | 0.047 | $7.787\times10^{+262}$ | 608.381 |
| | gPPI (w/ dec) | -1.846 | 0.100 | -18.439 | < .001 | 0.047 | $4.955\times10^{+118}$ | 276.357 |
| | sPPI (w/ dec) | -2.028 | 0.100 | -20.249 | < .001 | 0.047 | $1.535\times10^{+144}$ | 335.052 |
| BSC-LSA | BSC-LSS | -53.705 | 0.100 | -536.338 | < .001 | 0.047 | ∞ | 4917.987 |
| | gPPI (w/ dec) | -52.539 | 0.100 | -524.697 | < .001 | 0.047 | ∞ | 4883.486 |
| | sPPI (w/ dec) | -52.721 | 0.100 | -526.507 | < .001 | 0.047 | ∞ | 4907.266 |
| BSC-LSS | gPPI (w/ dec) | 1.166 | 0.100 | 11.641 | < .001 | 0.047 | $1.783\times10^{+68}$ | 160.206 |
| | sPPI (w/ dec) | 0.984 | 0.100 | 9.831 | < .001 | 0.047 | $2.661\times10^{+51}$ | 121.462 |
| gPPI (w/ dec) | sPPI (w/ dec) | -0.181 | 0.100 | -1.810 | 1.000 | 0.047 | 0.296 | 1.604 |
| | | | | **Event duration = 2000 ms** | | | | |
| BSC-FRR | BSC-LSA | 14.547 | 0.100 | 145.280 | < .001 | 0.047 | ∞ | 3438.020 |
| | BSC-LSS | -0.207 | 0.100 | -2.068 | 1.000 | 0.047 | $7.783\times10^{+192}$ | 447.200 |
| | gPPI (w/ dec) | -0.169 | 0.100 | -1.690 | 1.000 | 0.047 | $5.743\times10^{+122}$ | 258.715 |
| | sPPI (w/ dec) | -0.173 | 0.100 | -1.731 | 1.000 | 0.047 | $1.001\times10^{+133}$ | 309.296 |
| BSC-LSA | BSC-LSS | -14.754 | 0.100 | -147.348 | < .001 | 0.047 | ∞ | 3472.149 |
| | gPPI (w/ dec) | -14.717 | 0.100 | -146.970 | < .001 | 0.047 | ∞ | 3465.769 |
| | sPPI (w/ dec) | -14.721 | 0.100 | -147.011 | < .001 | 0.047 | ∞ | 3466.808 |
| BSC-LSS | gPPI (w/ dec) | 0.038 | 0.100 | 0.378 | 1.000 | 0.047 | $1.107\times10^{+14}$ | 35.389 |
| | sPPI (w/ dec) | 0.034 | 0.100 | 0.337 | 1.000 | 0.047 | $1.072\times10^{+12}$ | 30.752 |
| gPPI (w/ dec) | sPPI (w/ dec) | -0.004 | 0.100 | -0.042 | 1.000 | 0.047 | 0.004 | -2.591 |
| | | | | **Event duration = 4000 ms** | | | | |
| BSC-FRR | BSC-LSA | 0.508 | 0.100 | 5.075 | < .001 | 0.047 | ∞ | 1088.693 |
| | BSC-LSS | $-7.600\times10^{-4}$ | 0.100 | -0.008 | 1.000 | 0.047 | 6.450 | 4.915 |
| | gPPI (w/ dec) | $-8.400\times10^{-4}$ | 0.100 | -0.008 | 1.000 | 0.047 | 93.701 | 7.591 |
| | sPPI (w/ dec) | $-8.400\times10^{-4}$ | 0.100 | -0.008 | 1.000 | 0.047 | 93.701 | 7.591 |
| BSC-LSA | BSC-LSS | -0.509 | 0.100 | -5.083 | < .001 | 0.047 | ∞ | 1090.983 |
| | gPPI (w/ dec) | -0.509 | 0.100 | -5.084 | < .001 | 0.047 | ∞ | 1091.225 |
| | sPPI (w/ dec) | -0.509 | 0.100 | -5.084 | < .001 | 0.047 | ∞ | 1091.225 |
| BSC-LSS | gPPI (w/ dec) | $-8.000\times10^{-5}$ | 0.100 | $-7.989\times10^{-4}$ | 1.000 | 0.047 | 0.006 | -1.997 |
| | sPPI (w/ dec) | $-8.000\times10^{-5}$ | 0.100 | $-7.989\times10^{-4}$ | 1.000 | 0.047 | 0.006 | -1.997 |
| gPPI (w/ dec) | sPPI (w/ dec) | $8.289\times10^{-12}$ | 0.100 | $8.278\times10^{-11}$ | 1.000 | NaN | NaN | NaN |

p values were adjusted for 60 t tests using Bonferroni correction. Bayesian comparisons are based on the default t test with a Cauchy (0, $r = 1/2^{0.5}$) prior. The posterior odds have been corrected for multiple testing by fixing the prior probability that the null hypothesis holds across all comparisons to 0.5. The "U" in the Bayes factor denotes that it is uncorrected. PPI terms were calculated with the deconvolution step (w/dec). sPPI and gPPI matrices were symmetrised. NaN – data are essentially constant (sPPI and gPPI methods reached 100% sensitivity).

#### Supplementary Table S6

Pairwise comparison of the sensitivity of TMFC methods for event-related designs with different interstimulus intervals.

| Comparison | | Mean | SE | t | $p_{bonf}$ | Prior Odds | Posterior Odds | $Log(BF_{10,U})$ |
|---|---|---|---|---|---|---|---|---|
| | | | | **Mean ISI = 2 s** | | | | |
| BSC-FRR | BSC-LSA | 68.382 | 0.087 | 782.778 | < .001 | 0.057 | ∞ | 5321.434 |
| | BSC-LSS | -5.260 | 0.087 | -60.211 | < .001 | 0.057 | $1.009\times10^{+219}$ | 507.139 |
| | gPPI (w/ dec) | -10.759 | 0.087 | -123.165 | < .001 | 0.057 | ∞ | 1414.228 |
| | sPPI (w/ dec) | -11.257 | 0.087 | -128.862 | < .001 | 0.057 | ∞ | 1507.441 |
| BSC-LSA | BSC-LSS | -73.642 | 0.087 | -842.989 | < .001 | 0.057 | ∞ | 5659.180 |
| | gPPI (w/ dec) | -79.141 | 0.087 | -905.943 | < .001 | 0.057 | ∞ | 6077.271 |
| | sPPI (w/ dec) | -79.639 | 0.087 | -911.640 | < .001 | 0.057 | ∞ | 6171.758 |
| BSC-LSS | gPPI (w/ dec) | -5.500 | 0.087 | -62.954 | < .001 | 0.057 | $3.634\times10^{+279}$ | 646.576 |
| | sPPI (w/ dec) | -5.997 | 0.087 | -68.651 | < .001 | 0.057 | ∞ | 751.354 |
| gPPI (w/ dec) | sPPI (w/ dec) | -0.498 | 0.087 | -5.697 | < .001 | 0.057 | 22.545 | 5.980 |
| | | | | **Mean ISI = 4 s** | | | | |
| BSC-FRR | BSC-LSA | 74.820 | 0.087 | 856.473 | < .001 | 0.057 | ∞ | 5645.222 |
| | BSC-LSS | -4.831 | 0.087 | -55.306 | < .001 | 0.057 | ∞ | 743.103 |
| | gPPI (w/ dec) | -6.227 | 0.087 | -71.286 | < .001 | 0.057 | ∞ | 1139.939 |
| | sPPI (w/ dec) | -6.428 | 0.087 | -73.582 | < .001 | 0.057 | ∞ | 1199.142 |
| BSC-LSA | BSC-LSS | -79.651 | 0.087 | -911.779 | < .001 | 0.057 | ∞ | 5991.980 |
| | gPPI (w/ dec) | -81.047 | 0.087 | -927.758 | < .001 | 0.057 | ∞ | 6229.187 |
| | sPPI (w/ dec) | -81.248 | 0.087 | -930.054 | < .001 | 0.057 | ∞ | 6265.782 |
| BSC-LSS | gPPI (w/ dec) | -1.396 | 0.087 | -15.980 | < .001 | 0.057 | $1.443\times10^{+53}$ | 125.261 |
| | sPPI (w/ dec) | -1.596 | 0.087 | -18.275 | < .001 | 0.057 | $7.937\times10^{+70}$ | 166.117 |
| gPPI (w/ dec) | sPPI (w/ dec) | -0.201 | 0.087 | -2.296 | 1.000 | 0.057 | 0.112 | 0.675 |
| | | | | **Mean ISI = 6 s** | | | | |
| BSC-FRR | BSC-LSA | 50.692 | 0.087 | 580.283 | < .001 | 0.057 | ∞ | 4675.544 |
| | BSC-LSS | -3.013 | 0.087 | -34.487 | < .001 | 0.057 | $1.349\times10^{+263}$ | 608.744 |
| | gPPI (w/ dec) | -1.847 | 0.087 | -21.143 | < .001 | 0.057 | $7.750\times10^{+118}$ | 276.617 |
| | sPPI (w/ dec) | -2.028 | 0.087 | -23.218 | < .001 | 0.057 | $2.479\times10^{+144}$ | 335.344 |
| BSC-LSA | BSC-LSS | -53.705 | 0.087 | -614.769 | < .001 | 0.057 | ∞ | 4917.987 |
| | gPPI (w/ dec) | -52.539 | 0.087 | -601.426 | < .001 | 0.057 | ∞ | 4883.486 |
| | sPPI (w/ dec) | -52.721 | 0.087 | -603.501 | < .001 | 0.057 | ∞ | 4907.266 |
| BSC-LSS | gPPI (w/ dec) | 1.166 | 0.087 | 13.343 | < .001 | 0.057 | $2.150\times10^{+68}$ | 160.206 |
| | sPPI (w/ dec) | 0.984 | 0.087 | 11.269 | < .001 | 0.057 | $3.209\times10^{+51}$ | 121.462 |
| gPPI (w/ dec) | sPPI (w/ dec) | -0.181 | 0.087 | -2.075 | 1.000 | 0.057 | 0.284 | 1.604 |
| | | | | **Mean ISI = 8 s** | | | | |
| BSC-FRR | BSC-LSA | 23.827 | 0.087 | 272.755 | < .001 | 0.057 | ∞ | 3316.864 |
| | BSC-LSS | -2.908 | 0.087 | -33.283 | < .001 | 0.057 | ∞ | 826.693 |
| | gPPI (w/ dec) | -0.659 | 0.087 | -7.547 | < .001 | 0.057 | $1.187\times10^{+23}$ | 55.995 |
| | sPPI (w/ dec) | 0.709 | 0.087 | -8.112 | < .001 | 0.057 | $2.963\times10^{+27}$ | 66.120 |
| BSC-LSA | BSC-LSS | -26.735 | 0.087 | -306.038 | < .001 | 0.057 | ∞ | 3637.117 |
| | gPPI (w/ dec) | -24.487 | 0.087 | -280.302 | < .001 | 0.057 | ∞ | 3437.127 |
| | sPPI (w/ dec) | -24.536 | 0.087 | -280.867 | < .001 | 0.057 | ∞ | 3447.562 |
| BSC-LSS | gPPI (w/ dec) | 2.248 | 0.087 | 25.736 | < .001 | 0.057 | $2.922\times10^{+296}$ | 685.502 |
| | sPPI (w/ dec) | 2.199 | 0.087 | 25.171 | < .001 | 0.057 | $4.075\times10^{+292}$ | 676.624 |
| gPPI (w/ dec) | sPPI (w/ dec) | -0.049 | 0.087 | -0.565 | 1.000 | 0.057 | 0.004 | -2.553 |
| | | | | **Mean ISI = 12 s** | | | | |
| BSC-FRR | BSC-LSA | 4.492 | 0.087 | 51.403 | < .001 | 0.057 | ∞ | 1037.204 |
| | BSC-LSS | -3.602 | 0.087 | -41.219 | < .001 | 0.057 | ∞ | 1205.696 |
| | gPPI (w/ dec) | 0.398 | 0.087 | 4.557 | 0.002 | 0.057 | $3.651\times10^{+6}$ | 17.975 |
| | sPPI (w/ dec) | 0.211 | 0.087 | 2.411 | 1.000 | 0.057 | 1.384 | 3.189 |
| BSC-LSA | BSC-LSS | -8.093 | 0.087 | -92.645 | < .001 | 0.057 | ∞ | 2144.008 |
| | gPPI (w/ dec) | -4.093 | 0.087 | -46.858 | < .001 | 0.057 | ∞ | 945.145 |
| | sPPI (w/ dec) | -4.281 | 0.087 | -49.004 | < .001 | 0.057 | ∞ | 1020.727 |
| BSC-LSS | gPPI (w/ dec) | 4.000 | 0.087 | 45.787 | < .001 | 0.057 | ∞ | 1416.543 |
| | sPPI (w/ dec) | 3.812 | 0.087 | 43.641 | < .001 | 0.057 | ∞ | 1393.549 |
| gPPI (w/ dec) | sPPI (w/ dec) | -0.188 | 0.087 | -2.147 | 1.000 | 0.057 | 0.521 | 2.212 |

p values were adjusted for 50 t tests using Bonferroni correction. Bayesian comparisons are based on the default t test with a Cauchy (0, $r = 1/2^{0.5}$) prior. The posterior odds have been corrected for multiple testing by fixing the prior probability that the null hypothesis holds across all comparisons to 0.5. The "U" in the Bayes factor denotes that it is uncorrected. PPI terms were calculated with the deconvolution step (w/dec). sPPI and gPPI matrices were symmetrised.

#### Supplementary Table S7

Pairwise comparison of the sensitivity of TMFC methods for event related designs with different numbers of events per condition.

| Comparison | | Mean | SE | t | $p_{bonf}$ | Prior Odds | Posterior Odds | Log($BF_{10,U}$) |
|---|---|---|---|---|---|---|---|---|
| **Number of events = 20** | | | | | | | | |
| BSC-FRR | BSC-LSA | 3.845 | 0.118 | 32.493 | < .001 | 0.057 | ∞ | 809.599 |
| | BSC-LSS | -2.592 | 0.118 | -21.904 | < .001 | 0.057 | $6.353\times10^{+112}$ | 262.603 |
| | gPPI (w/ dec) | -0.752 | 0.118 | -6.352 | < .001 | 0.057 | $2.890\times10^{+11}$ | 29.254 |
| | sPPI (w/ dec) | -1.368 | 0.118 | -11.559 | < .001 | 0.057 | $6.672\times10^{+41}$ | 99.168 |
| BSC-LSA | BSC-LSS | -6.437 | 0.118 | -54.397 | < .001 | 0.057 | ∞ | 1276.956 |
| | gPPI (w/ dec) | -4.597 | 0.118 | -38.845 | < .001 | 0.057 | ∞ | 1063.888 |
| | sPPI (w/ dec) | -5.213 | 0.118 | -44.051 | < .001 | 0.057 | ∞ | 1225.250 |
| BSC-LSS | gPPI (w/ dec) | 1.840 | 0.118 | 15.552 | < .001 | 0.057 | $1.950\times10^{+61}$ | 143.990 |
| | sPPI (w/ dec) | 1.224 | 0.118 | 10.346 | < .001 | 0.057 | $4.238\times10^{+26}$ | 64.176 |
| gPPI (w/ dec) | sPPI (w/ dec) | -0.616 | 0.118 | -5.206 | < .001 | 0.057 | $1.647\times10^{+7}$ | 19.482 |
| **Number of events = 40** | | | | | | | | |
| BSC-FRR | BSC-LSA | 22.974 | 0.118 | 194.138 | < .001 | 0.057 | ∞ | 2647.594 |
| | BSC-LSS | -7.619 | 0.118 | -64.382 | < .001 | 0.057 | $2.688\times10^{+294}$ | 680.813 |
| | gPPI (w/ dec) | -6.653 | 0.118 | -56.217 | < .001 | 0.057 | $1.996\times10^{+269}$ | 622.951 |
| | sPPI (w/ dec) | -7.089 | 0.118 | -59.900 | < .001 | 0.057 | $8.267\times10^{+290}$ | 672.726 |
| BSC-LSA | BSC-LSS | -30.593 | 0.118 | -258.520 | < .001 | 0.057 | ∞ | 3204.757 |
| | gPPI (w/ dec) | -29.627 | 0.118 | -250.355 | < .001 | 0.057 | ∞ | 3370.030 |
| | sPPI (w/ dec) | -30.063 | 0.118 | -254.038 | < .001 | 0.057 | ∞ | 3362.369 |
| BSC-LSS | gPPI (w/ dec) | 0.966 | 0.118 | 8.165 | < .001 | 0.057 | 274141.661 | 15.386 |
| | sPPI (w/ dec) | 0.530 | 0.118 | 4.482 | 0.002 | 0.057 | 0.663 | 2.453 |
| gPPI (w/ dec) | sPPI (w/ dec) | -0.436 | 0.118 | -3.683 | 0.069 | 0.057 | 0.230 | 1.395 |
| **Number of events = 60** | | | | | | | | |
| BSC-FRR | BSC-LSA | 47.987 | 0.118 | 405.500 | < .001 | 0.057 | ∞ | 4134.22 |
| | BSC-LSS | -7.653 | 0.118 | -64.670 | < .001 | 0.057 | ∞ | 817.605 |
| | gPPI (w/ dec) | -4.785 | 0.118 | -40.434 | < .001 | 0.057 | $3.248\times10^{+186}$ | 432.323 |
| | sPPI (w/ dec) | -5.387 | 0.118 | -45.524 | < .001 | 0.057 | $1.069\times10^{+218}$ | 504.894 |
| BSC-LSA | BSC-LSS | -55.639 | 0.118 | -470.169 | < .001 | 0.057 | ∞ | 4454.717 |
| | gPPI (w/ dec) | -52.771 | 0.118 | -445.933 | < .001 | 0.057 | ∞ | 4468.663 |
| | sPPI (w/ dec) | -53.374 | 0.118 | -451.024 | < .001 | 0.057 | ∞ | 4434.657 |
| BSC-LSS | gPPI (w/ dec) | 2.868 | 0.118 | 24.236 | < .001 | 0.057 | $9.794\times10^{+76}$ | 180.143 |
| | sPPI (w/ dec) | 2.266 | 0.118 | 19.145 | < .001 | 0.057 | $4.023\times10^{+46}$ | 110.175 |
| gPPI (w/ dec) | sPPI (w/ dec) | -0.602 | 0.118 | -5.091 | < .001 | 0.057 | 31.672 | 6.320 |
| **Number of events = 80** | | | | | | | | |
| BSC-FRR | BSC-LSA | 48.182 | 0.118 | 407.149 | < .001 | 0.057 | ∞ | 4204.696 |
| | BSC-LSS | -8.349 | 0.118 | -70.548 | < .001 | 0.057 | ∞ | 1288.173 |
| | gPPI (w/ dec) | -4.282 | 0.118 | -36.182 | < .001 | 0.057 | $3.697\times10^{+221}$ | 513.043 |
| | sPPI (w/ dec) | -4.556 | 0.118 | -38.504 | < .001 | 0.057 | $3.109\times10^{+244}$ | 565.83 |
| BSC-LSA | BSC-LSS | -56.530 | 0.118 | -477.697 | < .001 | 0.057 | ∞ | 4772.541 |
| | gPPI (w/ dec) | -52.463 | 0.118 | -443.332 | < .001 | 0.057 | ∞ | 4599.074 |
| | sPPI (w/ dec) | -52.738 | 0.118 | -445.653 | < .001 | 0.057 | ∞ | 4611.084 |
| BSC-LSS | gPPI (w/ dec) | 4.067 | 0.118 | 34.365 | < .001 | 0.057 | $4.522\times10^{+266}$ | 616.861 |
| | sPPI (w/ dec) | 3.792 | 0.118 | 32.044 | < .001 | 0.057 | $1.235\times10^{+240}$ | 555.696 |
| gPPI (w/ dec) | sPPI (w/ dec) | -0.275 | 0.118 | -2.321 | 1.000 | 0.057 | 0.124 | 0.777 |
| **Number of events = 100** | | | | | | | | |
| BSC-FRR | BSC-LSA | 50.693 | 0.118 | 428.371 | < .001 | 0.057 | ∞ | 4675.406 |
| | BSC-LSS | -3.012 | 0.118 | -25.453 | < .001 | 0.057 | $9.388\times10^{+262}$ | 608.381 |
| | gPPI (w/ dec) | -1.846 | 0.118 | -15.603 | < .001 | 0.057 | $5.974\times10^{+118}$ | 276.357 |
| | sPPI (w/ dec) | -2.028 | 0.118 | -17.134 | < .001 | 0.057 | $1.850\times10^{+144}$ | 335.052 |
| BSC-LSA | BSC-LSS | -53.705 | 0.118 | -453.823 | < .001 | 0.057 | ∞ | 4917.987 |
| | gPPI (w/ dec) | -52.539 | 0.118 | -443.973 | < .001 | 0.057 | ∞ | 4883.486 |
| | sPPI (w/ dec) | -52.721 | 0.118 | -445.505 | < .001 | 0.057 | ∞ | 4907.266 |
| BSC-LSS | gPPI (w/ dec) | 1.166 | 0.118 | 9.850 | < .001 | 0.057 | $2.150\times10^{+68}$ | 160.206 |
| | sPPI (w/ dec) | 0.984 | 0.118 | 8.318 | < .001 | 0.057 | $3.209\times10^{+51}$ | 121.462 |
| gPPI (w/ dec) | sPPI (w/ dec) | -0.181 | 0.118 | -1.532 | 1.000 | 0.057 | 0.284 | 1.604 |

p values were adjusted for 50 t tests using Bonferroni correction. Bayesian comparisons are based on the default t test with a Cauchy (0, r = $1/2^{0.5}$) prior. The posterior odds have been corrected for multiple testing by fixing the prior probability that the null hypothesis holds across all comparisons to 0.5. The "U" in the Bayes factor denotes that it is uncorrected. PPI terms were calculated with the deconvolution step (w/dec). sPPI and gPPI matrices were symmetrised.

#### Supplementary Table S8

Pairwise comparison of the sensitivity of TMFC methods for event-related designs with different repetition times.

| **Comparison** | | **Mean** | **SE** | **t** | $p_{bonf}$ | **Prior Odds** | **Posterior Odds** | $\text{Log}(BF_{10,U})$ |
|---|---|---|---|---|---|---|---|---|
| | | | | **TR = 500 ms** | | | | |
| BSC-FRR | BSC-LSA | $1.200\times10^{-4}$ | 0.067 | 0.002 | 1.000 | 0.057 | 0.013 | -1.500 |
| | BSC-LSS | $2.558\times10^{-12}$ | 0.067 | $3.800\times10^{-11}$ | 1.000 | NaN | NaN | NaN |
| | gPPI (w/ dec) | $8.277\times10^{-12}$ | 0.067 | $1.230\times10^{-10}$ | 1.000 | NaN | NaN | NaN |
| | sPPI (w/ dec) | $1.240\times10^{-11}$ | 0.067 | $3.800\times10^{-10}$ | 1.000 | NaN | NaN | NaN |
| BSC-LSA | BSC-LSS | $-1.200\times10^{-4}$ | 0.067 | -0.002 | 1.000 | 0.057 | 0.013 | -1.500 |
| | gPPI (w/ dec) | $-1.200\times10^{-4}$ | 0.067 | -0.002 | 1.000 | 0.057 | 0.013 | -1.500 |
| | sPPI (w/ dec) | $-1.200\times10^{-4}$ | 0.067 | -0.002 | 1.000 | 0.057 | 0.013 | -1.500 |
| BSC-LSS | gPPI (w/ dec) | $5.719\times10^{-12}$ | 0.067 | $8.496\times10^{-11}$ | 1.000 | NaN | NaN | NaN |
| | sPPI (w/ dec) | $9.843\times10^{-12}$ | 0.067 | $1.462\times10^{-10}$ | 1.000 | NaN | NaN | NaN |
| gPPI (w/ dec) | sPPI (w/ dec) | $4.124\times10^{-12}$ | 0.067 | $6.126\times10^{-11}$ | 1.000 | NaN | NaN | NaN |
| | | | | **TR = 700 ms** | | | | |
| BSC-FRR | BSC-LSA | 0.018 | 0.067 | 0.262 | 1.000 | 0.057 | $5.802\times10^{+57}$ | 135.870 |
| | BSC-LSS | $3.537\times10^{-12}$ | 0.067 | $5.255\times10^{-11}$ | 1.000 | NaN | NaN | NaN |
| | gPPI (w/ dec) | $8.373\times10^{-12}$ | 0.067 | $1.244\times10^{-10}$ | 1.000 | NaN | NaN | NaN |
| | sPPI (w/ dec) | $5.723\times10^{-12}$ | 0.067 | $8.500\times10^{-11}$ | 1.000 | NaN | NaN | NaN |
| BSC-LSA | BSC-LSS | -0.018 | 0.067 | -0.262 | 1.000 | 0.057 | $5.802\times10^{+57}$ | 135.870 |
| | gPPI (w/ dec) | -0.018 | 0.067 | -0.262 | 1.000 | 0.057 | $5.802\times10^{+57}$ | 135.870 |
| | sPPI (w/ dec) | -0.018 | 0.067 | -0.262 | 1.000 | 0.057 | $5.802\times10^{+57}$ | 135.870 |
| BSC-LSS | gPPI (w/ dec) | $4.836\times10^{-12}$ | 0.067 | $7.183\times10^{-11}$ | 1.000 | NaN | NaN | NaN |
| | sPPI (w/ dec) | $2.185\times10^{-12}$ | 0.067 | $3.246\times10^{-11}$ | 1.000 | NaN | NaN | NaN |
| gPPI (w/ dec) | sPPI (w/ dec) | $-2.651\times10^{-12}$ | 0.067 | $-3.937\times10^{-11}$ | 1.000 | NaN | NaN | NaN |
| | | | | **TR = 1000 ms** | | | | |
| BSC-FRR | BSC-LSA | 1.919 | 0.067 | 28.507 | < .001 | 0.057 | ∞ | 1532.402 |
| | BSC-LSS | -0.002 | 0.067 | -0.031 | 1.000 | 0.057 | 50785.081 | 13.700 |
| | gPPI (w/ dec) | 0.005 | 0.067 | 0.075 | 1.000 | 0.057 | $7.830\times10^{+7}$ | 21.040 |
| | sPPI (w/ dec) | 0.006 | 0.067 | 0.090 | 1.000 | 0.057 | $2.448\times10^{+10}$ | 26.785 |
| BSC-LSA | BSC-LSS | -1.921 | 0.067 | -28.539 | < .001 | 0.057 | ∞ | 1534.286 |
| | gPPI (w/ dec) | -1.914 | 0.067 | -28.432 | < .001 | 0.057 | ∞ | 1527.832 |
| | sPPI (w/ dec) | -1.913 | 0.067 | -28.417 | < .001 | 0.057 | ∞ | 1526.842 |
| BSC-LSS | gPPI (w/ dec) | 0.007 | 0.067 | 0.106 | 1.000 | 0.057 | $9.450\times10^{+22}$ | 55.767 |
| | sPPI (w/ dec) | 0.008 | 0.067 | 0.122 | 1.000 | 0.057 | $1.074\times10^{+25}$ | 60.501 |
| gPPI (w/ dec) | sPPI (w/ dec) | 0.001 | 0.067 | 0.015 | 1.000 | 0.057 | 0.005 | -2.391 |
| | | | | **TR = 2000 ms** | | | | |
| BSC-FRR | BSC-LSA | 50.693 | 0.067 | 753.009 | < .001 | 0.057 | ∞ | 4675.406 |
| | BSC-LSS | -3.012 | 0.067 | -44.742 | < .001 | 0.057 | $9.388\times10^{+262}$ | 608.381 |
| | gPPI (w/ dec) | -1.846 | 0.067 | -27.427 | < .001 | 0.057 | $5.974\times10^{+118}$ | 276.357 |
| | sPPI (w/ dec) | -2.028 | 0.067 | -30.119 | < .001 | 0.057 | $1.850\times10^{+144}$ | 335.052 |
| BSC-LSA | BSC-LSS | -53.705 | 0.067 | -797.751 | < .001 | 0.057 | ∞ | 4917.987 |
| | gPPI (w/ dec) | -52.539 | 0.067 | -780.436 | < .001 | 0.057 | ∞ | 4883.486 |
| | sPPI (w/ dec) | -52.721 | 0.067 | -783.128 | < .001 | 0.057 | ∞ | 4907.266 |
| BSC-LSS | gPPI (w/ dec) | 1.166 | 0.067 | 17.315 | < .001 | 0.057 | $2.150\times10^{+68}$ | 160.206 |
| | sPPI (w/ dec) | 0.984 | 0.067 | 14.623 | < .001 | 0.057 | $3.209\times10^{+51}$ | 121.462 |
| gPPI (w/ dec) | sPPI (w/ dec) | -0.181 | 0.067 | -2.692 | 1.000 | 0.057 | 0.284 | 1.604 |
| | | | | **TR = 3000 ms** | | | | |
| BSC-FRR | BSC-LSA | 35.399 | 0.067 | 525.831 | < .001 | 0.057 | ∞ | 3969.703 |
| | BSC-LSS | -8.106 | 0.067 | -120.404 | < .001 | 0.057 | ∞ | 968.845 |
| | gPPI (w/ dec) | -18.961 | 0.067 | -281.653 | < .001 | 0.057 | ∞ | 2354.457 |
| | sPPI (w/ dec) | -19.332 | 0.067 | -287.163 | < .001 | 0.057 | ∞ | 2427.579 |
| BSC-LSA | BSC-LSS | -43.505 | 0.067 | -646.235 | < .001 | 0.057 | ∞ | 4370.359 |
| | gPPI (w/ dec) | -54.360 | 0.067 | -807.484 | < .001 | 0.057 | ∞ | 4916.715 |
| | sPPI (w/ dec) | -54.731 | 0.067 | -812.994 | < .001 | 0.057 | ∞ | 5003.281 |
| BSC-LSS | gPPI (w/ dec) | -10.855 | 0.067 | -161.249 | < .001 | 0.057 | ∞ | 1414.757 |
| | sPPI (w/ dec) | -11.226 | 0.067 | -166.759 | < .001 | 0.057 | ∞ | 1498.276 |
| gPPI (w/ dec) | sPPI (w/ dec) | -0.371 | 0.067 | -5.510 | < .001 | 0.057 | 0.167 | 1.073 |

p values were adjusted for 50 t tests using Bonferroni correction. Bayesian comparisons are based on the default t test with a Cauchy (0, r = 1/sqrt(2)) prior. The posterior odds have been corrected for multiple testing by fixing the prior probability that the null hypothesis holds across all comparisons to 0.5. The "U" in the Bayes factor denotes that it is uncorrected. PPI terms were calculated with the deconvolution step (w/dec). sPPI and gPPI matrices were symmetrised. NaN – data are essentially constant (sPPI, gPPI and BSC-LSS methods reached 100% sensitivity).

#### Supplementary Table S9

Parameters for the large-scale Wilson-Cowan neural mass model.

| Parameter | Description | Value |
|---|---|---|
| $\tau_E$ | Excitatory time constant | 2.50 ms |
| $\tau_I$ | Inhibitory time constant | 3.75 ms |
| $w_{EE}$ | Excitatory to excitatory synaptic weight | 16.0 |
| $w_{IE}$ | Inhibitory to excitatory synaptic weight | 12.0 |
| $w_{EI}$ | Excitatory to inhibitory synaptic weight | 15.0 |
| $w_{II}$ | Inhibitory to inhibitory synaptic weight | 3.0 |
| $a_E$ | Slope of excitatory response function | 1.5 |
| $a_I$ | Slope of inhibitory response function | 1.5 |
| $b_E$ | Position of maximum slope of excitatory response function | 3.0 |
| $b_I$ | Position of maximum slope of inhibitory response function | 3.0 |
| $c_E$ | Amplitude of excitatory response function | 1.0 |
| $c_I$ | Amplitude of inhibitory response function | 1.0 |
| $G$ | Global coupling parameter | 2.00 – 3.00 (2.63*) |
| $d$ | Signal transmission delay between brain regions | 25 ms |
| $P_E$ | Excitatory background drive | 0.700 – 0.800 (0.758*) |
| $P_I$ | Inhibitory background drive | 0 |
| $\tau_{ou}$ | Time scale of the Ornstein-Uhlenbeck process (background noise) | 5.0 ms |
| $\sigma_{ou}$ | Standard deviation of the Ornstein-Uhlenbeck process (background noise) | $(1.0 – 6.0)\times10^{-3}$ ($3.5\times10^{-3}$*) |
| $dt$ | Integration time step | 0.1 ms |
| $dt_{ds}$ | Downsampled time step | 5.0 ms |

[*] – The tuning parameters were chosen based on the maximum similarity between the ground-truth synaptic weight matrix and the task-modulated functional connectivity (TMFC) matrix estimated using the direct correlation difference (CorrDiff) approach for block design time series without scanner measurement noise.

#### Supplementary Table S10

Weighting factors for the construction of symmetrical and asymmetrical synaptic weight matrices.

**Symmetrical matrices**

| Condition | Rest | | | | Condition A | | | | Condition B | | | |
|---|---|---|---|---|---|---|---|---|---|---|---|---|
| ***Module*** | *№1* | *№2* | *№3* | *№4* | *№1* | *№2* | *№3* | *№4* | *№1* | *№2* | *№3* | *№4* |
| ***№1*** | 0.97 | 0.01 | 0.01 | 0.01 | 0.83 | 0.15 | 0.01 | 0.01 | 0.83 | 0.01 | 0.01 | 0.15 |
| ***№2*** | 0.01 | 0.97 | 0.01 | 0.01 | 0.15 | 0.83 | 0.01 | 0.01 | 0.01 | 0.83 | 0.15 | 0.01 |
| ***№3*** | 0.01 | 0.01 | 0.97 | 0.01 | 0.01 | 0.01 | 0.83 | 0.15 | 0.01 | 0.15 | 0.83 | 0.01 |
| ***№4*** | 0.01 | 0.01 | 0.01 | 0.97 | 0.01 | 0.01 | 0.15 | 0.83 | 0.15 | 0.01 | 0.01 | 0.83 |

**Asymmetrical matrices**

| Condition | Rest | | | | Condition A | | | | Condition B | | | |
|---|---|---|---|---|---|---|---|---|---|---|---|---|
| ***Module*** | *№1* | *№2* | *№3* | *№4* | *№1* | *№2* | *№3* | *№4* | *№1* | *№2* | *№3* | *№4* |
| ***№1*** | 0.97 | 0.01 | 0.01 | 0.01 | 0.83 | 0.15 | 0.01 | 0.01 | 0.83 | 0.01 | 0.01 | 0.15 |
| ***№2*** | 0.01 | 0.97 | 0.01 | 0.01 | 0.01 | 0.83 | 0.15 | 0.01 | 0.15 | 0.83 | 0.01 | 0.01 |
| ***№3*** | 0.01 | 0.01 | 0.97 | 0.01 | 0.01 | 0.01 | 0.83 | 0.15 | 0.01 | 0.15 | 0.83 | 0.01 |
| ***№4*** | 0.01 | 0.01 | 0.01 | 0.97 | 0.15 | 0.01 | 0.01 | 0.83 | 0.01 | 0.01 | 0.15 | 0.83 |

#### Supplementary Table S11

Fixed parameters for the Ballown-Windkessel haemodynamic model.

| Parameter | Description | Value |
|---|---|---|
| $\kappa$ | Rate of vasodilatory signal decay | 0.65 1/s |
| $\gamma$ | Rate of flow-dependent autoregulatory feedback | 0.41 1/s |
| $\tau$ | Haemodynamic transit time | 0.98 s |
| $\alpha$ | Grubb's exponent | 0.32 |
| $\rho$ | Resting net oxygen extraction fraction | 0.34 |
| $V_0$ | Resting volume fraction | 0.02 |

#### Supplementary Table S12

Variable parameters for the Ballown-Windkessel haemodynamic model.

| Parameter | Description | Mean value | SD |
|---|---|---|---|
| $\kappa$ | Rate of vasodilatory signal decay | 0.65 1/s | 0.0150 |
| $\gamma$ | Rate of flow-dependent autoregulatory feedback | 0.41 1/s | 0.0080 |
| $\tau$ | Haemodynamic transit time | 2.5 s | 0.2344 |
| $\alpha$ | Grubb's exponent | 0.32 | 0.0060 |
| $\rho$ | Resting net oxygen extraction fraction | 0.34 | 0.0096 |
| $V_0$ | Resting volume fraction | 0.02 | - |
